# Covalent adducts formed by the androgen receptor transactivation domain and small molecule drugs remain disordered

**DOI:** 10.1101/2024.11.12.623257

**Authors:** Jiaqi Zhu, Paul J. Robustelli

**Affiliations:** Department of Chemistry, Dartmouth College, Hanover, NH, USA

## Abstract

Intrinsically disordered proteins are implicated in many human diseases. Small molecules that target the disordered androgen receptor transactivation domain have entered human trials for the treatment of castration-resistant prostate cancer. These molecules have been shown to react with cysteine residues of the androgen receptor transactivation domain and form covalent adducts under physiological conditions. It is currently unclear how covalent attachment of these molecules alters the conformational ensemble of the androgen receptor. Here, we utilize all-atom molecular dynamics computer simulations to simulate covalent adducts of the small molecule ligands EPI-002 and EPI-7170 bound to the disordered androgen receptor transactivation domain. Our simulations reveal that the conformational ensembles of androgen receptor transactivation domain covalent adducts are heterogeneous and disordered. We find that covalent attachment of EPI-002 and EPI-7170 increases the population of collapsed helical transactivation domain conformations relative to the populations observed in non-covalent binding simulations and we identify networks of protein-ligand interactions that stabilize collapsed conformations in covalent adduct ensembles. We compare the populations of protein-ligand interactions observed in covalent adduct ensembles to those observed in non-covalent ligand-bound ensembles and find substantial differences. Our results provide atomically detailed descriptions of covalent adducts formed by small molecules and an intrinsically disordered protein and suggest strategies for developing more potent covalent inhibitors of intrinsically disordered proteins.

## Introduction

Intrinsically disordered proteins (IDPs) lack a rigid three dimensional (3D) structure under physiological conditions, and instead populate a dynamic conformational ensemble of rapidly interconverting structures ^1–4^. The structural plasticity of IDPs enables them to form complexes of varying affinities with multiple binding partners, which can exploited in cellular signalling and regulation pathways^5–7^. IDPs play important roles in a number of biological pathways, are implicated in various human diseases including neurodegenerative diseases, cancers and diabetes and represent a large pool of currently inaccessible drug targets ^8–14^.

A number of small molecules that directly bind IDPs and inhibit their interactions have been discovered ^15–25^ and several small molecules that bind IDPs have entered human trials^9,14,22^. Biophysical experiments have demonstrated that many IDPs remain disordered when bound to small molecule ligands^15–21,23,25–27^, spurring the development of new paradigms in molecular recognition ^11,12,18,27–31^. As biophysical measurements such as nuclear magnetic resonance (NMR) spectroscopy and small-angle x-ray scattering (SAXS) produce ensemble-averaged data that provide relatively sparse information about the conformational ensembles of IDP^4^, molecular dynamics (MD) computer simulations have become an essential tool for understanding the dynamic and heterogeneous interactions between IDPs and small molecule drugs and interpreting experimental data characterizing these binding events ^15,16,18,25,27–31^. MD simulation studies, together with validating biophysical experiments, suggest that the specificity and affinity of IDP ligands can be conferred through dynamic networks of transient interactions that only subtly shift the conformational ensemble of IDPs^15,16,18,27–30^.

The affinities of IDP-ligand binding interactions measured by conventional spectroscopic approaches thus far have been found to be relatively weak, with estimated K_D_ values in the *µ*M-mM range. ^15,16,18–21,25,26^. The spectroscopic signatures of IDP-ligand binding events, however, appear to vary considerably from the spectroscopic signatures of ligands binding to ordered binding sites of folded proteins and the interpretation of these measurements may not be straightforward. In several instances, small molecules that appear to bind IDPs with relatively weak millimolar affinities from residue-level NMR chemical shift perturbation measurements appear to bind with substantially tighter micromolar affinities from spectroscopic measurements from surface plasmon resonance (SPR) or biolayer interferometry^15,16,25^. Different spectroscopies used to measure ligand binding affinities are likely sensitive to different features of IDP-ligand binding modes and small molecule binding K_D_ estimates from spectroscopic measurements may therefore not always be directly comparable for IDPs and folded proteins.

Never-the-less, the affinities of IDP binders discovered thus far appear to be substantially weaker than the desired affinity of drugs that bind to structured binding pockets (K_D_ values in the pM-nM range). Several IDP ligands with weak *in vitro* affinities, however, have clear biological effects in cellular studies and animal models^16,22,24,25,27,32–34^. This suggests that lower affinity interactions may be sufficient to inhibit IDPs *in vivo* or that inhibition mechanisms may be more complex than reversible 1:1 stoichiometric inhibition, potentially involving interactions with high order molecular species ^19^ or biomolecular condensates ^27^. It is presently unclear how tightly small molecules with dynamic and heterogeneous noncovalent binding mechanisms can bind IDPs, and as many of the physiological interactions of IDPs are also relatively weak, it is also unclear how tightly small molecules must bind IDPs to exhibit biological activity and therapeutic effects in human trials.

The rational design of covalent drugs has gathered increasing interest in recent years ^35^. It is estiamted that roughly one-third of the FDA approved drugs act through covalent mechanisms ^36^. Covalent ligand discovery may be an attractive strategy for targeting IDPs^24,27,33,34,37^. The prolonged target engagement of covalent drugs can provide distinct pharmacodynamic profiles and increased potency, which may be especially important when trying to enhance the therapeutic effects of lower affinity IDP ligands. A series of covalently reactive compounds targeting the disordered N-terminal transactivation domain of the androgen receptor (AR) have shown promise for the treatment of castration-resistant prostate cancer (CRPC), as they inhibit constitutively active splice variants of AR that lack a ligand-binding domain and confer resistance to FDA-approved prostate cancer drugs^14,22,27,32,38–41^. The androgen receptor N-terminal transactivation domain (AR-NTD) inhibitor EPI-002, later named Ralaniten, was previously tested in clinical trials for CRPC but was discontinued after phase I due to excessive pill burden and poor metabolic properties ^14^. A second generation AR-NTD inhibitor, EPI-7170, was found to have improved potency and metabolic properties compared to EPI-002^39–41^. In March 2020 the compound EPI-7386, a third generation EPI AR-NTD inhibit later named Masofaniten, entered human trials as part of a combination treatment with the antiandrogen enzalutamide ^37^. This clinical trial was discontinued in October 2024 during phase 2 due to insufficient potency relative to treatment with only enzalutamide.

EPI-002 and EPI-7170 are both bisphenol-A derivatives that contain a chlorhydrin group (Figure 1A). The chlorohydrin group of EPI-002 was found to be weakly covalently reactive with cysteines in the androgen receptor N-terminal transactivation domain (AR-NTD), and this reactivity was found to be essential to its biological activity^22,32,38^. Bisphenol-A diglycidic ether (BADGE), an EPI-002 analog that contains a diol in place of a chlorohydrin group, was shown to have no biological activity^22,32^. It is currently hypothesized that covalent attachment to the AR-NTD is required for the biological activity of EPI compounds and other families of small molecule AR-NTD inhibitors^27,32–34^. Nuclear magnetic resonance (NMR) spectroscopy has been used to characterize the reversible non-covalent binding of EPI-002 to the AR-NTD^23^. NMR chemical shift perturbations (CSPs) localize the strongest interactions between EPI-002 and the AR-NTD to the transactivation unit 5 domain (Tau-5; AR residues A350-C448). The AR-NTD Tau-5 domain contains three regions with transiently populated helices (termed R1, R2 and R3) ^42,43^ and the R2 and R3 helices were found to have the largest NMR CSPs in EPI-002 binding titrations^23^.

**Figure 1:**
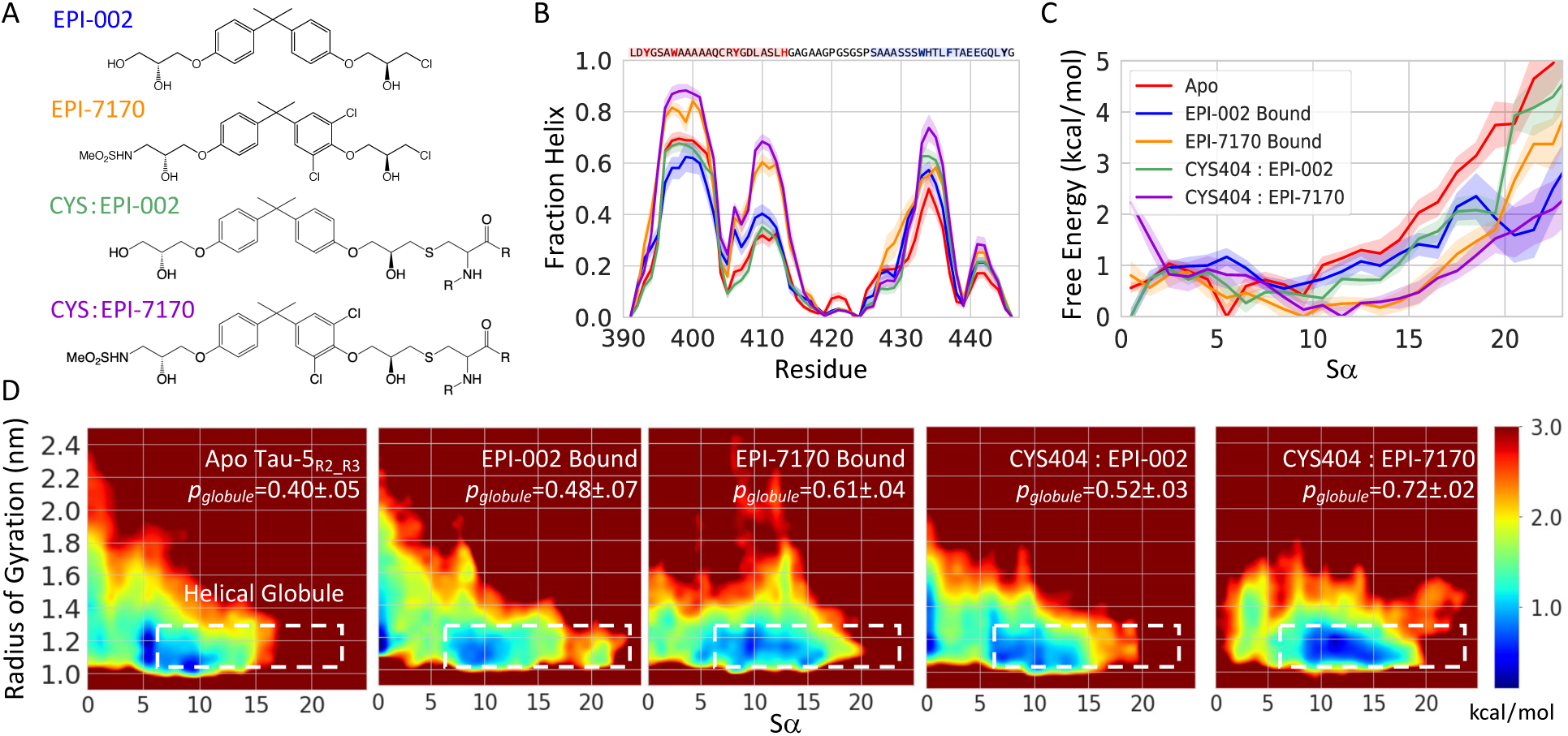
Covalent attachment of EPI-002 and EPI-7170 stabilizes collapsed helical molten-globule-like states of the Tau-5_R2 R3_ region of the androgen receptor transactivation domain. A) Chemical structures of EPI-002 and EPI-7170 and covalent adducts of EPI-002 and EPI-7170 attached to a cysteine residue. B) Helical propensities obtained from 300K replicas of REST2 MD simulations. Helical propensities are shown for apo Tau-5_R2 R3_ (red), the Tau-5_R2 R3_-CYS404:EPI-002 covalent adduct (green), the Tau-5_R2 R3_-CYS404:EPI-7170 covalent adduct (purple), a non-covalent ligand-bound ensemble of Tau-5_R2 R3_ and EPI-002 (blue) and a non-covalent ligand-bound ensemble of Tau-5_R2 R3_ and EPI-7170 (orange). Simulated helical propensities are presented as mean values ± statistical error estimates from blocking. C) Free energy of Tau-5_R2 R3_ conformations in each ensemble as a function of the helical collective variable *Sα*. D) Free energy surfaces as a function of the radius of gyration (*R_g_*) and *Sα*. The dotted white lines indicate the defined boundary of the “helical globule” state (*Sα >* 6.0, *R_g_ <* 1.3 nm). The population of the helical globule state in each ensemble is reported as *p_Glob_*.

Previously, we employed enhanced sampling all-atom molecular dynamics (MD) computer simulations to study the reversible non-covalent binding of EPI-002 and EPI-7170 to a 56-residue Tau-5 fragment (residues L391-G446) containing the R2 and R3 regions, which we refer to as Tau-5_R2 R3_ ^29^. This study revealed a heterogeneous ensemble of dynamic binding modes that localize the binding of EPI-002 and EPI-7170 to the interface between the R2 and R3 regions. We found that both compounds induced the formation of compact helical moltenglobule-like states, but that EPI-7170 had a 2.5 fold higher affinity to Tau-5_R2 R3_ and the Tau-5_R2 R3_:EPI-7170 bound ensemble was substantially more helical than the Tau-5_R2 R3_:EPI-002 bound ensemble. We identified a network of intermolecular interactions that confer higher affinity binding to EPI-7170, including stacking interactions between the dichlorinated phenyl ring of EPI-7170 and aromatic sidechains that form an interface between the R2 and R3 regions of Tau-5_R2 R3_. We observed that higher affinity non-covalent binding of EPI-7170 increased the proximity of the EPI-7170 chlorohydrin group to the reactive thiol of Tau-5_R2 R3_ residue cysteine 404 relative to the proximity of the EPI-002 chlorohydrin group to the cysteine 404 thiol group observed in EPI-002 binding simulations. These simulations support the previously proposed hypothesis that fast reversible non-covalent binding localizes reactive ligands to specific cysteines in the AR-NTD as the first step in AR inhibition^32^.

While there has been substantial progress characterizing non-covalent binding mechanisms of small molecules to IDPs^15,16,18,28–30^, relatively little is known about how covalent attachment of small molecule modifies the conformational ensembles of IDPs. In this study, we use all-atom explicit solvent enhanced sampling MD simulations with the state-of-the-art a99SB-*disp* force field^44^ to model covalent adducts of EPI-002 and EPI-7170 bound to the disordered AR-NTD. We report simulations of EPI-002 and EPI-7170 covalently attached to the thiol sulfur atom of residue cysteine 404 of the previously studied Tau-5_R2 R3_ AR construct ^29^.

We observe that covalent adducts of EPI-002 and EPI-7170 bound to Tau-5_R2 R3_ residue cysteine 404 remain heterogeneous and disordered, and that covalent attachment of these compounds does not induce Tau-5_R2 R3_ to fold into rigid structured conformations. We compare the conformational ensembles of covalent adducts of EPI-002 and EPI-7170 bound to Tau-5_R2 R3_ to the conformational ensembles of Tau-5_R2 R3_ observed in non-covalent binding simulations of EPI-002 and EPI-7170. We find that covalent attachment of these ligands increases the population of collapsed helical molten-globule-like Tau-5_R2 R3_ conformations relative to the populations observed in non-covalent binding simulations. We compare the populations of protein-ligand interactions observed in covalent adduct simulations to those observed in non-covalent binding simulations and find substantial differences in the populations of the most dominant interactions.

To obtain deeper insight into the effect of ligand binding and covalent ligand attachment on the conformational ensemble of the androgen receptor transactivation domain, we use a recently developed t-distributed stochastic neighbor embedding (t-SNE) clustering method^30^ to compare the conformational ensembles of Tau-5_R2 R3_ covalent adducts and non-covalent ligand-bound Tau-5_R2 R3_ ensembles. We identify several conformational states and binding modes that are present in both EPI-002 and EPI-7170 covalent adduct ensembles and characterize the structural properties and dominant protein:ligand interactions observed in these states. We find that there is substantially less overlap in the conformational space of Tau-5_R2 R3_ ensembles obtained from non-covalent ligand-binding simulations of EPI-002 and EPI-7170 compared to the overlap observed in covalent adduct ensembles, but still identify several conformational states and binding modes present in both non-covalent ligand-bound ensembles. Our results provide atomically detailed descriptions of covalent adducts formed by small molecules and an IDP, reveal differences in protein ligand interactions observed in IDP covalent adducts and non-covalent IDP-ligand bound ensembles and suggest possible strategies for developing more potent covalent IDP inhibitors.

## Results

We report unbiased all-atom explicit solvent MD simulations of covalent adducts of the small molecules EPI-002 and EPI-7170 bound to residue cysteine 404 of the previously studied 56-residue androgen transactivation domain fragment Tau-5_R2 R3_ (residues L391-G446) ^29^. We subsequently refer to these covalent adducts as “Tau-5_R2 R3_-CYS404:EPI-002” and “Tau-5_R2 R3_-CYS404:EPI-7170”. Chemical structures of the covalently modified cysteine sidechains, which we refer to as “CYS:EPI-002” and “CYS:EPI-7170”, are shown in Figure 1. Covalent adduct simulations were parameterized using the the a99SB-*disp* protein force field and a99SB-*disp* water model^44^ for canonical amino acids and water molecules. We generated parameters for the covalently modified CYS:EPI-002 and CYS:EPI-7170 sidechains using the generaized AMBER force field (GAFF1)^45^ (See “Parameterization of covalent cysteine adducts of EPI-002 and EPI-7170” in Methods). Simulations were run using the replica exchange with solute tempering (REST2) enhanced sampling algorithm ^46,47^ with 16 replicas spanning solute temperatures from 300-500K and all covalent adduct atoms selected for solute tempering (See “Molecular Dynamics Simulations” in Methods).

Simulations of Tau-5_R2 R3_-CYS404:EPI-002 were run for 4.8*µ*s/replica (aggregate simulation time of 77*µ*s) and simulations of Tau-5_R2 R3_-CYS404:EPI-7170 were run for 4.5*µ*s/replica (aggregate simulation time of 72*µ*s). Convergence of REST2 simulations was assessed by computing statistical error estimates by a blocking analysis^48,49^ and by comparing the secondary structure propensities and populations of intramolecular contacts observed in REST2 temperature rungs and independent demultiplexed replicas, which follow the continuous trajectories of simulated replicas through temperature space (SI Figures 1-8). The relatively smooth temperature dependence of these conformational properties among temperature replicas, and the size of the statistical deviations of the conformational properties of demulitplexed replicas suggest that the Tau-5_R2 R3_-CYS404:EPI-002 and Tau-5_R2 R3_-CYS404:EPI-7170 REST2 simulations are well converged. Further details are provided in the ”Statistical error estimates” section in Methods and the supporting information section “MD Simulation Convergence Analysis”.

Simulations of covalent adducts are compared to a previously reported REST2 simulation of apo Tau-5_R2 R3_ and previously reported REST2 non-covalent ligand-binding simulations of Tau-5_R2 R3_ in the presence of EPI-002 and EPI-7170^29^. Unless otherwise noted, all analyses in the main text pertain to the 300K replicas of each REST2 simulation. Conformational ensembles obtained from the 300K replicas of REST2 Tau-5_R2 R3_-CYS404:EPI-002 and Tau-5_R2 R3_-CYS404:EPI-7170 covalent adduct simulations are displayed in Supplementary Movie 1 and Supplementary Movie 2, respectively. The conformational ensembles of Tau-5_R2 R3_-CYS404:EPI-002 and Tau-5_R2 R3_-CYS404:EPI-7170 are heterogeneous and disordered, illustrating that covalent attachment of these ligands does not induce Tau-5_R2 R3_ to fold into rigid structured conformations. Conformational ensembles of Tau-5_R2 R3_-CYS404:EPI-002 and Tau-5_R2 R3_-CYS404:EPI-7170 have also been deposited in the protein ensemble database (PED)^50^ (accession codes pending).

### Covalent attachment of EPI-002 and EPI-7170 stabilizes collapsed helical molten-globule-like states of Tau-5_R2 R3_

We compare the helical propensities of the 300K REST2 ensembles of the Tau-5_R2 R3_-CYS404:EPI-002 and Tau-5_R2 R3_-CYS404:EPI-7170 covalent adducts with previously reported^29^ non-covalent ligand-bound ensembles of Tau-5_R2 R3_:EPI-002 and Tau-5_R2 R3_:EPI-7170 and an apo Tau-5_R2 R3_ ensemble in Figure 1B. We observe that the helical propensity of the Tau-5_R2 R3_-CYS404:EPI-002 ensemble is similar to the helical propensities of the noncovalent Tau-5_R2 R3_:EPI-002 bound ensemble and apo Tau-5_R2 R3_ ensemble. The average helical fraction of the Tau-5_R2 R3_-CYS404:EPI-002 ensemble (24.3 ± 0.7%), the full Tau-5_R2 R3_ ensemble (containing both bound and unbound frames) sampled in the non-covalent EPI-002 binding simulation (22.5 ± 1.7%), the bound frames of the non-covalent EPI-002 binding simulation (25.4 ± 1.3%), and the apo Tau-5_R2 R3_ ensemble (23.3 ± 0.6%) are largely within statistical error estimates. We observe a marginal increase in the helical fraction of Tau-5_R2 R3_-CYS404:EPI-7170 ensemble (34.3 ± 0.7%) relative to helical fraction of the non-covalent Tau-5_R2 R3_:EPI-7170 bound ensemble (32.8 ± 0.5%) and the full ensemble of Tau-5_R2 R3_ conformations sampled in the non-covalent EPI-7170 binding simulation (31.3 ± 0.8%). The similarity of these helical propensities demonstrates that Tau-5_R2 R3_ covalent adducts and non-covalent ligand-bound Tau-5_R2 R3_ ensembles have a similar degree of conformational disorder.

We previously observed that the reversible non-covalent binding of EPI-002 and EPI-7170 had a relatively small effect on the average helical fraction of Tau-5_R2 R3_ ensembles but stabilized the cooperative formation of multiple helical elements in collapsed states that have similar properties to molten-globule states observed in protein folding studies ^29,51^. We quantify the cooperative formation of helical elements using the *α*-helical order parameter *Sα*, which is a measure of the number of seven-residue fragments in a structure that resemble an ideal *α*-helix ^52^ (See Methods). The Tau-5_R2 R3_-CYS404:EPI-002 and Tau-5_R2 R3_-CYS404:EPI-7170 covalent adduct ensembles have average *Sα* values of 7.2 ± 0.3 and 11.0 ± 0.3, respectively. Tau-5_R2 R3_ ensembles containing all frames of non-covalent binding simulations of EPI-002 or EPI-7170 have average *Sα* values of 6.1 ± 0.4 and 9.1 ± 0.3, respectively, and Tau-5_R2 R3_ ensembles containing only bound frames of non-covalent binding simulations of EPI-002 or EPI-7170 have average *Sα* values of 7.3 ± 0.4 and 9.6 ± 0.2 respectively. The free energy surfaces of the covalent adduct ensembles and non-covalent ligand-bound ensembles are shown as a function of *Sα* in Figure 1C.

We compare the free energy surfaces of the covalent adduct ensembles, non-covalent ligand bound ensembles and the apo Tau-5_R2 R3_ ensemble as a function of the radius of gyration (*R_g_*) and *Sα* in Figure 1D. We note the apo Tau-5_R2 R3_ ensemble has a pronounced free energy minimum centered at (*Sα* = 5, *R_g_* = 1.2 nm). To quantify the relative populations of collapsed helical conformations, we utilize our previously proposed definition of Tau-5_R2 R3_ “helical globule” states, which is defined as Tau-5_R2 R3_ conformations with *Sα >* 6 and *R_g_ <* 1.3 nm ^29^. We report the population of helical globule conformations (*p_Glob_*) for each Tau-5_R2 R3_ ensemble in Figure 1D. The population of helical globule states in the apo Tau-5_R2 R3_ simulation is 40 ± 5%. The helical globule population of the EPI-002 covalent adduct ensemble (52 ± 3%) is increased relative to the apo Tau-5_R2 R3_ ensemble, the Tau-5_R2 R3_ ensemble obtained from the EPI-002 non-covalent binding simulation (38 ± 6%), and the Tau-5_R2 R3_ ensemble containing only bound frames of the EPI-002 non-covalent binding simulation (48 ± 7%). The helical globule population of the EPI-7170 covalent adduct ensemble (72 ± 2%) is substantially larger than the helical globule population of the noncovalent EPI-7170-bound ensemble (61 ± 4%) and the Tau-5_R2 R3_ ensemble containing all frames from the EPI-7170 non-covalent binding simulation (51 ± 5%)

### Identifying conformational substates of Tau-5_R2 R3_ ensembles

To obtain deeper insight into the effect of covalent ligand attachment and non-covalent ligand binding on the conformational ensemble of the androgen receptor transactivation domain, we employ a recently developed t-distributed stochastic neighbor embedding (t-SNE) clustering approach to compare the conformational ensembles of Tau-5_R2 R3_ covalent adducts and non-covalent bound ensembles^30^. The t-SNE clustering method is described in the Methods section “Clustering conformational ensembles of Tau-5_R2 R3_ with t-stochastic neighbor embedding (t-SNE)” and Eq. 2-7. Briefly, t-SNE takes a measure of the distance between between data points in a high-dimensional dataset as input and seeks to identify a low-dimensional projection where points that are nearby in the high-dimensional dataset have a high probability of being found in the same local neighborhood in the low-dimensional projection. Distances between points in the low-dimensional t-SNE embedding are calculated using a heavy-tailed Student’s t-distribution (Eq. 3). This ensures that points that are nearby in the high-dimensional dataset remain nearby in the low-dimensional projections, but allows dissimilar points to be modeled as further apart in low-dimensional projections. This is particularly useful for clustering IDP conformations, as structural metrics such as the root-mean-squared deviation (RMSD) of atomic positions are meaningful for describing smaller differences between IDP conformations with similar topologies, but are are relatively uninformative when comparing dissimilar IDP conformations with distinct topologies.

To identify Tau-5_R2 R3_ conformational states that are present in multiple conformational ensembles (ie. in both covalent adduct ensembles or both non-covalent ligand-bound ensembles) we first concatenate the ensembles we want to compare into a single *merged ensemble*. We then compute the pairwise RMSD of Tau-5_R2 R3_ C*α* atoms of all structures in the merged ensemble and utilize the resulting all-to-all RMSD matrix as input for dimensionality reduction with t-SNE. We perform t-SNE dimensionality reduction to obtain two-dimensional (2D) projections of our data with a range of values of the perplexity (*perp*) hyperparameter (Eq. 5). For each 2D projection, we subsequently perform *k* -means clustering of the data points using a range of values of the number of clusters (*N*). We evaluate the silhouette score (Eq. 7) of the cluster assignments obtained for each pair of perplexity and *N* values to identify optimal parameters for clustering at each desired level of resolution^30^ (SI Figure 9). We initially attempted to cluster Tau-5_R2 R3_ conformations from a merged ensemble containing all structures in the apo Tau-5_R2 R3_ ensemble, the Tau-5_R2 R3_-CYS404:EPI-002 covalent adduct ensemble, the Tau-5_R2 R3_-CYS404:EPI-7170 covalent adduct ensemble, and ensembles containing all frames of the non-covalent ligand-binding simulations of EPI-002 and EPI-7170. We found however, that clusters obtained from this merged ensemble had poor silhouette scores (data not shown). We observed that the apo Tau-5_R2 R3_ ensemble had little overlap with the covalent adduct ensembles or non-covalent ligand-bound ensembles in 2D t-SNE projections. We also observed that covalent adduct ensembles and non-covalent ligand-bound ensembles had relatively little overlap in 2D t-SNE projections when attempting to cluster a merged ensemble containing covalent adduct ensembles and ensembles from non-covalent binding simulations (data not shown).

We ultimately performed t-SNE clustering separately on i) a merged ensemble containing EPI-002 and EPI-7170 covalent adduct ensembles and ii) a merged ensemble containing all frames (bound and unbound) from EPI-002 and EPI-7170 non-covalent ligand-binding simulations (Figure 2, SI Figure 9). This produced clusters with higher silhouette scores, and identified Tau-5_R2 R3_ conformational substates with similar conformational properties and ligand binding modes in each of the individual conformational ensembles that were compared. We chose to analyze conformational states of covalent adducts and Tau-5_R2 R3_ ensembles from non-covalent ligand binding simulations at two levels of resolution. For each pair of ensembles we analyzed the cluster assignments that produced the highest silhouette scores when we restrict the number of clusters to *N* =4 (Figures 2-4, Tables 1-2, SI Figures 9-16). To obtain a higher resolution description of Tau-5_R2 R3_ conformational states, we also analyzed the cluster assignments that produced the highest silhouette scores for a larger numbers of clusters (10 ≤ *N* ≤ 20) (Figure 2, SI Figures 9-10, SI Figures 17-30, SI Tables 1-2). We display the t-SNE projections and average helical propensities of the Tau-5_R2 R3_ conformational states identified at both levels of clustering resolution in Figure 2. We display the average *β*-sheet propensities of each cluster in SI Figure 10. We provide visualizations of subsets of conformations from each of the clusters obtained with *N* =4 clusters for covalent adduct ensembles in Figure 3 and Tau-5_R2 R3_ ensembles obtained from non-covalent binding simulations in Figure 4.

**Figure 2:**
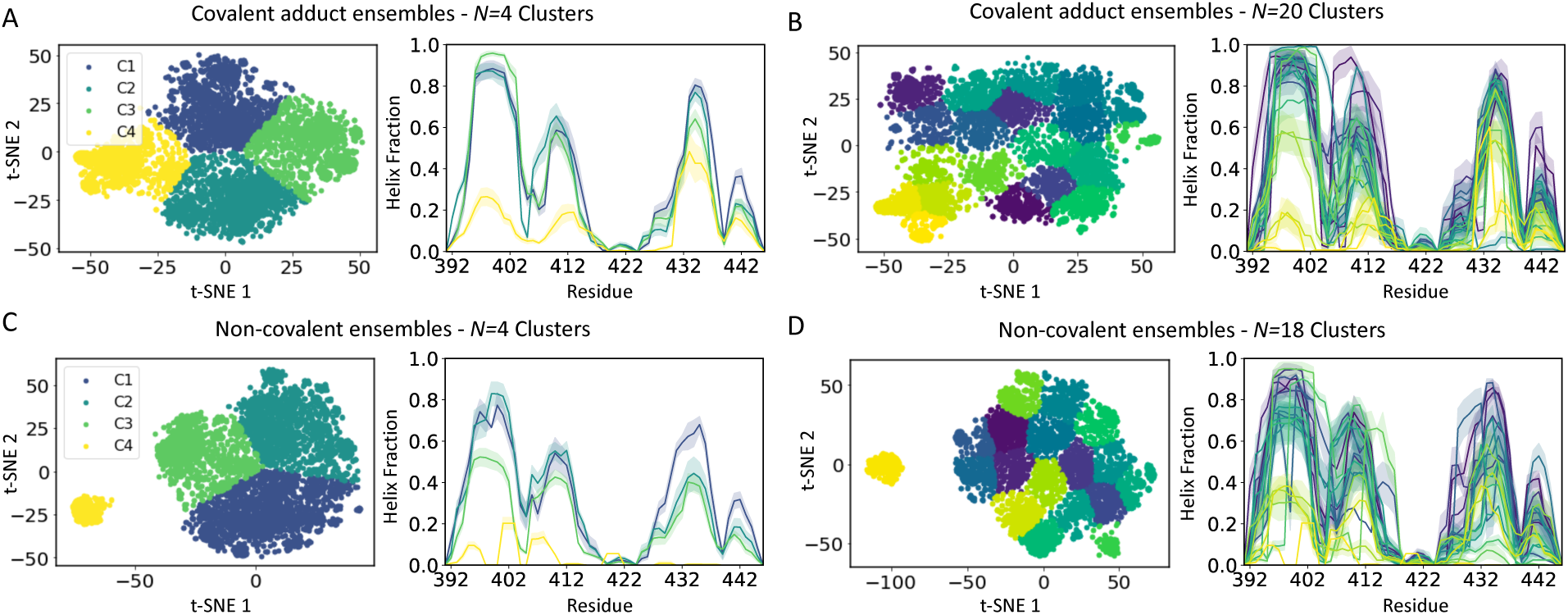
t-SNE clustering of Tau-5_R2 R3_ conformational states. t-SNE projections and cluster assignments of conformations from Tau-5_R2 R3_-CYS404:EPI-002 and Tau-5_R2 R3_-CYS404:EPI-7170 covalent adduct ensembles **(A, B)** and Tau-5_R2 R3_ conformations obtained from non-covalent EPI-002 and EPI-7170 ligand-binding simulations **(C, D)**. Cluster assignments by were obtained by performing t-SNE clustering on a merged ensemble containing the EPI-002 and EPI-7170 covalent adduct ensembles with *N* =4 and *N* =20 clusters and by performing t-SNE clustering on a merged ensemble containing all frames (bound and unbound) from EPI-002 and EPI-7170 non-covalent ligand-binding simulations with *N* =4 and *N* =18 clusters. t-SNE projections and average helical propensities of each t-SNE cluster are colored according to cluster assignments. Helical propensity are presented as mean values statistical error estimates from blocking.

**Figure 3:**
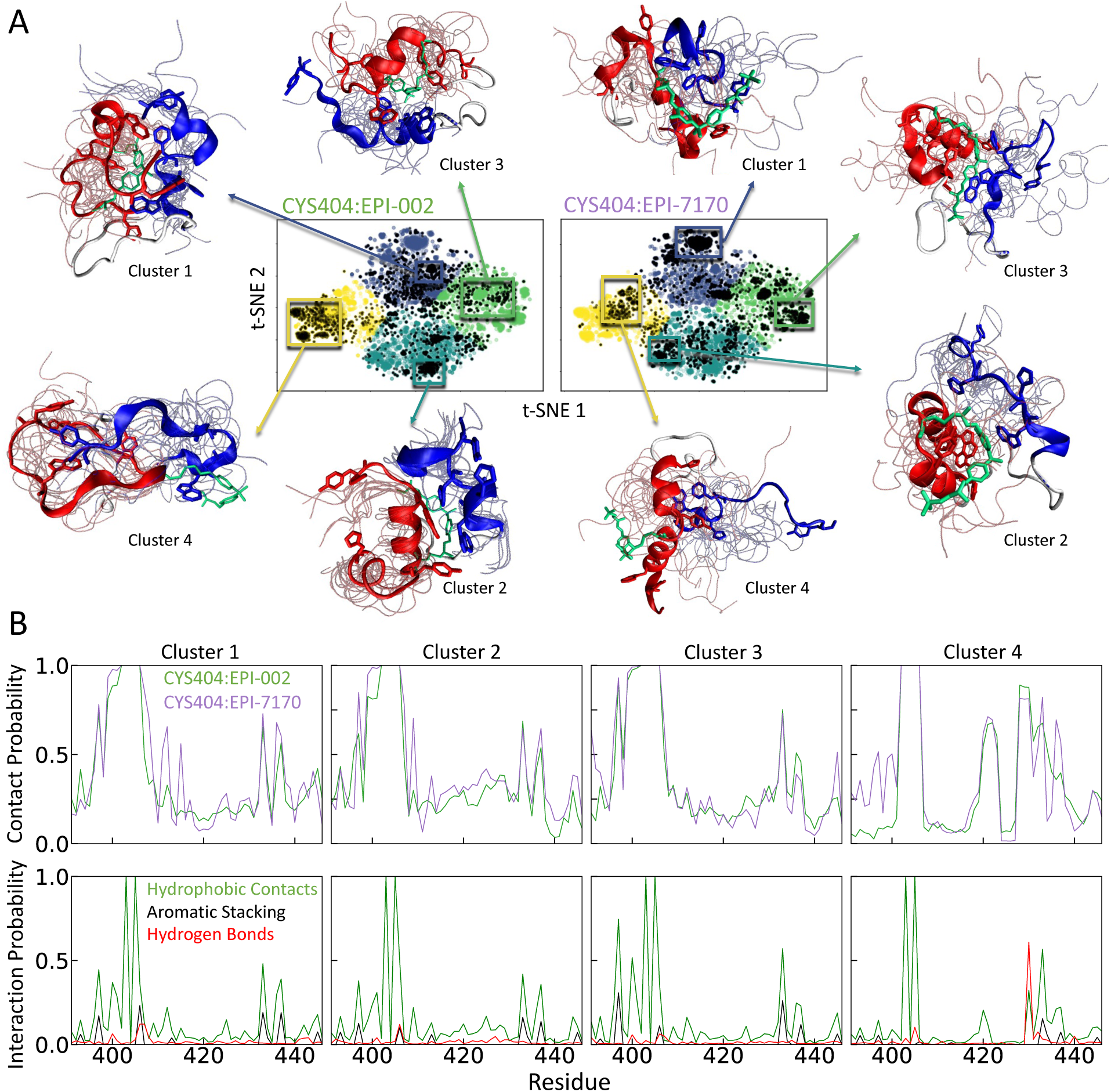
Protein-ligand interactions in Tau-5_R2 R3_ covalent adduct ensembles. **A)** t-SNE projections of covalent adduct ensembles of Tau-5_R2 R3_-CYS404:EPI-002 and Tau-5_R2 R3_-CYS404:EPI-7170 obtained with *N* =4 clusters. Colored dots correspond to conformations in the merged ensemble of both covalent adducts and black dots represent conformations from the specified individual covalent adduct ensemble. Illustrative snapshots of Tau-5_R2 R3_ are shown for selected subensembles of each cluster. A representative conformation of each subensemble is shown as a cartoon with the R2 and R3 regions of Tau-5_R2 R3_ colored red and blue, respectively, and the covalently attached ligand colored cyan. Backbone traces of additional conformations are shown as transparent tubes. **B)** Populations of intramolecular contacts and specific intramolecular interactions observed between covalently modified CYS404:EPI-002 and CYS404:EPI-7170 residues and Tau-5_R2 R3_ residues in each cluster.

**Figure 4:**
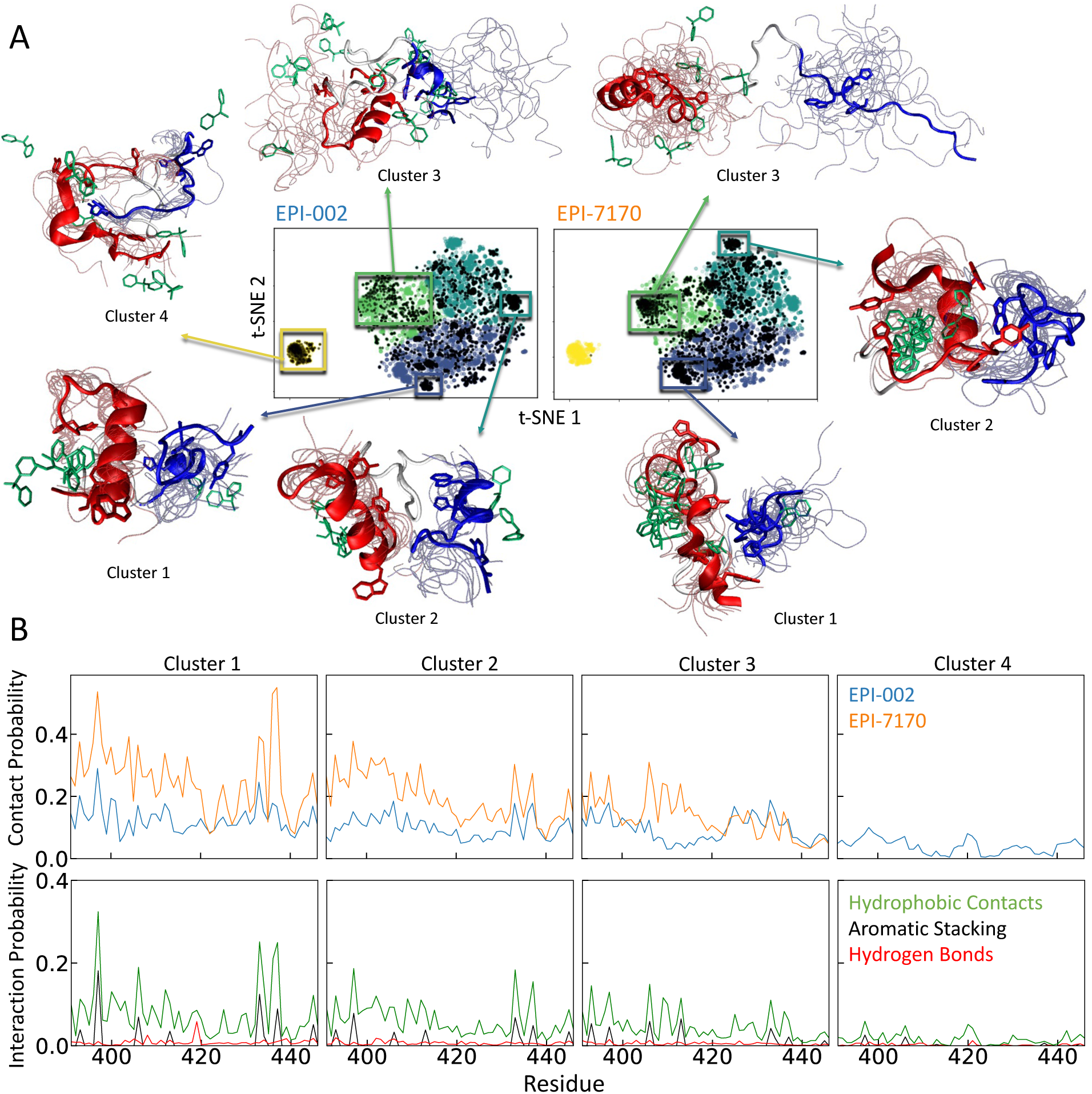
Protein-ligand interactions in Tau-5_R2 R3_ non-covalent ligand binding simulations of EPI-002 and EPI-7170. **A)** t-SNE projections of Tau-5_R2 R3_ conformations from non-covalent ligand binding simulations obtained with *N* =4 clusters. Colored dots correspond to conformations in a merged ensemble from both binding simulations and black dots represent conformations from the specified individual binding simulation. Illustrative snapshots of Tau-5_R2 R3_ are shown for selected subensembles of each cluster. A representative conformation of each subensemble is shown as a cartoon with the R2 and R3 regions of Tau-5_R2 R3_ colored red and blue, respectively. Backbone traces of additional conformations are shown as transparent tubes. The location of the bisphenol A scaffold of EPI ligands are shown for selected illustrative conformations in cyan. **B)** Populations of intermolecular contacts and specific intermolecular interactions observed between Tau-5_R2 R3_ and EPI-002 (blue) and Tau-5_R2 R3_ and EPI-7170 (orange) in each cluster.

**Table 1:** Cluster population (*p*), helical globule population (*pGlob*) and helix fraction (HF) of clusters obtained from t-SNE clustering of a merged ensemble of Tau-5R2 R3-CYS404:EPI-002 and Tau-5R2 R3-CYS404:EPI-7170 conformations with *N* =4 clusters. We compare the properties of the clusters in the merged ensemble to the properties of the clustered conformations from the the individual Tau-5R2 R3-CYS404:EPI-002 and Tau-5R2 R3-CYS404:EPI-7170 ensembles.

| Cluster | Merged Ensemble |  |  | Tau-5<sub>R2_R3</sub>-<br>CYS404:EPI-002 |  |  | Tau-5<sub>R2_R3</sub>-<br>CYS404:EPI-7170 |  |  |
| --- | --- | --- | --- | --- | --- | --- | --- | --- | --- |
| | $p$ | $p_{Glob}$ | HF | $p$ | $p_{Glob}$ | HF | $p$ | $p_{Glob}$ | HF |
| 1 | 0.27 | 0.76 | 0.34 | 0.27 | 0.72 | 0.30 | 0.27 | 0.80 | 0.38 |
| 2 | 0.29 | 0.72 | 0.32 | 0.25 | 0.56 | 0.25 | 0.32 | 0.84 | 0.37 |
| 3 | 0.25 | 0.81 | 0.31 | 0.24 | 0.80 | 0.30 | 0.25 | 0.82 | 0.32 |
| 4 | 0.20 | 0.09 | 0.12 | 0.24 | 0.02 | 0.08 | 0.15 | 0.21 | 0.18 |

**Table 2:** Cluster population (*p*), bound fraction (BF), helical globule population (*pGlob*) and helix fraction (HF) of clusters obtained from t-SNE clustering of a merged ensemble containing all frames from Tau-5R2 R3:EPI-002 and Tau-5R2 R3:EPI-7170 non-covalent binding simulations with *N* =4 clusters. We compare the properties of the clusters in the merged ensemble to the properties of the clustered conformations from the individual Tau-5R2 R3:EPI-002 and Tau-5R2 R3:EPI-7170 non-covalent binding simulations.

| Cluster | Merged Ensemble |  |  |  | Tau-5<sub>R2_R3</sub>:EPI-002 |  |  |  | Tau-5<sub>R2_R3</sub>:EPI-7170 |  |  |  |
| --- | --- | --- | --- | --- | --- | --- | --- | --- | --- | --- | --- | --- |
| | $p$ | BF | $p_{Glob}$ | HF | $p$ | BF | $p_{Glob}$ | HF | $p$ | BF | $p_{Glob}$ | HF |
| 1 | 0.39 | 0.62 | 0.69 | 0.31 | 0.37 | 0.49 | 0.67 | 0.29 | 0.41 | 0.74 | 0.70 | 0.33 |
| 2 | 0.35 | 0.57 | 0.64 | 0.28 | 0.33 | 0.41 | 0.54 | 0.25 | 0.37 | 0.70 | 0.72 | 0.31 |
| 3 | 0.19 | 0.52 | 0.15 | 0.20 | 0.16 | 0.48 | 0.11 | 0.14 | 0.22 | 0.54 | 0.18 | 0.23 |
| 4 | 0.06 | 0.30 | 0.00 | 0.03 | 0.13 | 0.29 | 0.00 | 0.03 | - | - | - | - |

We compare the populations of the Tau-5_R2 R3_ covalent adduct conformational states identified with t-SNE clustering with *N* =4 clusters alongside the average helical fraction and helical globule populations of each state in Table 1. We observe that three most helical clusters (clusters 1-3) have slightly larger populations in the Tau-5_R2 R3_-CYS404:EPI-7170 ensemble than in the Tau-5_R2 R3_-CYS404:EPI-002 ensemble. We observe cluster 4, which is the least helical and has a greater population of *β*-sheets (SI Figure 10), has a larger population in the Tau-5_R2 R3_-CYS404:EPI-002 ensemble. We find that within each cluster identified from the merged ensemble, conformations from the Tau-5_R2 R3_-CYS404:EPI-7170 ensemble have a larger helical fraction and substantially larger helical globule populations than the conformations from the Tau-5_R2 R3_-CYS404:EPI-002 ensemble. We compare the free energy surfaces of each cluster of the EPI-002 and EPI-7170 covalent adduct ensembles as a function of *R_g_* and *Sα* in SI Figure 12 and the populations of intramolecular contacts between Tau-5_R2 R3_ residues in SI Figure 13. These analyses reveal that the conformational properties of each cluster are fairly distinct and that the conformations from the EPI-7170 covalent adduct ensemble in each cluster have higher populations of collapsed helical states than the conformations from the EPI-002 covalent adduct ensemble. The same analyses are shown for Tau-5_R2 R3_ covalent adduct conformational states identified with t-SNE clustering with *N* =20 clusters in Table SI 1 ans SI Figures 20-23.

We compare the populations and helical content of Tau-5_R2 R3_ conformational states identified from non-covalent ligand binding simulations of EPI-002 and EPI-7170 with t-SNE clustering with *N* =4 clusters in Table 2. We observe that the conformations in clusters 1-3 from the non-covalent EPI-7170 binding simulation have larger helical fractions and substantially larger helical globule populations than the conformations in clusters 1-3 from the non-covalent EPI-002 binding simulation. We observe that cluster 4, which is highly collapsed and contains very little helical content, is only appreciably populated in the EPI-002 binding simulation. As only one conformation from the EPI-7170 non-covalent binding simulation was assigned to cluster 4, we omit ensemble analyses on this single conformation. We compare the free energy surfaces of each cluster identified from EPI-002 and EPI-7170 non-covalent binding simulations as a function of *R_g_* and *Sα* in SI Figure 15 and the populations of intramolecular contacts between Tau-5_R2 R3_ residues in SI Figure 16. The same analyses are shown for Tau-5_R2 R3_ conformational states identified from non-covalent binding simulations with t-SNE clustering with *N* =18 clusters in SI Table 2 and SI Figures 27-30. At both levels of resolution, we observe that binding EPI-7170 increases the populations of Tau-5_R2 R3_ conformational states with more compact helical conformations and also shifts the distribution of structures within each conformational state to contain compact helical conformations compared to binding EPI-002.

### Comparing protein-ligand interactions in Tau-5_R2 R3_ covalent adduct ensembles

For each Tau-5_R2 R3_ covalent adduct conformational state identified by t-SNE clustering, we compare the populations of the intramolecular interactions formed between the modified CYS404:EPI-002 and CYS404:EPI-7170 residues of the covalent adducts with each Tau-5_R2 R3_ residue. We compare the populations of intramolecular interactions formed by CYS404:EPI-002 and CYS404:EPI-7170 in the conformational states identified with t-SNE clustering with *N* =4 clusters in Figure 3B and in the conformational states identified with *N* =20 clusters in SI Figures 17-19. We define intramolecular contacts as occurring with a Tau-5_R2 R3_ residue in any frame where at least one heavy (non-hydrogen) atom of that residue is within 6.0 ^°^A of a heavy atom of the modified CYS404 residue. We note that by this definition CYS404:EPI-002 and CYS404:EPI-7170 posses contacts with the neighboring Tau-5_R2 R3_ residues ^402^AQCRY^406^ in all frames (Figure 3B). We calculate the populations of specific interactions (hydrophobic contacts, aromatic stacking interactions, and hydrogen bonding interactions) between CYS404:EPI-002 and CYS404:EPI-7170 and each residue of Tau-5_R2 R3_ in each cluster as specified in the “Protein-Ligand Interactions” section in Methods.

The populations of intramolecular contacts formed by CYS404:EPI-002 and CYS404:EPI-7170 are remarkably similar in the conformational states identified with *N* =4 clusters (Figure 3B). The values of the coefficient of determination (*r*^2^) between the populations of intramolecular contacts formed by CYS404:EPI-002 and CYS404:EPI-7170 observed in clusters 1-4 are 0.83, 0.89, 0.86 and 0.77, respectively. This demonstrates that the interactions of the covalently modified CYS404 sidechains are extremely similar in the clustered states of both covalent adduct ensembles even though cluster assignments were obtained considering only the positions of backbone C*α* atoms. We display the populations of hydrophobic contacts, aromatic stacking interactions and hydrogen bonding interactions between modified CYS404 residues and each Tau-5_R2 R3_ residue in each cluster of the merged covalent adduct ensemble in Figure 3B and compare the populations of interactions in the clustered conformations of each individual covalent adduct ensemble in SI Figure 11.

The populations of the intramolecular interactions formed by CYS404:EPI-002 and CYS4 04:EPI-7170 are relatively similar in clusters 1-3 (Figure 3, SI Figure 11) despite the fact that we observe substantial differences in the structural properties of these states (SI Figures 12-13). This demonstrates that similar protein-ligand interactions can be formed in conformational states of Tau-5_R2 R3_ with distinct structural properties. Clusters 1-3 contain highly populated contacts between CYS404:EPI-002 and CYS404:EPI-7170 with neighboring residues in the R2 region of Tau-5_R2 R3_ (residues ^396^AWAAAAAQCRY^406^) as well as substantially populated contacts with residues in the R3 region (^430^SSSWHTLFTAE^440^), which includes the partially helical ^432^SWHTLF^437^ molecular recognition motif. In these clusters, the covalently modified CYS404 sidechains make hydrophobic and aromatic stacking interactions with residues from both the R2 and R3 regions to form dynamic and heterogeneous aromatic cores, stablizing the the formation of compact helical states (Table 1). The dominant interactions of the CYS404:EPI-002 and CYS404:EPI-7170 sidechains are with the aromatic residues Y393, W397, Y406, W433 and F437, and each cluster is differentiated by the relative populations of these interactions.

Cluster 4, which has a larger population in the Tau-5_R2 R3_-CYS404:EPI-002 ensemble, has substantially different conformational properties than clusters 1-3 in both covalent adduct ensembles. Cluster 4 has substantially lower populations of helical conformations and collapsed helical globule states and higher populations of *β*-sheets relative to clusters 1-3 (Figure 2, SI Figure 10). The *β*-sheets formed in cluster 4 generally do not contain stretches of contiguous residues, and frequently consist of only pairs of residues. In cluster 4, CYS404:EPI-002 and CYS404:EPI-7170 form highly populated contacts with residues ^418^AGPGS^422^ and residues ^426^SAAASSSWHTLF^437^. These interactions frequently include a highly populated hydrogen bond between the backbone carbonyl oxygens of CYS404:EPI-002 or CYS404:EPI-7170 and the backbone amide of SER430 as well as additional hydrogen bonds formed by the SER430 sidechain hydroxyl group and diols or methyl sulfonamide oxygens in CYS404:EPI-002 and CYS404:EPI-7170. We observe hydrogen bonds between SER430 and CYS404:EPI-002 or CYS404:EPI-7170 in over sixty percent of the conformations in cluster 4. We also observe substantial hydrophobic and aromatic stacking interactions with residues W433, L436 and F437 (Figure 3, SI Figure 11).

Analysis of the conformational properties (SI Table 1, SI Figures 20-23) and intramolecular interactions of the CYS404:EPI-002 and CYS404:EPI-7170 residues (SI Figures 17-19) of the covalent adduct conformational states identified by t-SNE clustering with *N* =20 clusters reveals that the four clusters discussed above can be effectively split into more homogeneous conformational states. At this finer level of resolution we find larger differences in the populations of intramolecular protein-ligand interactions formed by CYS404:EPI-002 and CYS404:EPI-7170 in several clusters. The average 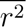 value (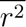) of the populations of intramolecular contacts formed by CYS404:EPI-002 and CYS404:EPI-7170 decreases from 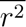 = 0.84 for *N* =4 clusters to 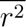 = 0.66 for *N* =20 clusters (SI Figure 17). We observe that the interactions of the covalently modified CYS404 residues are similar in many clusters; nine of the twenty clusters have 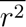 values greater than 0.75. SI Figures 18-19 reveal that identities of the hydrophobic contacts, aromatic stacking interactions and hydrogen bonding interactions formed by CYS404:EPI-002 and CYS404:EPI-7170 are very similar in many clusters, but their populations can substantially vary. We note that several of the conformational states identified by t-SNE clustering with *N* =20 clusters predominantly contain specific intramolecular interactions between covalently modified CYS404 residues and residues in either the R2 region or residues in the R3 region of Tau-5_R2 R3_ (SI Figures 17-19). This demonstrates that not all interactions made by the covalently modified CYS404 sidechains involve simultaneous interactions with both regions.

### Comparing protein-ligand interactions in non-covalent ligand-bound Tau-5_R2 R3_ ensembles

We compare the populations of intermolecular protein-ligand interactions formed between each residue of Tau-5_R2 R3_ and EPI-002 or EPI-7170 in each conformational state identified by t-SNE clustering of ensembles from non-covalent ligand binding simulations in Figure 4. We display the populations of intermolecular interactions formed by EPI-002 and EPI-7170 in conformational states identified with t-SNE clustering with *N* =4 clusters in Figure 4 and in conformational states identified with *N* =18 clusters in SI Figures 24-26. There are larger differences in the populations of protein-ligand interactions of EPI-002 and EPI-7170 in clusters obtained from non-covalent binding simulations compared to the differences observed in clusters obtained from covalent adduct simulations. The average 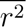 value of the populations of intermolecular contacts formed by EPI-002 and EPI-7170 in each cluster is 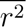 = 0.36 for *N* =4 clusters to 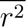 = 0.17 for *N* =18 clusters (SI Figure 24). This indicates that the protein-ligand binding modes observed in non-covalent binding simulations are more heterogeneous and have substantially greater variations between ligands. This is somewhat unsurprising given the relatively restricted set of orientations accessible to covalently bound ligands in covalent adduct ensembles relative to free ligands.

As we cluster both bound and unbound frames of ensembles obtained from non-covalent ligand binding simulations we can compare the fraction of bound frames observed in each cluster, where bound frames are defined as containing at least one pair of ligand and protein heavy atoms within 6.0 ^°^A. We compare the fraction of frames bound to EPI-002 or EPI-7170 for *N* =4 clusters in Table 2 and for *N* =18 clusters in SI Table 2. For clusters obtained by t-SNE clustering with *N* =4, we observe that fraction of bound frames in clusters 1-3 is substantially higher in the EPI-7170 binding simulation than the EPI-002 binding simulation (we omit a comparison of cluster 4, which contains only one frame from the EPI-7170 binding simulation). For clusters obtained by t-SNE clustering with *N* =18, we observe that the fraction of bound frames is substantially higher in the EPI-7170 binding simulation in fifteen of the seventeen clusters populated in both binding simulations. These results demonstrate that the higher affinity binding of EPI-7170 does not result from stabilizing binding-competent Tau-5_R2 R3_ conformational states that are not sampled in EPI-002 binding simulations - but instead is the result of EPI-7170 having a substantially higher affinity to a large diversity of conformations sampled in both ligand-binding simulations.

We compare the populations of intermolecular hydrophobic contacts, aromatic stacking interactions and hydrogen bonding interactions formed by EPI-002 and EPI-7170 in clusters obtained from the non-covalent ligand-binding simulation ensembles with *N* =4 clusters in Figure 4B and compare the populations of these interactions in each individual non-covalent ligand-binding simulations in SI Figure 14. The same analyses are shown for the Tau-5_R2 R3_ conformational states identified from non-covalent ligand-binding simulations by t-SNE clustering with *N* =18 clusters in SI Figures 25-26. We observe substantially lower populations of nearly all specific intermolecular protein-ligand interactions in EPI-002 binding simulations compared to EPI-7170 simulations in all identified clusters. This is consistent with previous results demonstrating that the increased aromatic stacking propensity of the chlorinated phenyl ring of EPI-7170 more effectively localizes this ligand into dynamic hydrophobic cores of collapsed Tau-5_R2 R3_ states where it can form dynamic networks of interconverting intermolecular interactions ^29^.

## Discussion

The development of small molecule inhibitors targeting the disordered N-terminal transactivation domain of the androgen receptor (AR-NTD) is a potential approach for treating castration-resistant prostate cancer (CRPC)^9,14^. Several small molecules that bind to the disordered Tau-5 domain of the AR-NTD and inhibit the transcriptional activity of AR have been discovered, and a number studies suggest that covalent attachment of these molecules to the AR-NTD is required for their biological activity^27,32–34^. Biophysical experiments^23,27^ and computer simulations^29^ have been used to characterize the non-covalent binding mechanisms of small molecules to the AR-NTD and other IDPs^15,16,18,25,28,31^. Despite the biological importance of the covalent reactivity of AR-NTD inhibitors, to our knowledge, prior to this investigation the effect of covalent attachment of small molecules on the conformational ensemble of the AR-NTD has not been characterized at a molecular level.

In this investigation we report atomic resolution conformational ensembles of covalent adducts formed by the small molecules and a disordered region of the androgen receptor transactivation domain obtained from MD simulations with a state-of-the-art force field^44^ and enhanced sampling technique^46,47^. We performed MD simulations of a covalent adduct formed by the intrisically disordered AR Tau-5_R2 R3_ construct and EPI-002, a compound that was previously tested in phase I clincal trials for CRPC under the name Ralaniten, and a covalent adduct formed by Tau-5_R2 R3_ and EPI-7170, a second generation bisphenol-A scaffold EPI inhibitor that showed improved potency and in cellular assays and animal CRPC models^39–41^. While several additional AR-NTD inhibitors have been discovered ^27,32–34^ who chose to focus on covalent adducts formed by EPI-002 and EPI-7170 to enable direct comparisons to our previous work studying the non-covalent binding mechanisms of these compounds to AR-NTD^29^.

We observed that the conformational ensembles of covalent adducts formed by Tau-5_R2 R3_ with EPI-002 and EPI-7170 contain a similar degree of conformational heterogeneity and disorder as conformational ensembles of apo Tau-5_R2 R3_ and ensembles of Tau-5_R2 R3_ noncovalently bound to EPI-002 and EPI-7170. Our simulations do not suggest that covalent attachment of these molecules stabilizes the formation of more rigid folded states of Tau-5_R2 R3_, or that covalent attachment substantially restricts the conformational space accessible to Tau-5_R2 R3_. We do find, however, that covalent attachment of EPI-002 and EPI-7170 drives the Tau-5_R2 R3_ ensemble to populate more compact helical molten-globule like states relative to the apo ensemble and non-covalent ligand-bound ensembles.

To obtain deeper insight into how non-covalent ligand-binding and covalent adduction of EPI-002 and EPI-7170 modifies the conformational ensemble of Tau-5_R2 R3_, we employed a recently developed t-SNE based clustering method^30^ to identify conformational states that are populated in multiple Tau-5_R2 R3_ ensembles and quantified how the populations and properties of these states change in the presence of each ligand. In both non-covalent binding simulations and covalent adduct simulations, we find that EPI-7170 increases the populations of states with higher helical propensities and larger populations of helical globule states relative to EPI-002. We also observe that the identity of the ligand shifts the conformational properties within each clustered conformational state. Tau-5_R2 R3_ conformations within each cluster, which have similar C*α* coordinates, generally have higher helical fractions and helical globule populations in the presence of EPI-7170 compared to EPI-002. The identity of the ligand therefore both affects the populations of conformational states, in a manner analogous to the concept of conformational selection, and the conformational properties of each conformational state, in a manner analogous to concept of induced fit, in both covalent adduct ensembles and non-covalent bound ensembles.

We note that attempting to simultaneously cluster conformations from apo, covalent adduct and non-covalent ligand-bound Tau-5_R2 R3_ ensembles produced substantially worse clustering results and less homogeneous clusters compared to clustering conformations from only covalent adduct ensembles or from only non-covalent ligand-binding simulations. This suggests that while the average properties of these ensembles, like the fraction helix or radius of gyration, are not dramatically different, there are still substantial shifts in the distribution of conformational states accessible to Tau-5_R2 R3_ in its apo state and ligand-bound states. Covalent adduction and non-covalent ligand binding appear to have an effect on the conformational ensemble of Tau-5_R2 R3_ that more closely resembles an induced-fit binding mechanism than conformational selection. This makes sense considering the binding modes we observe in this study and previous work. ^29^ EPI-002 and EPI-7170 frequently interact and intercalate with aromatic Tau-5_R2 R3_ residues that form a dynamic hydrophobic core. In the absence of these ligands, we expect that the aromatic residues of Tau-5_R2 R3_ will tightly pack with one another, and not leave large void volumes that could be filled by a ligand in a conformational selection mechanism.

By comparing conformational substates of covalent adduct ensembles of Tau-5_R2 R3_-CYS404:EPI-002 and Tau-5_R2 R3_-CYS404:EPI-7170 with t-SNE clustering, we identify several extremely similar conformational states, with very similar protein-ligand interactions in both ensembles. While this is not entirely unexpected given the similarity of EPI-002 and EPI-7170, it does highlight the possibility of utilizing atomic resolution conformational ensembles of covalent adducts from MD simulations as a basis for rational structure-based design of novel ligands that shift the conformational ensembles of IDP covalent adducts in a desired fashion. For example, if one sought to design covalent ligands that further stabilized collapsed helical-globule states relative to the ligands studied here, one could attempt to use structures from the Tau-5_R2 R3_-CYS404:EPI-002 or Tau-5_R2 R3_-CYS404:EPI-7170 ensembles to identify regions where ligand substitutions or fragment decoration are predicted to increase the stability of collapsed conformations.

If the covalent adduct ensembles of relatively similar ligands like EPI-002 and EPI-7170 were extremely different, and had very little conformational overlap - one would not expect to be able to use the populations of protein-ligand interactions, or conformational ensembles of specific ligand interaction modes, to inform the design of novel ligands that stabilize specific substates of a conformational ensemble. If many of the conformational substates and protein-ligand interactions are highly similar in covalent adduct ensembles, an MD ensemble may provide a useful starting point for structure based design of modified ligands. We observe much larger differences in the ligand-binding modes observed in conformational substates identified in non-covalent ligand-binding simulations, suggesting that it may be more challenging to utilize non-covalent ligand-bound ensembles to predict perturbations in IDP ensembles that might be obtained by modifying compounds based on simulated ligandbinding poses from MD simulations.

It has previously been hypothesized that fast reversible non-covalent binding may preferentially localize ligands to specific cysteines in the AR-NTD as a first step in an AR transcriptional inhibition^14,32^. The R3 domain of the AR-NTD folds-upon-binding the RAP74 domain of the general transcription regulator TFIIF and the disruption of this interaction causes AR to lose its transciptional activity^42,43,53,54^. One potential AR inhibition mechanism supported by our simulations is that fast reversible non-covalent binding may localize ligands to specific cysteine residues and induce the formation collapsed Tau-5 helical states that sequester the reactive ligand moieties and cysteine thiol groups from solvent, accelerating rates of attachment to specific cysteines. Once attached, protein-ligand interactions, predominantly driven by the aromatic stacking and hydrophobic interactions observed in this work, may sequester the R3 region of Tau-5 into compact molten-globule like states that are incompatible with TFIIF binding.

Based on the conformational ensembles determined in this study, we speculate that that covalent attachment of EPI ligands to the AR-NTD could also influence the activity in AR in other ways beyond direct inhibition of Tau-5:TFIIF binding interactions. For example, covalent attachment of ligands to the AR-NTD could increase the formation of clusters of aromatic residues sequestered from solvent, potentially reducing rates of nuclear translocation by competing with interactions with nuclear pore proteins.

Recent work from several laboratories has demonstrated that the transcriptional activity of AR is linked to the formation of biomolecular condenstates in cells and that small molecule AR inhibitors can affect the propensity of AR to form condensates ^27,33,34,55^. EPI-002 and other ligands have been shown to partition into condensates ^27^, and NMR studies have identified the R2 and R3 regions of the Tau-5 region of the AR-NTD are essential for driving the formation of higher order AR states that proceed phase separation in-vitro ^56^. Covalent adduct formation my induce the formation of collapsed helical conformations in AR-NTD that have a higher propensity to oligomerize and form condensates. It is also possible that once a covalent adduct is formed, covalently attached ligands could form intermolecular interactions with other AR-NTD molecules facilitating the formation of higher order species that accelerate the formation of biomolecular condensates or modulate the physical or cellular properties of condensates, such as condensate stability or condensate stiffness, which may affect AR transcriptional activity levels in cells. One can also envision a positive feedback loop where covalent attachment of small molecules to AR stabilizes the formation of biomolecular condensates and additional ligands partition into the condensates, further accelerating the rate of covalent attachment.

This study provides atomic-resolution structural ensembles of covalent adducts formed small molecule drugs and an IDP, insight into potential inhibition mechanisms of AR inhibitors and insight into how covalent attachment of small molecules can influence the conformational ensembles IDPs relative to non-covalent ligand binding. The atomic resolution conformational ensembles described here may provide a useful model system for attempting to develop ensemble-based approaches to design ligands or protein mutations that rationally perturb the conformational ensembles of IDP covalent adducts. The cysteine adduct force field parametrization strategy and enhanced sampling strategy presented in this work provides a template for performing future studies of covalent adducts formed by IDPs and small molecules, and could be used to examine how covalent attachment of other recently discovered AR inhibitors^27,33,34^ affect the conformational ensemble of the AR-NTD.

As most IDP-ligand interactions discovered thus far have relatively weak binding affinities, developing covalent ligands may be an appealing strategy for discovering IDP therapeutics for currently untreatable diseases. We believe that molecular simulations of covalent IDP adducts will play a valuable role in understanding how covalent attachment of ligands modulate the conformational ensembles of IDPs. MD simulations of IDP covalent adducts could facilitate the selection of optimal covalent attachment sites to pursue in IDP drug discovery campaigns and the rational design of novel IDP covalent ligands with therapeutic potential.

## Methods

### Parameterization of covalent cysteine adducts of EPI-002 and EPI-7170

We parameterized covalent cysteine adducts of EPI-002 (“CYS:EPI-002”) and EPI-7170 (“CYS:EPI-7170”) as new residues within the a99SB-*disp* force field^44^. The structures of these covalent adduct residues are depicted in Figure 1A. We initially parameterized CYS:EPI-002 and CYS:EPI-7170 residues as individual peptide moieties with N-terminal acetyl (ACE) and C-terminal N-methyl (NME) capping groups using the general AMBER force field (GAFF1)^45^. We refer to these molecules as ACE-CYS:EPI-002-NME and ACE-CYS:EPI-7170-NME. GAFF1 was chosen for consistency with our previous MD study of non-covalent binding of EPI-002 and EPI-7170 to Tau-5_R2 R3_ ^29^. GAFF1 parameters were obtained with ACPYPE^57^. We ran explicit solvent REST2 simulations of these molecules with a99SB-*disp* water using 8 replicas spanning solute temperatures from 300-600K in 4.0nm water boxes and selecting all non-water atoms as solute for tempering. We also ran REST2 simulations of EPI-002 and EPI-7170 parameterized with GAFF1 in a 4.0nm water box using the same solute temperature ladder. We compared the distributions of the dihedral angles in the sidechain EPI-002 and EPI-7170 moieties of the ACE-CYS:EPI-002-NME and ACE-CYS:EPI-7170-NME observed in the 300K REST2 replicas to the distributions of the dihedral angles observed in the 300K replicas of EPI-002 and EPI-7170 REST2 simulations to ensure there were no unphysical deviations in the conformational ensembles resulting from the GAFF1 parameterization of the (*CH*_2_ − *S* − *CH*_2_) linkage. We observed close agreement between these distributions of dihedral angles (data not shown).

We then proceeded to adjust the petide backbone parameters of ACE-CYS:EPI-002-NME and ACE-CYS:EPI-7170-NME to be consistent with a99SB-*disp* force field backbone parameters. We did so by introducing new residues into the a99SB-*disp* force field, which we refer to as CYE2 (for the CYS:EPI-002 adduct) and CYE7 (for the CYS:EPI-7170 adduct). These residues used the a99SB-*disp* CYS residue force field bonded parameters (bond lengths, bond angles, diehdral angles, improper dihedral angles) and non-bonded parameters (partial charges, Lennard-Jones van der Waals atom types) for all peptide backbone atoms. We also used the a99SB-*disp* CYS force field bonded parameters and Lennard-Jones atom types for the the sidechain *C_β_* and S atoms of CYE2 and CYE7. Starting with the partial charges of the a99SB-*disp* CYS sidechain *C_β_* and S atoms and GAFF1 partial charges for all other sidechain atoms, we manually adjusted the partial charges within the connecting (*CH*_2_ − *S* − *CH*_2_) regions of the CYE2 and CYE7 sidechains to maintain charge neutrality for the new residues with minimal deviations from the initial partial charges. The final parameters of the CYE2 and CYE7 residues are included in the supporting information (SI Tables 3-9) and GROMACS parameters files are provided in the accompanying github repository (https://github.com/paulrobustelli/Zhu_Robustelli_AR_Covalent_Adducts_24). We ran REST2 simulations of ACE-CYE2-NME and ACE-CYE7-NME using the newly parameterized CYE2 and CYE7 residues and the standard a99SB-*disp* ACE and NME parameters. We compared the sidechain dihedral distributions observed in the 300K replicas to those observed in the 300K replicas of the ACE-CYS:EPI-002-NME and ACE-CYS:EPI-7170-NME REST2 simulations run with GAFF1 parameters and observed excellent agreement (data not shown). We also compared the backbone *ϕ*/*ψ* and sidechain *χ*_1_ dihedral angles of ACE-CYE2-NME and ACE-CYE7-NME to those observed in a REST2 simulation of ACE-CYS-NME using a99SB-*disp* force field parameters and observed close agreement (data not shown).

### Molecular Dynamics Simulations

All MD simulations were performed using GROMACS 2019.2^58,59^ patched with PLUMED v2.6.0^60^. REST2 Simulations of apo Tau-5_R2 R3_ (AR residues L391-G446, capped with ACE and NH2 groups), Tau-5_R2 R3_ in the presence of EPI-002 and Tau-5_R2 R3_ in the presence of EPI-7170 were previously reported^29^. These simulations were run with the a99SB-*disp* protein force field and a99SB-*disp* water model ^44^ and GAFF1 ligand parameters generated by ACPYPE^45,57^.

Simulations of covalent adducts of EPI-002 and EPI-7170 bound to CYS404 of Tau-5_R2 R3_ (“Tau-5_R2 R3_-CYS404:EPI-002” and “Tau-5_R2 R3_-CYS404:EPI-7170”) were set-up using an identical REST2 protocol to match our previously published work ^29^. Starting structures of Tau-5_R2 R3_-CYS404:EPI-002 and Tau-5_R2 R3_-CYS404:EPI-7170 were built using PyMOL^61^ to attach EPI-002 and EPI-7170 onto residue CYS404 of Tau-5_R2 R3_ structures previously used as starting structures for apo and non-covalent ligand binding simulations^29^, omitting structures where covalent attachment introduced large steric clashes. Tau-5_R2 R3_-CYS404:EPI-002 and Tau-5_R2 R3_-CYS404:EPI-7170 were parameterized using the a99SB-*disp* protein force field for canonical protein residues, and parameters for cysteine covalent adducts CYS:EPI-002 and CYS:EPI-7170 derived as described in the section “Parameterization of covalent cysteine adducts of EPI-002 and EPI-7170”.

Each system was solvated with 13200 water molecules in a cubic box with a length of 7.5 nm and neutralized with a salt concentration of 20 mM NaCl by 8 Na^+^ ions and 5 Cl^−^ ions. Energy minimization of each system was performed with the steepest descent minimization algorithm until the maximum force obtained was smaller than 1000.0 *kJ/*(*mol/nm*). Equilibration was first performed in the NVT ensemble for 2000 ps at the temperature of 300K using the Berendsen thermostat ^62^. Systems were further equilibrated in the NPT ensemble for 200 ps at a target pressure of 1 bar with the temperature at 300K maintained by Berendsen thermostat, with position restraints added to all heavy atoms. Bond lengths and bond angles of protein and ligand atoms were constrained with the LINCS algorithm ^63^ and water constraints were applied using the SETTLE algorithm ^64^. Canonical sampling in the NVT ensemble algorithms was obtained using the Bussi et al. velocity rescaling thermostat^65^ with a 2 fs timestep. The PME algorithm ^66^ was utilized for electrostatics with a grid spacing of 1.6 nm. Van der Waals forces were calculated using a 0.9 nm cut-off length.

The REST2 algorithm ^46,47^ was utilized with exchanges attempted every 80 ps. All covalent adduct atoms were selected as solute with a 16-replica solute temperature ladder from 300–500K. Simulations of Tau-5_R2 R3_-CYS404:EPI-002 were run for 4.8*µ*s/replica (aggregate simulation time of 77*µ*s) and simulations of Tau-5_R2 R3_-CYS404:EPI-7170 were run for 4.5*µ*s/replica (aggregate simulation time of 72*µ*s). Previously reported REST2 simulations of apo Tau-5_R2 R3_, Tau-5_R2 R3_ and EPI-002 and Tau-5_R2 R3_ and EPI-7170 were simulated for 4.6, 4.0, 4.5*µ*s per replica respectively, for total simulation times of 74, 64 and 72*µ*s respectively. Frames were saved every 80 ps for analysis. Secondary structure populations were calculated from MD trajectories using the DSSP algorithm^67^. Analyses were run utilizing MDtraj^68^ and the NumPy^69^ python package.

### Statistical error estimates

Statistical error estimates of the simulated properties from MD simulations were calculated using a blocking analysis^48^ with an optimal block size selection determined, using the *pyblock* python package^49^. In this procedure, the trajectory is divided into a given number of equally sized “blocks”, average values of simulated quantities are computed for each block, and the standard error of the average values calculated across all blocks is used as an error estimate. Optimal block size is selected to minimize the estimated error of the standard error across blocks^49^.

### *Sα α*-helical order parameter

The *α*-helical order parameter *Sα*, measures the similarity each seven-residue segment in a protein to an ideal helical structure (*φ* = −57,*ψ* = −47) ^52^. *Sα* is calculated according

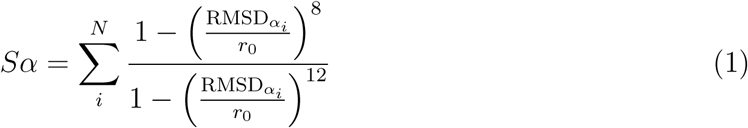

where the sum is over N consecutive seven-residue segments, RMSD*_αi_* is the C*α*-RMSD between an ideal *α*-helical geometry a seven-residue fragment (spanning from residue i to residue i + 6), and *r*_0_ = 1.0 ^°^A. When *r*_0_ = 1.0 ^°^A, a seven-residue fragment with a value of RMSD*_α_<* 0.5 ^°^A contributes a value of ∼1 to the *Sα* sum, a seven-residue fragment with a value of RMSD*_α_*= 1.1 ^°^A contributes a value of ∼0.5 to the *Sα* sum, and a seven-residue fragment with a value of RMSD*_α_ >* 3.0 ^°^A contributes a value of ∼0 to the *Sα* sum. The value of *Sα* for a protein conformation can therefore be interpreted as a proxy for the number of seven-residue fragments closely resembling an ideal helical conformation. A completely helical conformation of the 56-residue Tau-5_R2 R3_ construct has an *Sα* value of 50, and a Tau-5_R2 R3_ with no helical content has an *Sα* value of 0.

### Clustering conformational ensembles of Tau-5_R2 R3_ with t-stochastic neighbor embedding (t-SNE)

Given a set of *n* conformations *X* = {*x*_1_*, x*_2_*, …, x_n_*} with *d*-dimensional input features, t-SNE finds a lower-dimensional embedding *Y* = {*y*_1_*, y*_2_*, …, y_n_*} with *s*-dimensional features (where typically *s* = 2 or *s* = 3) based on the similarity and dissimilarity of the conformations. t-SNE seeks to identify a low-dimensional embedding such that the conditional probability *p*_(_*_i_*_|_*_j_*_)_ of finding two points *x_i_* and *x_j_* in the same local neighborhood in the high-dimensional feature space is as close as possible to the conditional probability *q*_(_*_i_*_|_*_j_*_)_ of finding two points *y_i_* and *y_j_* in the same local neighborhood in the low-dimensional feature space. *p*_(_*_i_*_|_*_j_*_)_ is defined in the high-dimensional feature space using Gaussian functions

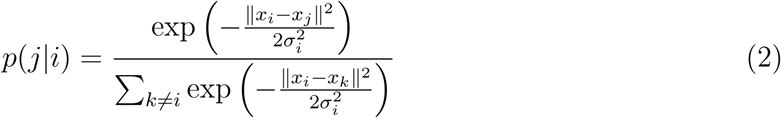

while *q*_(_*_i_*_|_*_j_*_)_ is defined in the low-dimensional feature space using a heavy tailed Student’s t-distribution

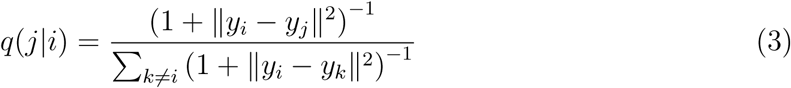

and all values *p*_(_*_i_*_|_*_i_*_)_ = 0 and *q*_(_*_i_*_|_*_i_*_)_ = 0. To ensure pairwise symmetry, joint probabilities are calculated from conditional probabilities according to

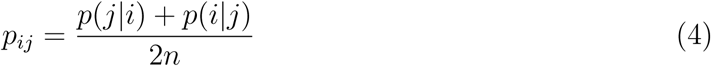

The size of the local neighborhood in the high dimensional feature space is controlled through the bandwidth of the Gaussian kernels *σ_i_* in Eq. 2. The values of *σ_i_* are defined such that the entropy of the conditional distributions *P_i_* match a predefined entropy determined by a preselected perplexity (*perp*) hyperparameter

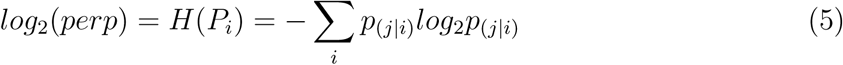

The difference between the high-dimensional joint probability distribution *P* and lowdimensional joint probability distribution *Q* is calculated as a Kullback–Leibler (KL) divergence over all data points

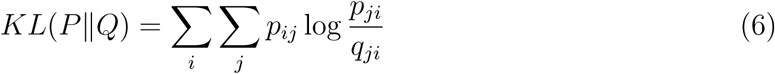

The spatial distribution of points in the low-dimensional embedding is initialized randomly, and the final distribution is determined by iteratively rearranging the points using gradient descent optimization to minimize the KL divergence for each selected value of the perplexity hyperparameter. *k* -means clustering is then applied to partition points in the low-dimensional embedding into *N* non-overlapping clusters. The quality of cluster assignments is assessed by calculating the silhouette score for each data point *i*

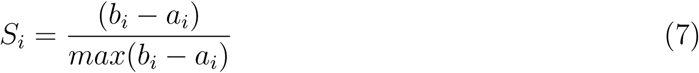

where *a_i_* is the intracluster distance defined as the average distance to all other points in the cluster to which it belongs and *b_i_* represents the intercluster distance measured as the average distance to the closest cluster of data point *i* excluding the cluster that it is assigned to. Typically the silhouette score ranges between 1 and -1, where a high value indicates good clustering and values closer to 0 indicate poor clustering. A silhouette score with a negative value indicates the clustering configuration is wrong or inappropriate.

The distance between points in Eq. 7 is usually measured in terms of the Euclidean distance metric. Since the clusters are identified in a reduced representation with t-SNE, computing the silhouette score based only on the distances in the low-dimensional space (*S_ld_*) may be misleading if the points are poorly embedded during the dimensional reduction step by t-SNE. We therefore also check the quality of clustering with respect to the original distance in the high-dimensional space (*S_hd_*), and evaluate an integrated silhouette score defined as (*S_i_* = *S_ld_* × *S_hd_*).

We performed t-SNE based clustering on i) a merged ensemble containing the Tau-5_R2 R3_-CYS404:EPI-002 and Tau-5_R2 R3_-CYS404:EPI-7170 covalent adduct ensembles and ii) a merged ensemble containing all frames (bound and unbound) from EPI-002 and EPI-7170 non-covalent ligand-binding simulations. For clustering, we down sampled each individual ensemble to contain 5,000 frames, and each merged ensemble contained 10,000 frames total. We calculated the root-mean-square deviation (RMSD) of backbone C*α* coordinates between each pair of conformations in each merged ensemble and utilize the resulting all-to-all C*α* RMSD matrix as input for dimensionality reduction with t-SNE as described above. The distance ∥,*x_i_* − *x_j_* ∥, between two conformations *x_i_* and *x_j_* in the high-dimensional space is therefore defined by the difference in the length 10,000 vectors containing the values of the C*α* RMSD of each conformation to all other conformations in the merged ensemble.

t-SNE is highly sensitive to the choice of the value of the perplexity hyperparameter (Eq. 5), which can be interpreted as a smoothed measure of the number of nearest neighbors that each point is attracted to during dimensionality reduction. In our clustering protocol, perplexity therefore controls the granularity of the resulting cluster assignments, with smaller perplexity values identifying smaller and more structurally homogeneous clusters. We identify locally optimal values of perplexity and the number of clusters *N* using the silhouette score (Eq. 7) as described previously^30^. We perform t-SNE dimensionality reduction to obtain 2D projections of our data with a range of values of perplexity. For each 2D projection, we subsequently perform *k* -means clustering of the data points using a range of values of the number of clusters (*N*). We evaluate the silhouette score of the cluster assignments obtained for each pair of perplexity and *N* values to identify optimal parameters for clustering at each desired level of resolution (SI Figure 9). For each pair of ensembles analyzed, we consider cluster assignments obtained at two levels of resolution. We identify the cluster assignment that produced the highest silhouette score when we restrict the number of clusters to *N* =4 (Figures 2-4, Tables 1-2, SI Figures 9-16) and we identify a more granular cluster assignment that produces the highest silhouette score when *N* is restrictied to values between *N* =10 and *N* =20 (Figure 2, SI Figures 9-10, SI Figures 17-30, SI Tables 1-2).

## Code Availability

All analysis code used in this investigation is freely available from the GitHub repository https://github.com/paulrobustelli/Zhu_Robustelli_AR_Covalent_Adducts_24.

## Data Availability

All trajectories, simulation input files, and force field parameters created in this investigation are freely available from the GitHub repository https://github.com/paulrobustelli/ Zhu_Robustelli_AR_Covalent_Adducts_24. We provide conformational ensembles of the Tau-5_R2 R3_-CYS404:EPI-002 and Tau-5_R2 R3_-CYS404:EPI-7170 covalent adducts and previously reported conformational ensembles of apo Tau-5_R2 R3_ and non-covalent ligand binding simulations of Tau-5_R2 R3_ with EPI-002 and Tau-5_R2 R3_ with EPI-7170^29^. We provide force field parameters for the covalently modified cysteine residues CYS:EPI-002 and CYS:EPI-7170 and all covalent adduct GROMACS simulation input files. GROMACS simulation input files for previously reported REST2 simulations of apo Tau-5_R2 R3_ and non-covalent ligand binding simulations of Tau-5_R2 R3_ with EPI-002 and EPI-7170 are freely available from the GitHub repository https://github.com/paulrobustelli/AR_ligand_binding

## Acknowledgement

This work was supported by the National Institutes of Health under award R35GM142750 and the China Scholarship Council (CSC ID: 201906320040) The authors thank Xavier Salvatella and Stase Bielskute for valuable conversations and feedback.

