## Supporting Information for "Covalent adducts formed by the androgen receptor transactivation domain and small molecule drugs remain disordered"

### MD Simulation Convergence Analyses

The convergence of each covalent adduct simulation was assessed by comparing the and secondary structure propensities and the populations of intramolecular protein-ligand contacts for each temperature rung in the REST2 simulation in SI Figure 1, SI Figure 3, SI Figure 5 and SI Figure 7. The relative smooth temperature dependence of these properties suggest that simulations are reasonably well converged. The same analyses were performed on the demultiplexed replicas, which follow each independent replica through temperature space, to determine if any individual replicas became stuck in local minima as they diffuse through the temperature ladder in SI Figure 2, SI Figure 4, SI Figure 6, and SI Figure 8. We found the statistical fluctuations to be reasonably well converged among demultiplexed replicas.

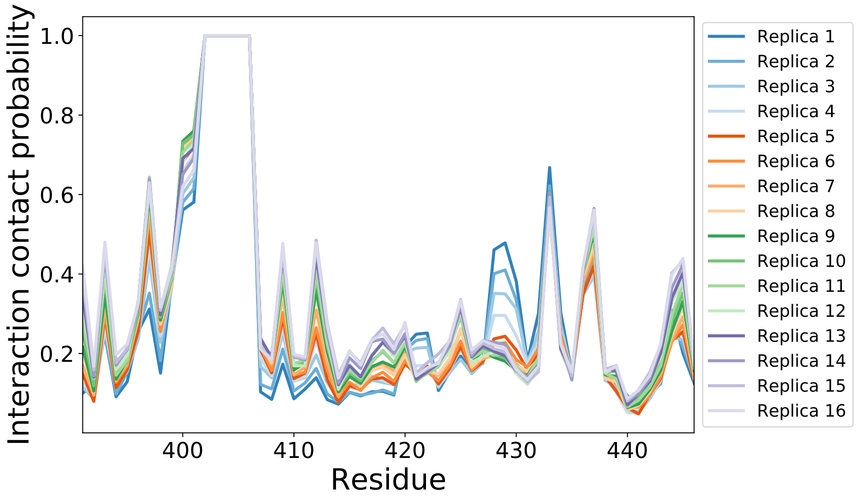


**Supplementary Figure 1: Intramolecular protein-ligand contacts observed in a REST2 MD simulation of Tau-5$_{\mathbf{R2_R3}}$-CYS404:EPI-002.** Intramolecular contact probabilities between the covalently modified CYS404:EPI-002 residue and Tau-5$_{R2_R3}$ residues observed in the 16 solute temperature rungs spanning 300K-500K. Contacts between CYS404:EPI-022 and Tau-5$_{R2_R3}$ residues are defined using a cutoff of 6Å between heavy atoms.

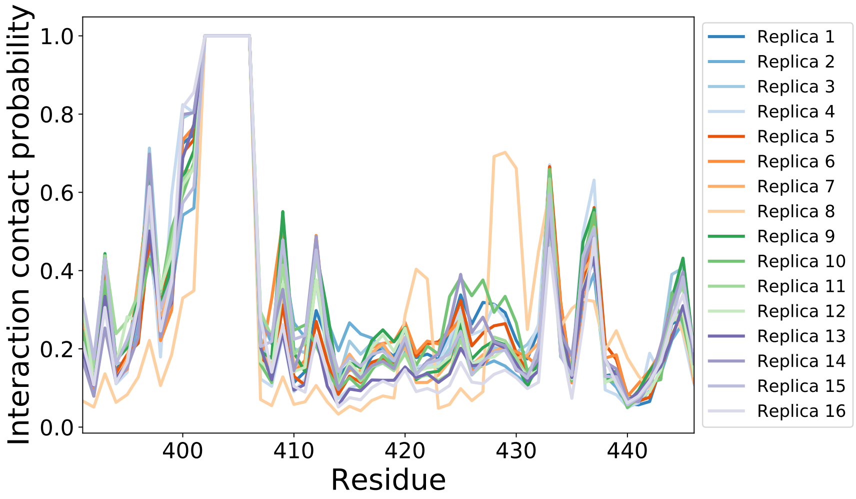


**Supplementary Figure 2: Intramolecular protein-ligand contacts observed in the demultiplexed replicas of a REST2 MD simulation of Tau-5$_{\mathbf{R2_R3}}$-CYS404:EPI-002.** Intramolecular contact probabilities between the covalently modified CYS404:EPI-002 residue and Tau-5$_{R2_R3}$ residues observed in the 16 demultiplexed replicas. Contacts between CYS404:EPI-002 and Tau-5$_{R2_R3}$ residues are defined using a cutoff of 6Å between heavy atoms.

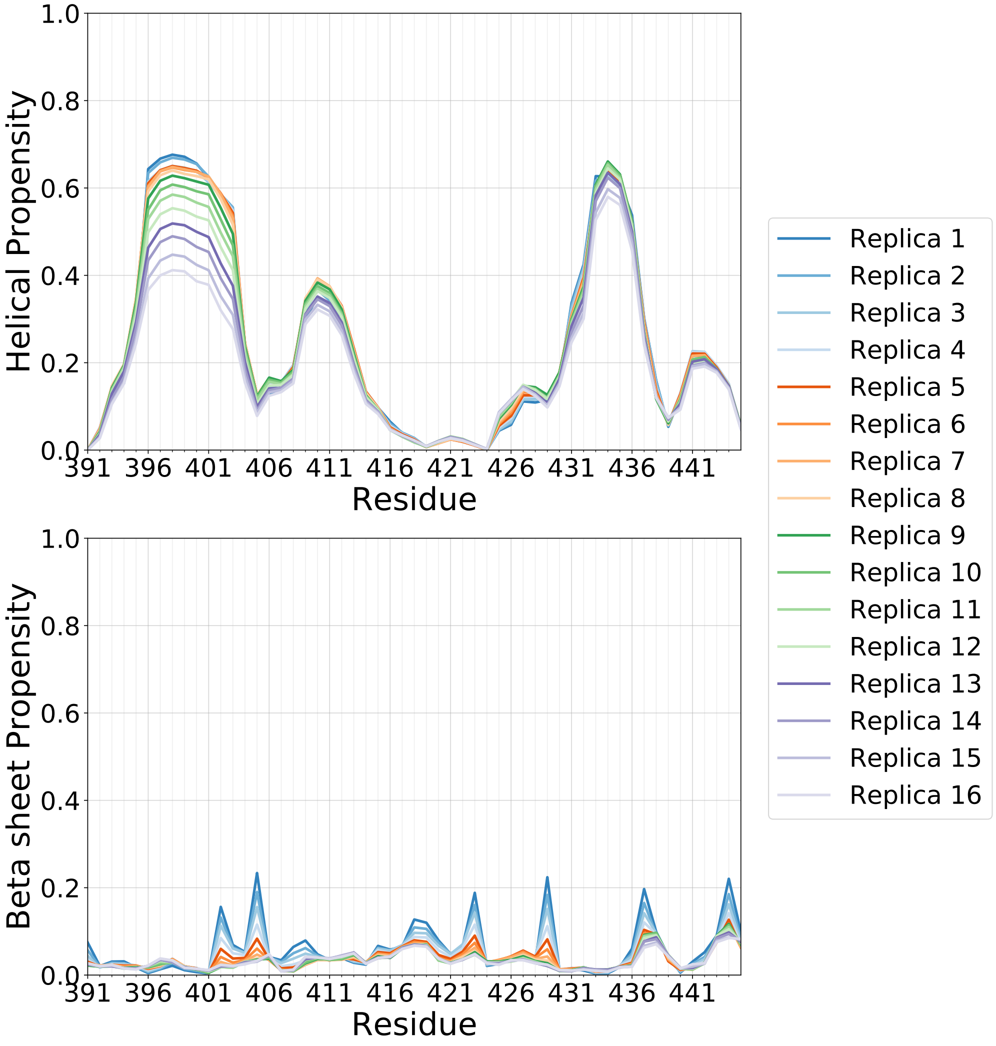


**Supplementary Figure 3: Secondary structure propensities observed in a REST2 MD simulation of Tau-5$_{\mathbf{R2_R3}}$-CYS404:EPI-002.** Comparison of $\alpha$-helical and $\beta$-sheet propensities of Tau-5$_{\mathrm{R2_R3}}$ observed in the 16 solute temperature rungs spanning 300K-500K. Secondary structure content is calculated by the DSSP algorithm.

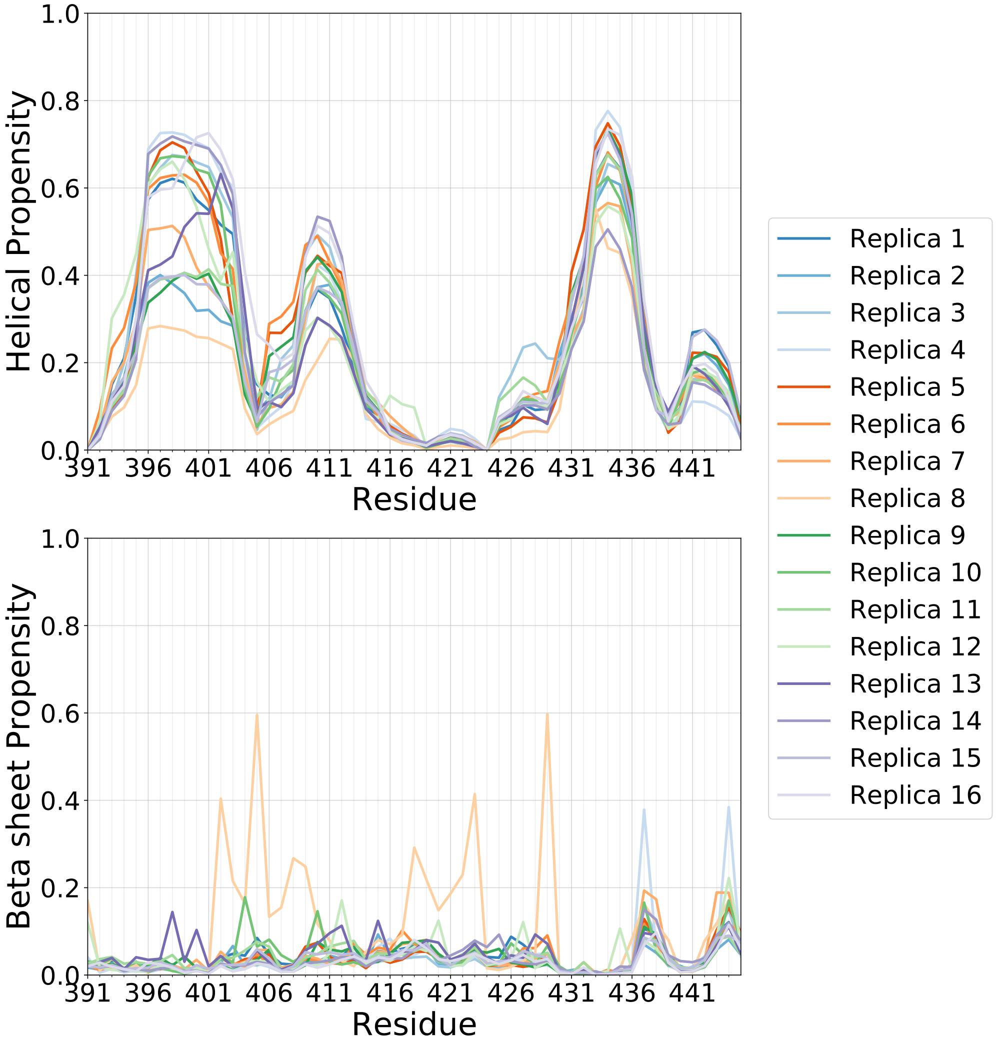


**Supplementary Figure 4: Secondary structure propensities observed in the demultiplexed replicas of a REST2 MD simulation of Tau-5$_{\mathbf{R2_R3}}$-CYS404:EPI-002.** Comparison of $\alpha$-helical and $\beta$-sheet propensities of Tau-5$_{\mathrm{R2_R3}}$ observed in the 16 demultiplexed replicas of a REST2 MD simulation of Tau-5$_{\mathrm{R2_R3}}$-CYS404:EPI-002. Secondary structure content is calculated by the DSSP algorithm. Replica 8 has the largest deviations from the average secondary structure propensities of the demultiplexed replicas.

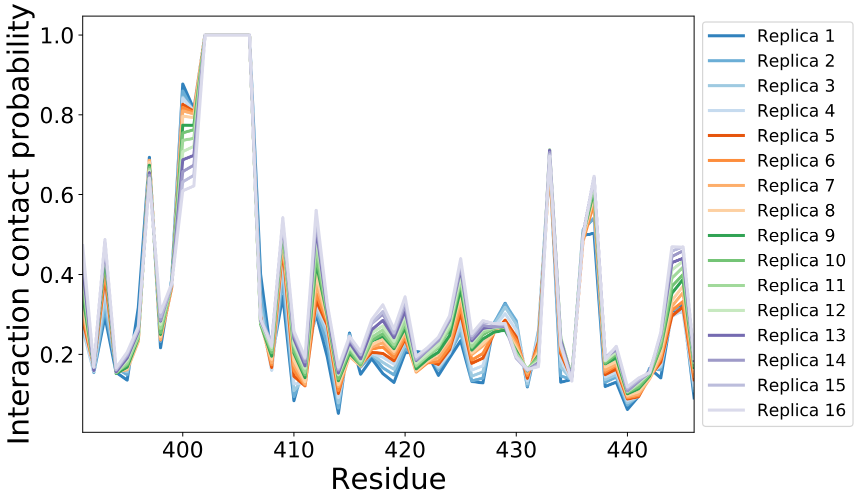


**Supplementary Figure 5: Intramolecular protein-ligand contacts observed in a REST2 MD simulation of Tau-5$_{\mathbf{R2_R3}}$-CYS404:EPI-7170.** Intramolecular contact probabilities between the covalently modified CYS404:EPI-7170 residue and Tau-5$_{\mathrm{R2_R3}}$ residues observed in the 16 solute temperature rungs spanning 300K-500K. Contacts between CYS404:EPI-7170 and Tau-5$_{\mathrm{R2_R3}}$ residues are defined using a cutoff of 6Å between heavy atoms.

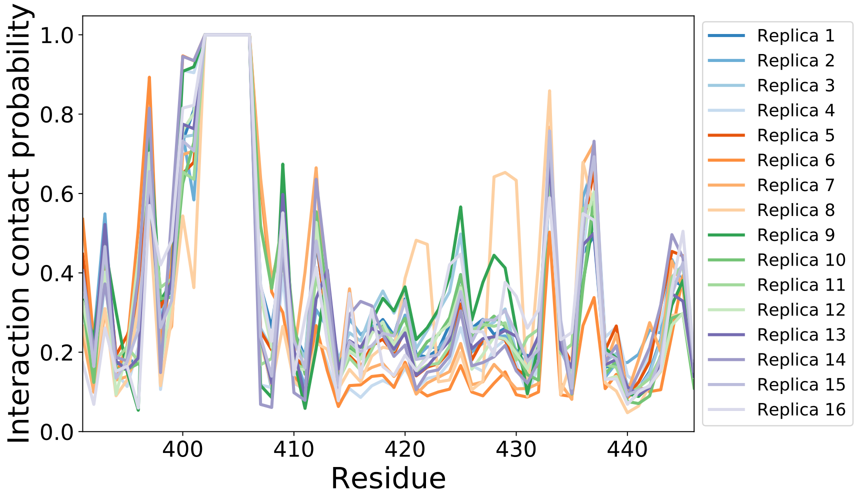


**Supplementary Figure 6: Intramolecular protein-ligand contacts observed in the demultiplexed replicas of a REST2 MD simulation of Tau-5$_{\mathbf{R2_R3}}$-CYS404:EPI-7170.** Intramolecular contact probabilities between the covalently modified CYS404:EPI-7170 residue and Tau-5$_{\mathrm{R2_R3}}$ residues observed in the 16 demultiplexed replicas. Contacts between CYS404:EPI-7170 and Tau-5$_{\mathrm{R2_R3}}$ residues are defined using a cutoff of 6Å between heavy atoms.

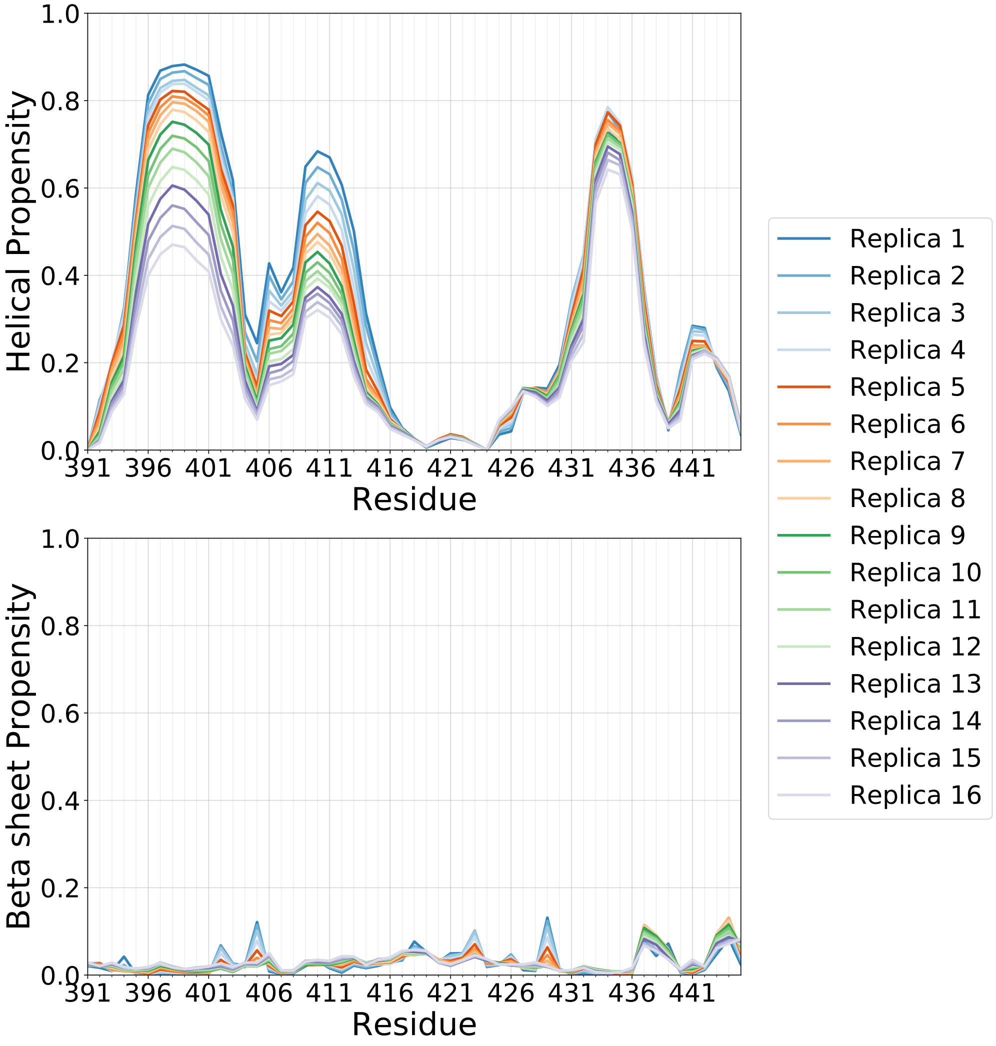


**Supplementary Figure 7: Secondary structure propensities observed in a REST2 MD simulation of Tau-5$_{\mathbf{R2_R3}}$-CYS404:EPI-7170.** Comparison of $\alpha$-helical and $\beta$-sheet propensities of Tau-5$_{\mathrm{R2_R3}}$ observed in the 16 solute temperature rungs spanning 300K-500K. Secondary structure content is calculated by the DSSP algorithm.

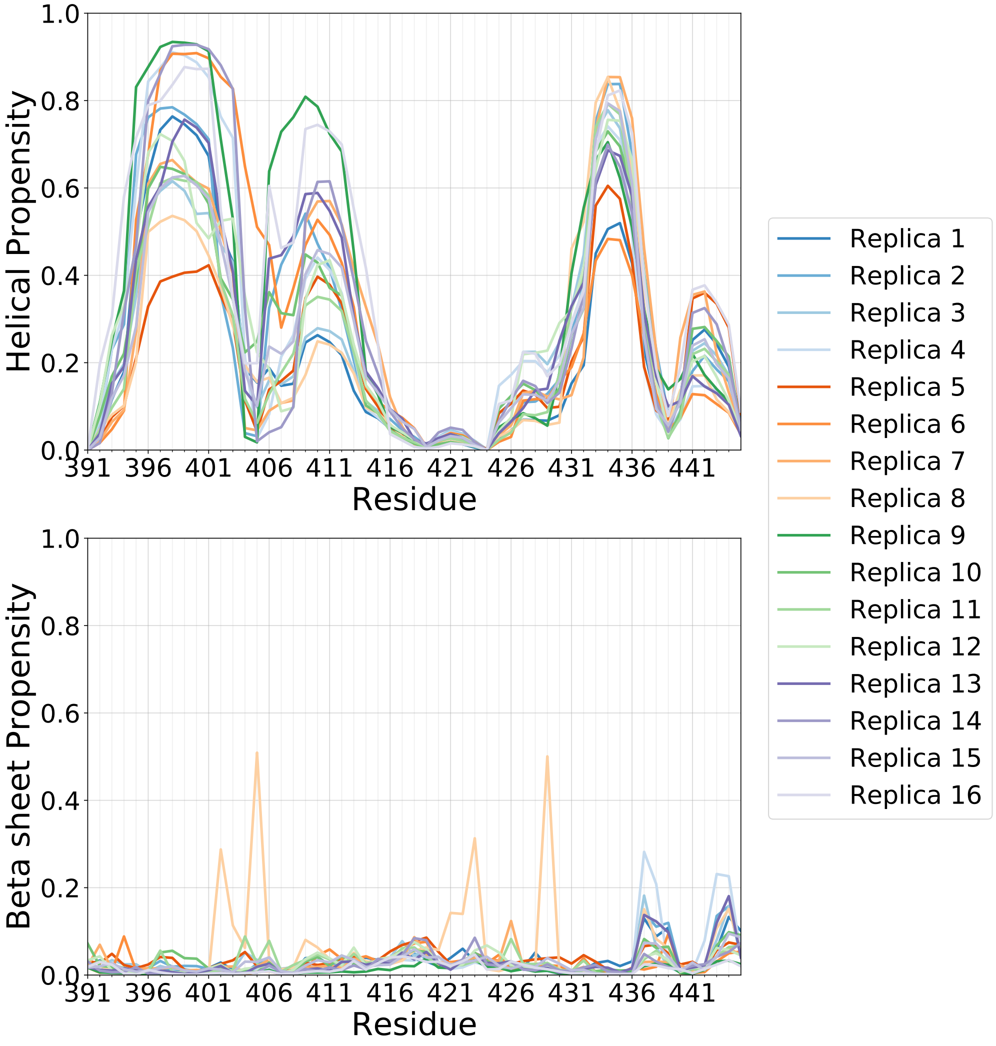


**Supplementary Figure 8: Secondary structure propensities observed in the demultiplexed replicas of a REST2 MD simulation of Tau-5$_{\mathbf{R2_R3}}$-CYS404:EPI-7170.** Comparison of $\alpha$-helical and $\beta$-sheet propensities of Tau-5$_{\mathrm{R2_R3}}$ observed in the 16 demultiplexed replicas of a REST2 MD simulation of Tau-5$_{\mathrm{R2_R3}}$-CYS404:EPI-7170. Secondary structure content is calculated by the DSSP algorithm. Replica 8 has the largest deviations from the average secondary structure propensities of the demultiplexed replicas.

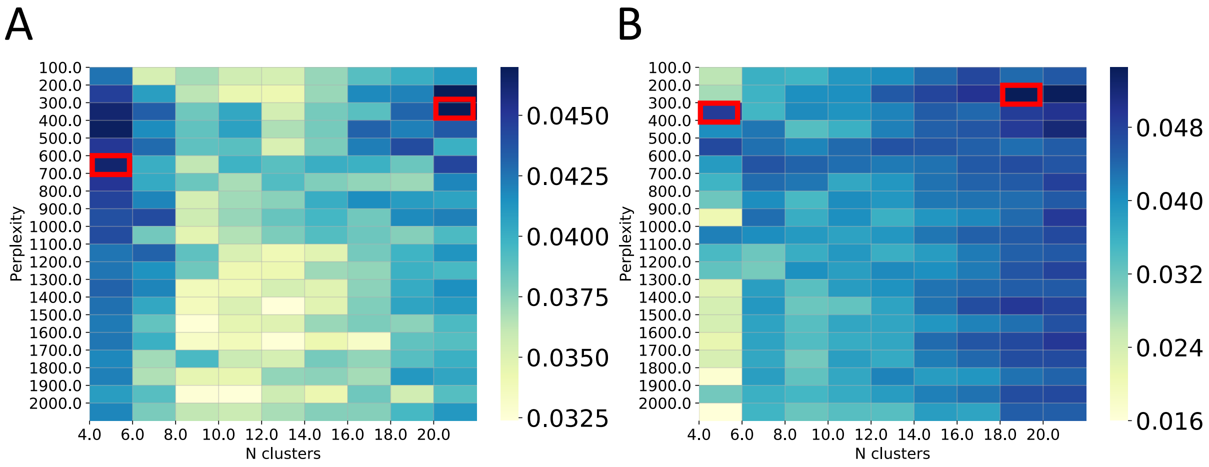


**Supplementary Figure 9: t-SNE clustering hyperparameter selection.** Identification of optimal t-SNE clustering hyperparameters perplexity (*perp*) and number of clusters ($N$) for a merged ensemble containing the Tau-$5_{R2\_R3}$-CYS404:EPI-002 and Tau-$5_{R2\_R3}$-CYS404:EPI-7170 covalent adduct ensembles **(A)** and a merged ensemble containing all frames from non-covalent Tau-$5_{R2\_R3}$ EPI-002 and EPI-7170 ligand-binding simulations **(B)**. We perform t-SNE clustering of each merged ensemble with a range of values of *perp* and $N$ and compute the integrated silhouette score (Eq. 7) of the resulting cluster assignments. We identify two locally optimal t-SNE projections for coavalent adduct ensembles (*perp* = 600, $N$=4 clusters and *perp* = 300, $N$=20 clusters) and two locally optimal t-SNE projections for ensembles from non-covalent binding simulations (*perp* = 300, $N$=4 clusters and *perp* = 200, $N$=18 clusters) highlighted by red boxes.

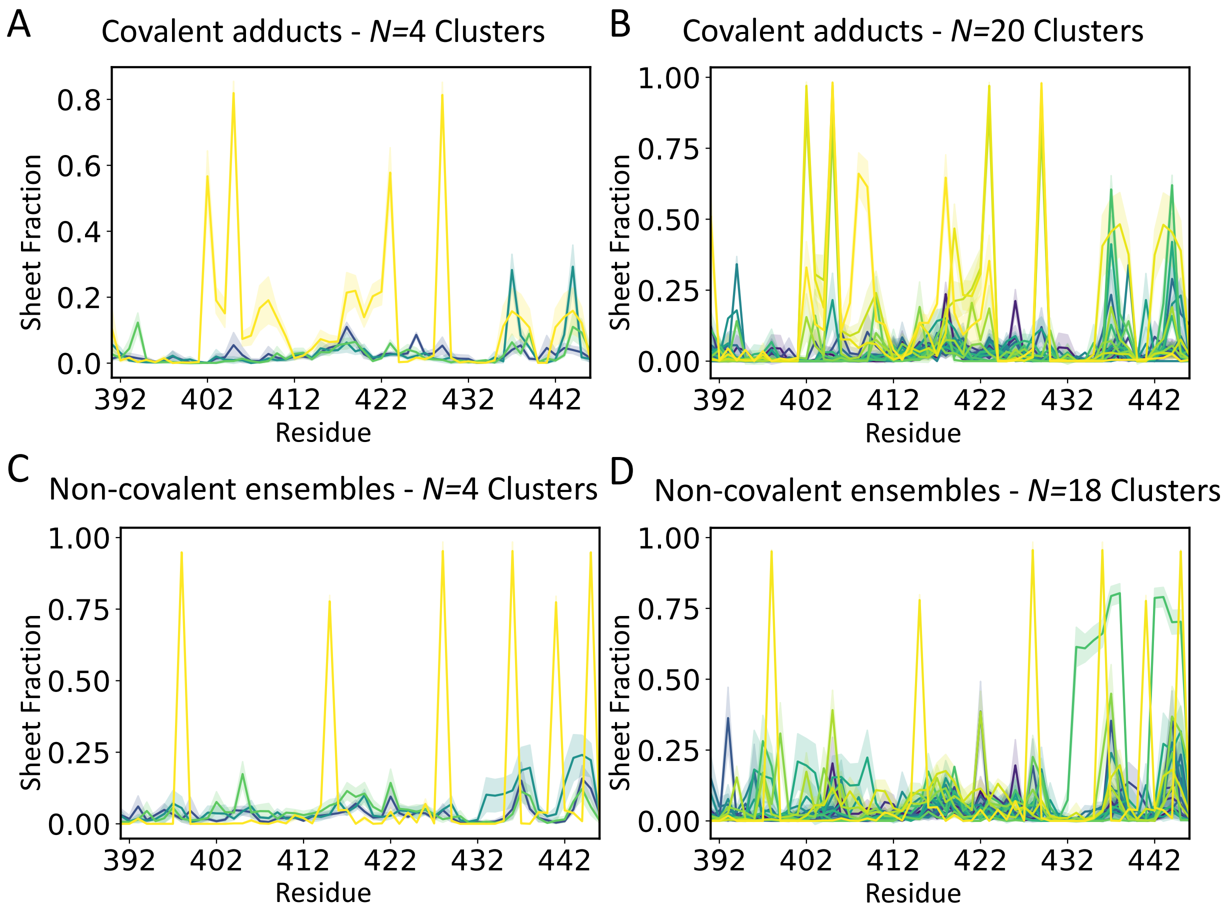


**Supplementary Figure 10: $\beta$-sheet propensities of clusters obtained from t-SNE clustering. A)** $\beta$-sheet propensities of covalent adduct clusters identified with $perp$ = 600 and $N$=4 clusters. **B)** $\beta$-sheet propensities of covalent adduct clusters identified with $perp$ = 300 and $N$=20 clusters. **C)** $\beta$-sheet propensities propensities of clusters identified from non-covalent ligand-binding simulations with $perp$ = 300 and $N$=4 clusters. **D)** $\beta$-sheet propensities propensities of clusters identified from non-covalent ligand-binding simulations with $perp$ = 200 and $N$=18 clusters.

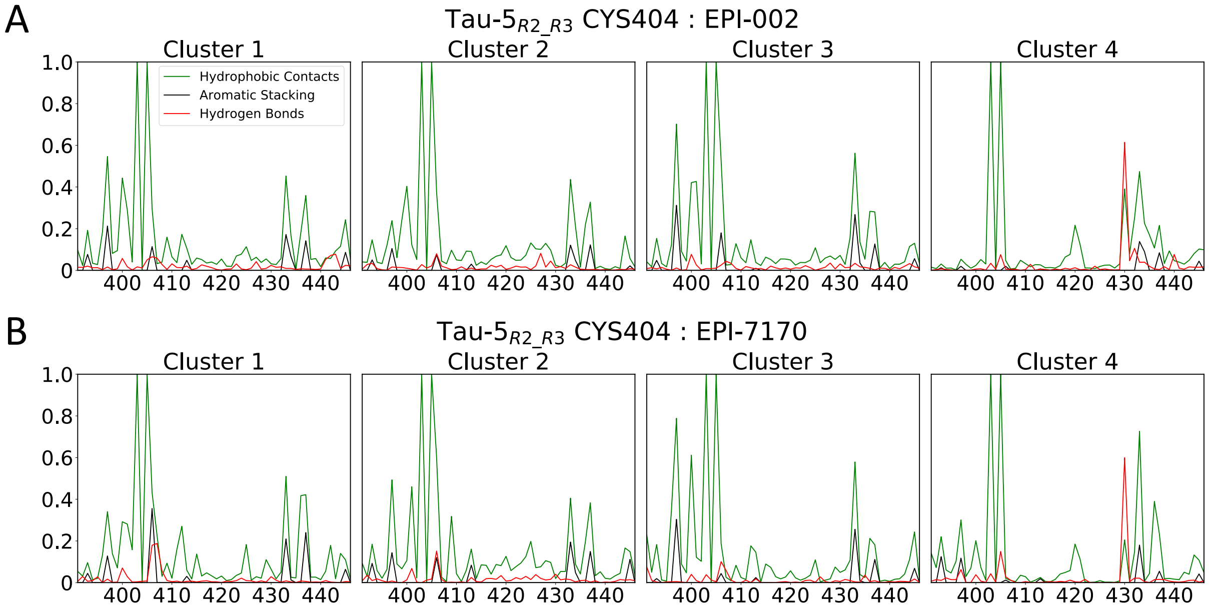


**Supplementary Figure 11: Populations of protein-ligand interactions in Tau-5$_{\mathbf{R2_R3}}$ covalent adduct conformational states identified by t-SNE clustering with $N$=4 clusters. A)** Populations of intramolecular interactions between CYS404:EPI-002 and Tau-5$_{R2_R3}$ residues in each cluster of the Tau-5$_{R2_R3}$-CYS404:EPI-002 ensemble. **B)** Populations of intramolecular interactions between CYS404:EPI-7170 and Tau-5$_{R2_R3}$ residues in each cluster of the Tau-5$_{R2_R3}$-CYS404:EPI-7170 ensemble.

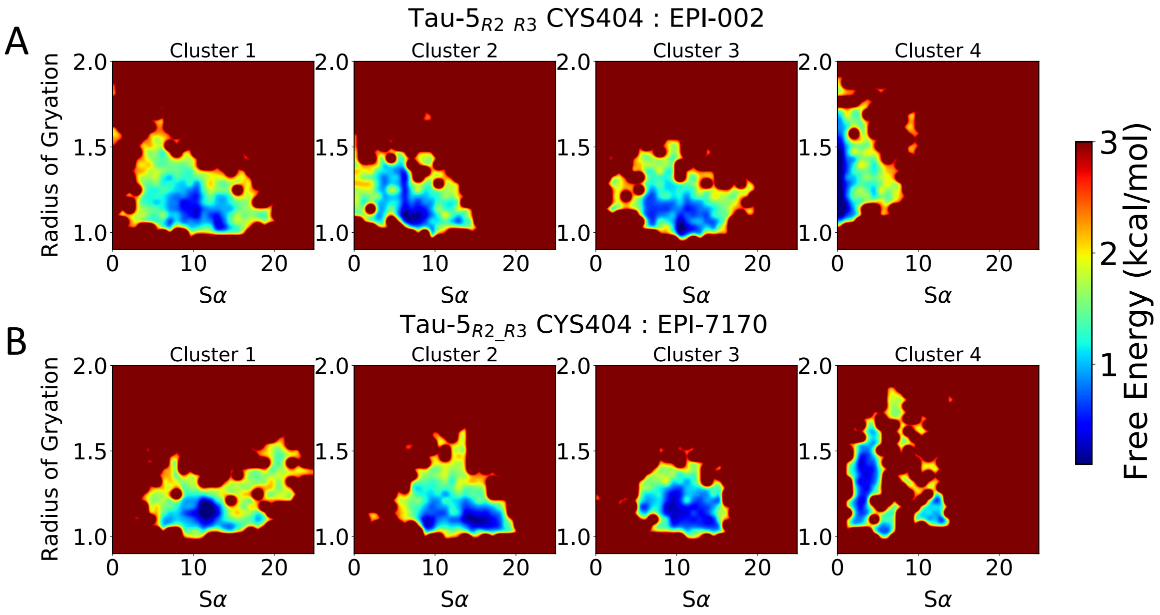


**Supplementary Figure 12: Free energy surfaces of Tau-5$_{\mathbf{R2_R3}}$ covalent adduct conformational states identified by t-SNE clustering with *N*=4 clusters.** Free energy surfaces as a function of the radius of gyration (reported in nm) and S$\alpha$ of Tau-5$_{\text{R2_R3}}$ conformations for covalent adduct conformational states identified by t-SNE clustering with $perp$ = 600 and $N$=4 clusters for **A)** Tau-5$_{\text{R2_R3}}$-CYS404:EPI-002 and **B)** Tau-5$_{\text{R2_R3}}$-CYS404:EPI-7170.

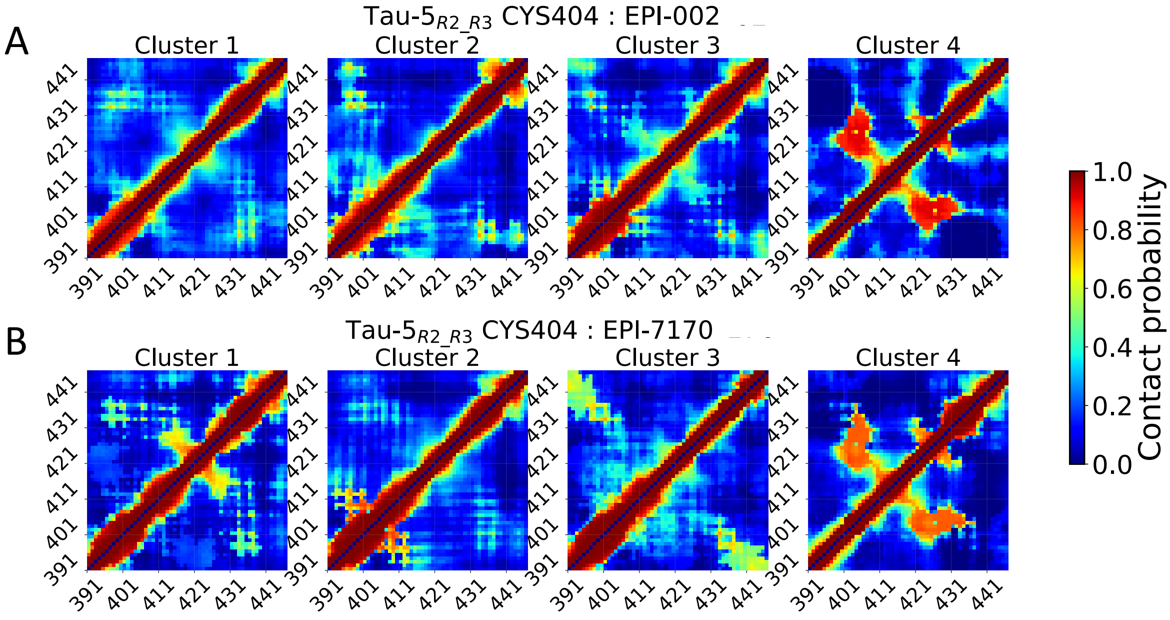


**Supplementary Figure 13: Intramolecular contact populations of Tau-5$_{\mathbf{R2_R3}}$ covalent adduct conformational states identified by t-SNE clustering with $N$=4 clusters.** Intramolecular contact populations of covalent adduct conformational states identified by t-SNE clustering with $perp = 600$ and $N$=4 clusters for **A)** Tau-5$_{\mathrm{R2_R3}}$-CYS404:EPI-002 and **B)** Tau-5$_{\mathrm{R2_R3}}$-CYS404:EPI-7170. Contacts between residues are defined using a distance cutoff of 12Å between closest heavy atoms.

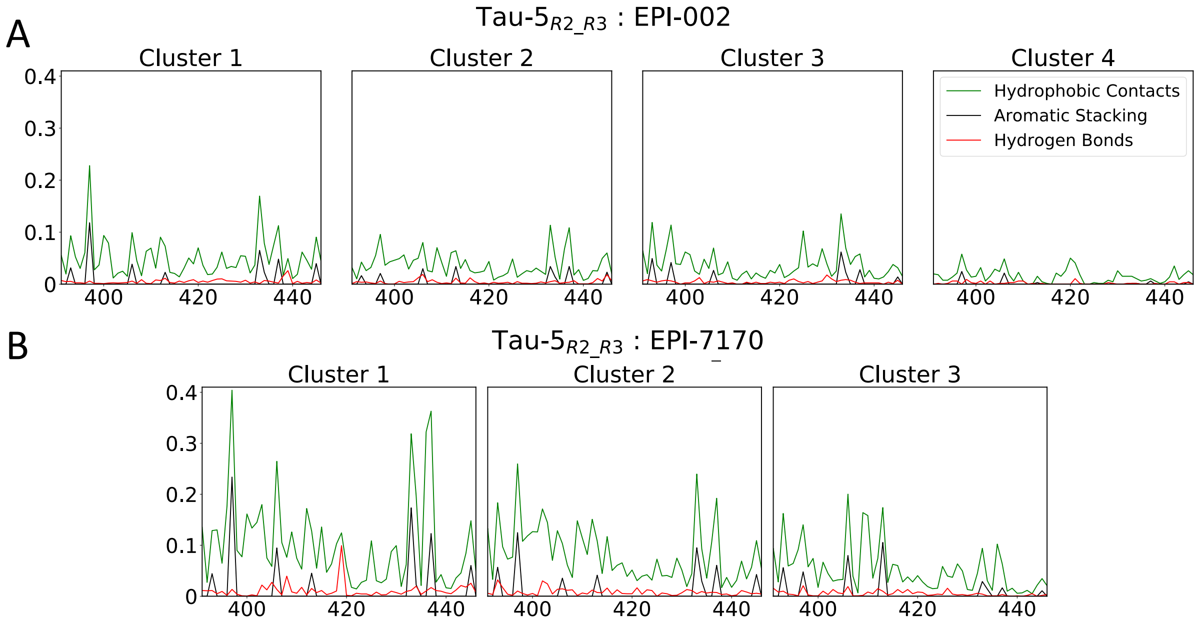


**Supplementary Figure 14: Populations of intermolecular protein-ligand interactions in Tau-5$_{\mathbf{R2_R3}}$ conformational states identified from non-covalent ligand-binding simulations of EPI-002 and EPI-7170 by t-SNE clustering with $\boldsymbol{N}$=4 clusters. A)** Populations of intermolecular interactions between EPI-002 and Tau-5$_{\mathrm{R2_R3}}$ in each cluster. **B)** Populations of intermolecular interactions between EPI-7170 and Tau-5$_{\mathrm{R2_R3}}$ in clusters 1-3. The interaction plot of cluster 4 is excluded because only one frame from was this ensemble was assigned to the this cluster.

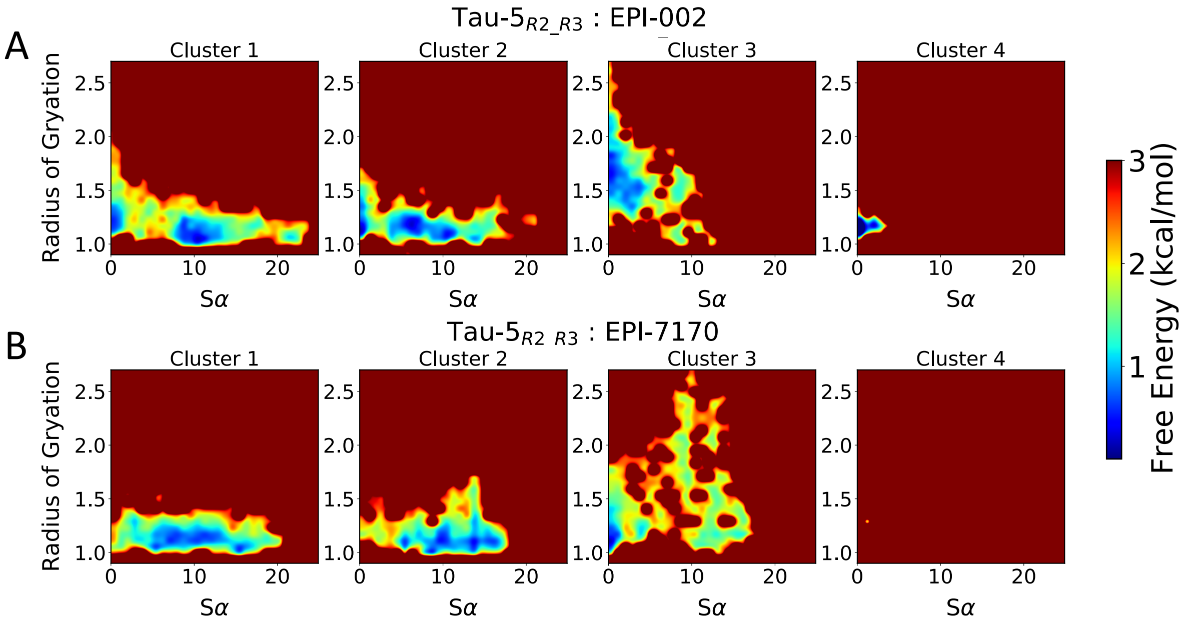


**Supplementary Figure 15: Free energy surfaces of Tau-5$_{\mathbf{R2_R3}}$ conformational states identified from non-covalent ligand binding simulations by t-SNE clustering with $\boldsymbol{N}$=4 clusters.** Free energy surfaces as a function of the radius of gyration (reported in nm) and S$\alpha$ of Tau-5$_{\mathrm{R2_R3}}$ conformations of conformational states identified from non-covalent ligand-binding simulations of EPI-002 and EPI-7170 by t-SNE clustering with $perp$ = 300 $N$=4 clusters for **A)** Tau-5$_{\mathrm{R2_R3}}$ in the presence of EPI-002 and **B)** Tau-5$_{\mathrm{R2_R3}}$ in the presence of EPI-7170.

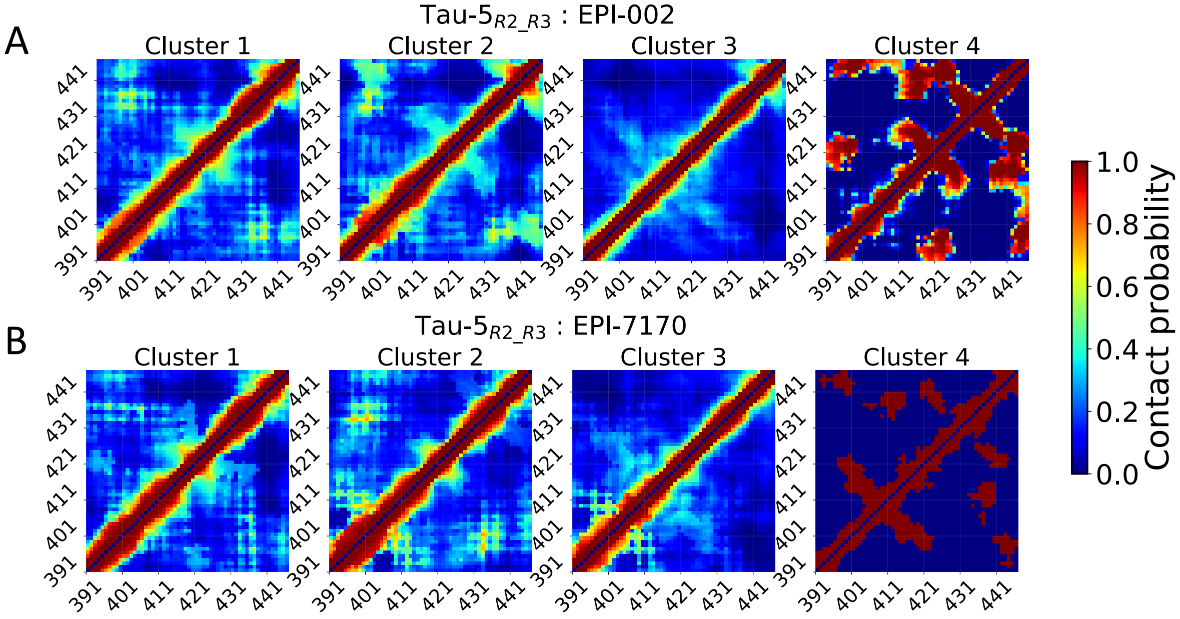


**Supplementary Figure 16: Intramolecular contact populations of Tau-5$_{\mathbf{R2_R3}}$ conformational states identified from non-covalent ligand binding simulations by t-SNE clustering with $N$=4 clusters.** Intramolecular contact populations of Tau-5$_{\mathrm{R2_R3}}$ conformational states identified by t-SNE clustering with $perp$ = 300 and $N$=4 clusters for **A)** Tau-5$_{\mathrm{R2_R3}}$ in the presence of EPI-002 and **B)** Tau-5$_{\mathrm{R2_R3}}$ in the presence of EPI-7170. We note that cluster 4 contains only one frame from this ensemble. Contacts between residues are defined using a distance cutoff of 12Å between closest heavy atoms.

**Supplementary Table 1:** Cluster population ($p$), helical globule population ($p_{Glob}$) and helix fraction (HF) of clusters obtained from t-SNE clustering of a merged ensemble of Tau-$5_{R2\_R3}$-CYS404:EPI-002 and Tau-$5_{R2\_R3}$-CYS404:EPI-7170 conformations with $N$=20 clusters. We compare the properties of the clusters in the merged ensemble to the properties of the clustered conformations from the the individual Tau-$5_{R2\_R3}$-CYS404:EPI-002 and Tau-$5_{R2\_R3}$-CYS404:EPI-7170 ensembles.

| Cluster | Merged Ensembles | | | Tau-$5_{R2\_R3}$-CYS404:EPI-002 | | | Tau-$5_{R2\_R3}$-CYS404:EPI-7170 | | |
|---|---|---|---|---|---|---|---|---|---|
| | $p$ | $p_{Glob}$ | HF | $p$ | $p_{Glob}$ | HF | $p$ | $p_{Glob}$ | HF |
| 1 | 0.06 | 0.80 | 0.42 | 0.01 | 0.29 | 0.20 | 0.11 | 0.85 | 0.44 |
| 2 | 0.07 | 0.88 | 0.36 | 0.04 | 0.66 | 0.05 | 0.10 | 0.96 | 0.39 |
| 3 | 0.06 | 0.82 | 0.35 | 0.06 | 0.76 | 0.29 | 0.07 | 0.87 | 0.40 |
| 4 | 0.04 | 0.90 | 0.35 | 0.01 | 0.66 | 0.27 | 0.07 | 0.95 | 0.37 |
| 5 | 0.05 | 0.55 | 0.34 | 0.05 | 0.58 | 0.29 | 0.06 | 0.51 | 0.38 |
| 6 | 0.04 | 0.75 | 0.33 | 0.07 | 0.75 | 0.33 | 0.02 | 0.72 | 0.32 |
| 7 | 0.05 | 0.65 | 0.04 | 0.07 | 0.58 | 0.29 | 0.03 | 0.80 | 0.32 |
| 8 | 0.06 | 0.94 | 0.30 | 0.08 | 0.92 | 0.30 | 0.04 | 0.97 | 0.37 |
| 9 | 0.05 | 0.92 | 0.30 | 0.00 | 0.67 | 0.30 | 0.10 | 0.93 | 0.30 |
| 10 | 0.04 | 0.84 | 0.30 | 0.06 | 0.87 | 0.30 | 0.02 | 0.71 | 0.29 |
| 11 | 0.05 | 0.79 | 0.30 | 0.07 | 0.78 | 0.28 | 0.03 | 0.80 | 0.34 |
| 12 | 0.06 | 0.59 | 0.09 | 0.06 | 0.38 | 0.28 | 0.06 | 0.81 | 0.36 |
| 13 | 0.04 | 0.68 | 0.29 | 0.05 | 0.54 | 0.26 | 0.04 | 0.87 | 0.34 |
| 14 | 0.07 | 0.89 | 0.28 | 0.10 | 0.86 | 0.28 | 0.04 | 0.96 | 0.30 |
| 15 | 0.02 | 0.25 | 0.24 | 0.00 | 0.00 | 0.17 | 0.04 | 0.26 | 0.24 |
| 16 | 0.04 | 0.03 | 0.22 | 0.04 | 0.00 | 0.09 | 0.04 | 0.05 | 0.28 |
| 17 | 0.04 | 0.44 | 0.12 | 0.01 | 0.12 | 0.11 | 0.07 | 0.49 | 0.22 |
| 18 | 0.05 | 0.04 | 0.12 | 0.06 | 0.08 | 0.06 | 0.05 | 0.00 | 0.13 |
| 19 | 0.05 | 0.00 | 0.06 | 0.09 | 0.00 | 0.06 | 0.00 | 0.00 | 0.10 |
| 20 | 0.04 | 0.00 | 0.06 | 0.07 | 0.00 | 0.06 | 0.00 | 0.00 | 0.08 |

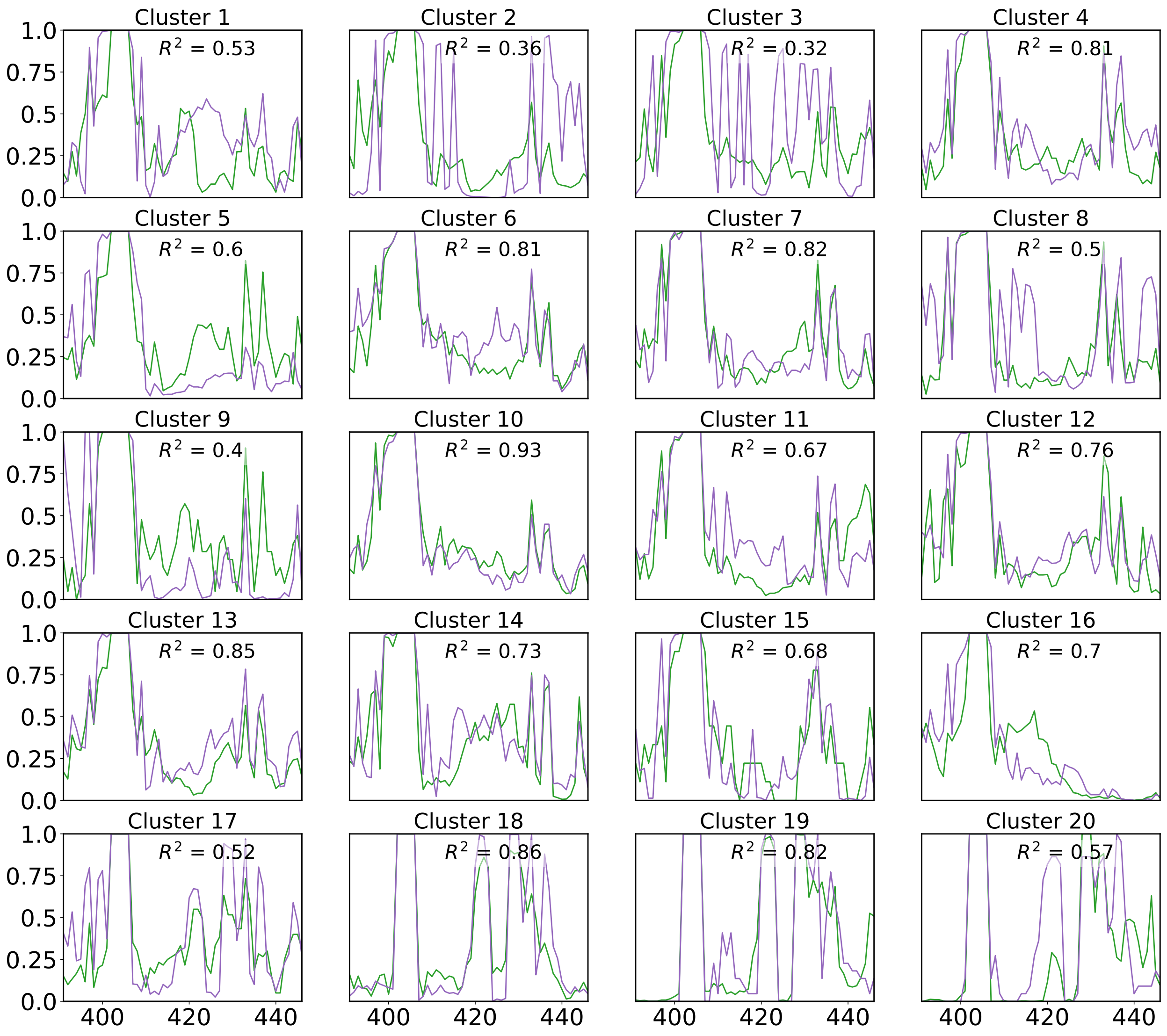


**Supplementary Figure 17: Intramolecular contact probabilities observed between covalently modified CYS404 residues and Tau-5$_{R2_R3}$ residues in covalent adduct conformational states identified by t-SNE clustering with $N$=20 clusters.** Populations of intramolecular contacts between Tau-5$_{R2_R3}$ residues and CYS404:EPI-002 (green) and Tau-5$_{R2_R3}$ residues and CYS404:EPI-7170 (purple) are shown for each cluster identified by t-SNE clustering with *perp*=300 and $N$=20 clusters. The coefficient of determination ($R^2$) of the populations of intramolecular contacts formed by CYS404:EPI-002 and CYS404:EPI-7170 are reported for each cluster.

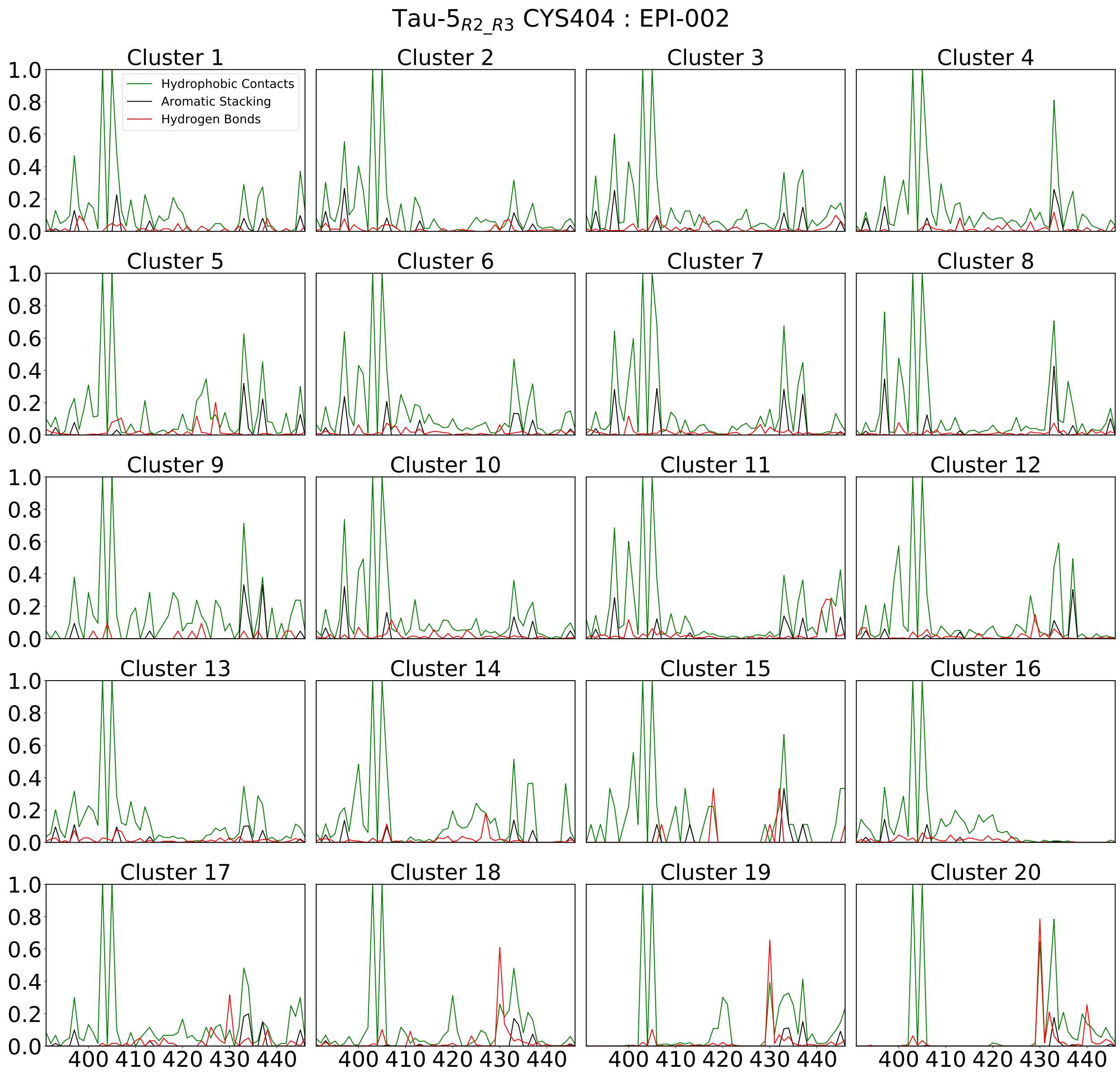


**Supplementary Figure 18: Populations of protein-ligand interactions in Tau-5$_{\mathbf{R2_R3}}$-CYS404:EPI-002 covalent adduct conformational states identified by t-SNE clustering with $\boldsymbol{N}$=20 clusters.** Populations of intramolecular interactions between CYS404:EPI-002 and Tau-5$_{R2_R3}$ residues in each cluster of the Tau-5$_{R2_R3}$-CYS404:EPI-002 ensemble.

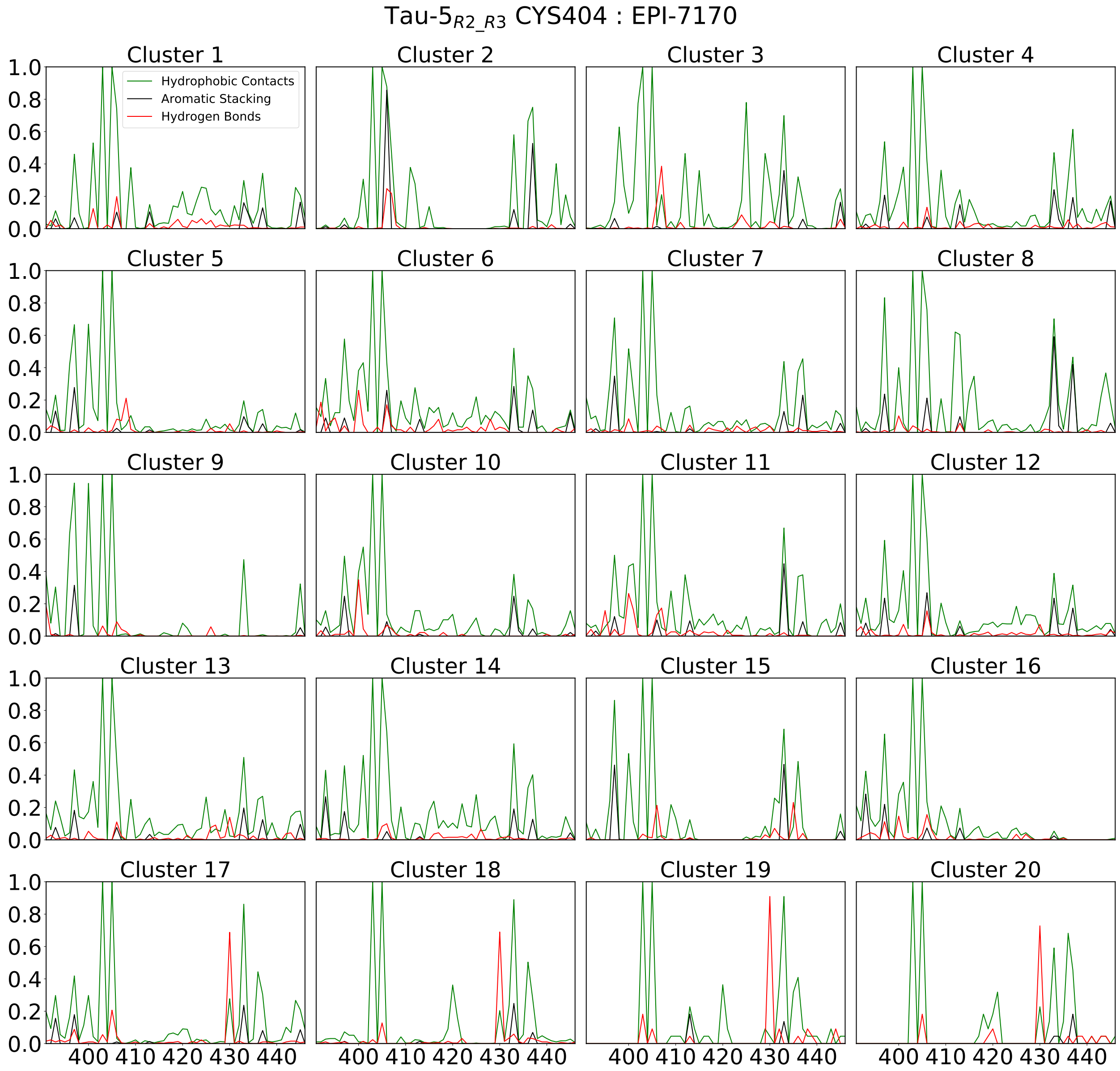


**Supplementary Figure 19: Populations of protein-ligand interactions in Tau-5$_{R2_R3}$-CYS404:EPI-7170 covalent adduct conformational states identified by t-SNE clustering with $N$=20 clusters.** Populations of intramolecular interactions between CYS404:EPI-7170 and Tau-5$_{R2_R3}$ residues in each cluster of the Tau-5$_{R2_R3}$-CYS404:EPI-7170 ensemble.

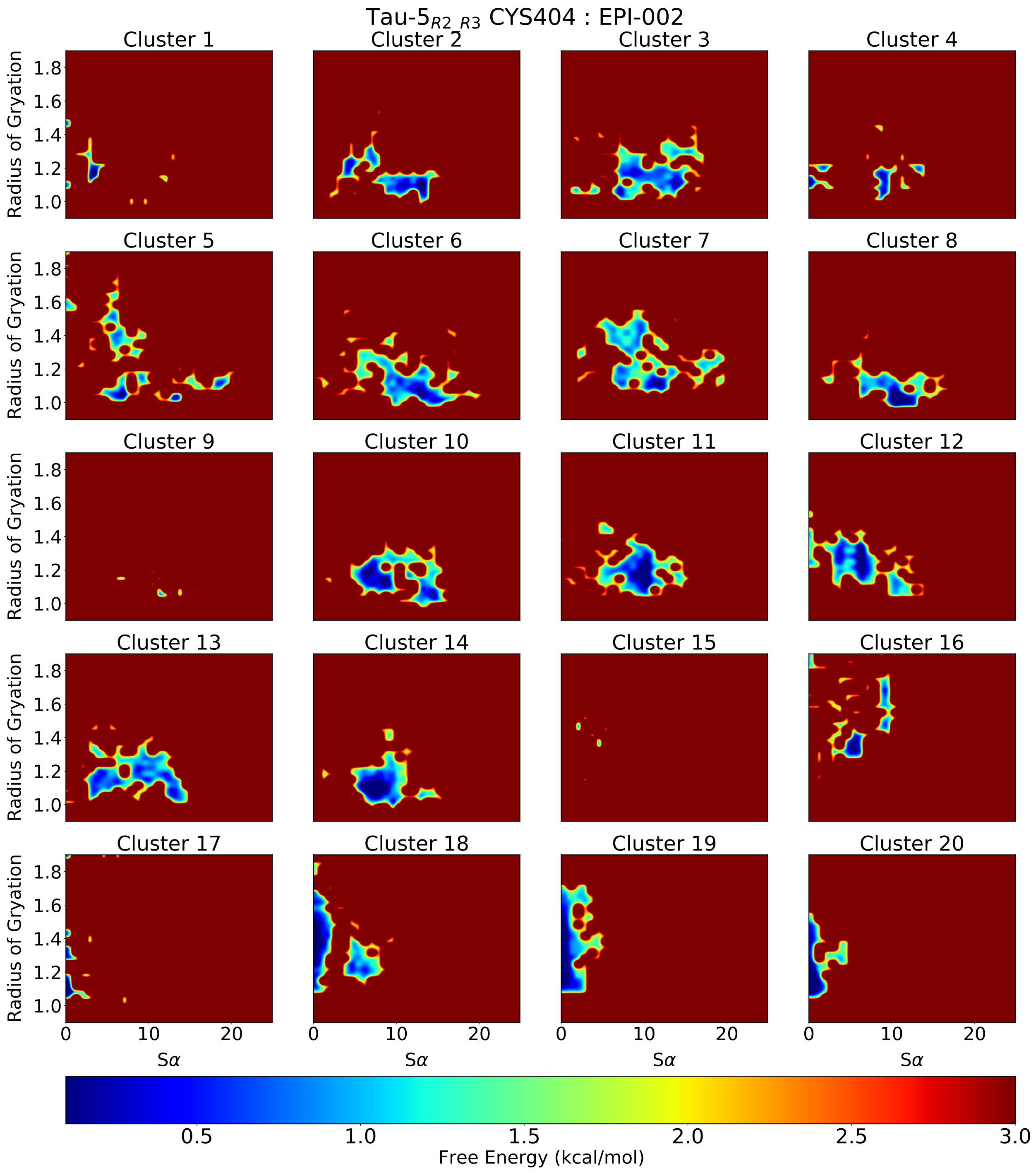


**Supplementary Figure 20: Free energy surfaces of Tau-5$_{\mathbf{R2_R3}}$-CYS404:EPI-002 covalent adduct conformational states identified by t-SNE clustering with $\boldsymbol{N}$=20 clusters.** Free energy surfaces as a function of the radius of gyration (reported in nm) and S$\alpha$ of Tau-5$_{\mathrm{R2_R3}}$ conformations for Tau-5$_{R2_R3}$-CYS404:EPI-002 covalent adduct conformational states identified by t-SNE clustering with $perp$ = 300 and $N$=20 clusters.

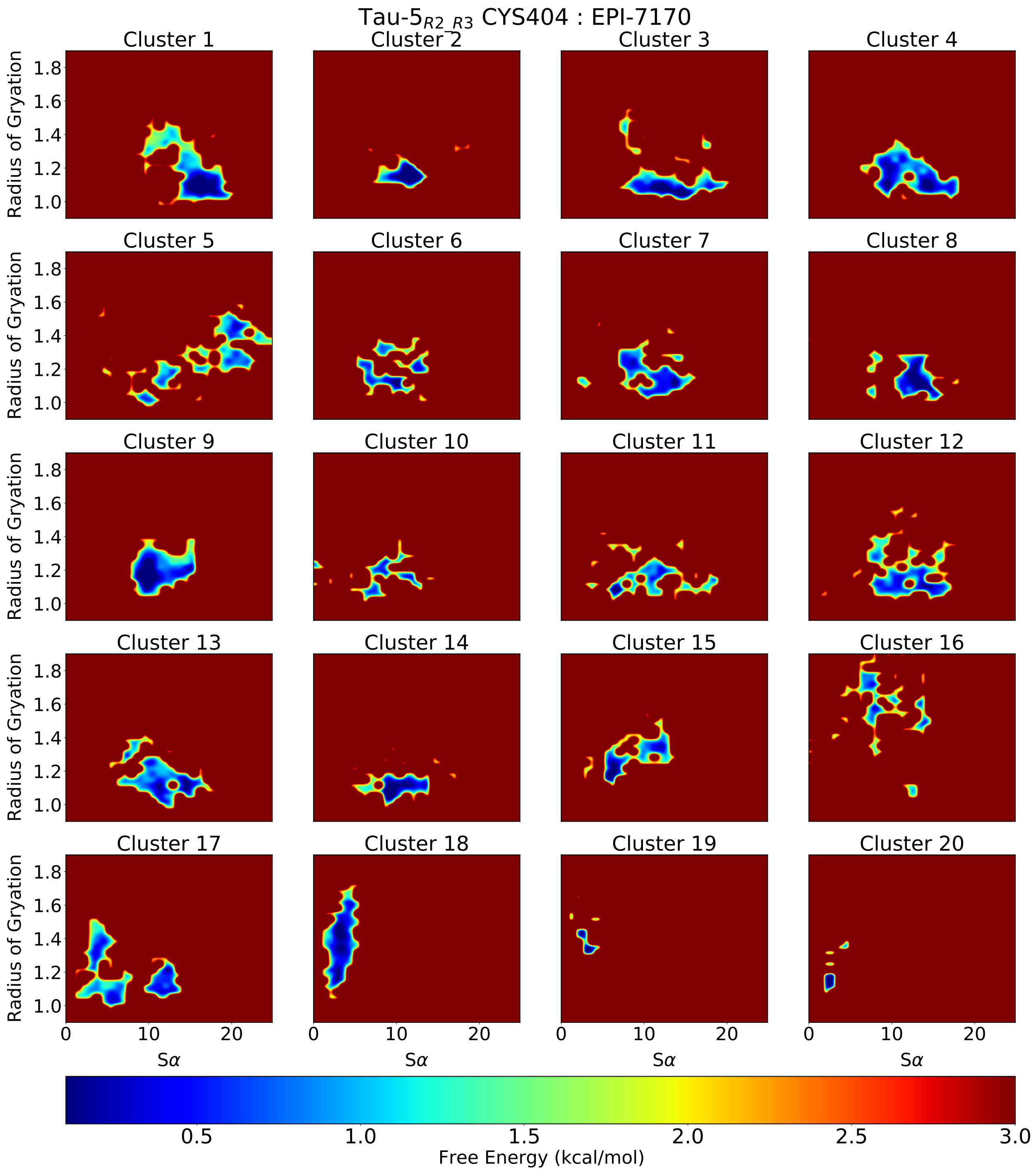


**Supplementary Figure 21: Free energy surfaces of Tau-5$_{\mathbf{R2_R3}}$-CYS404:EPI-7170 covalent adduct conformational states identified by t-SNE clustering with $N$=20 clusters.** Free energy surfaces as a function of the radius of gyration (reported in nm) and S$\alpha$ of Tau-5$_{\mathrm{R2_R3}}$ conformations for Tau-5$_{R2_R3}$-CYS404:EPI-7170 covalent adduct conformational states identified by t-SNE clustering with *perp* = 300 and $N$=20 clusters.

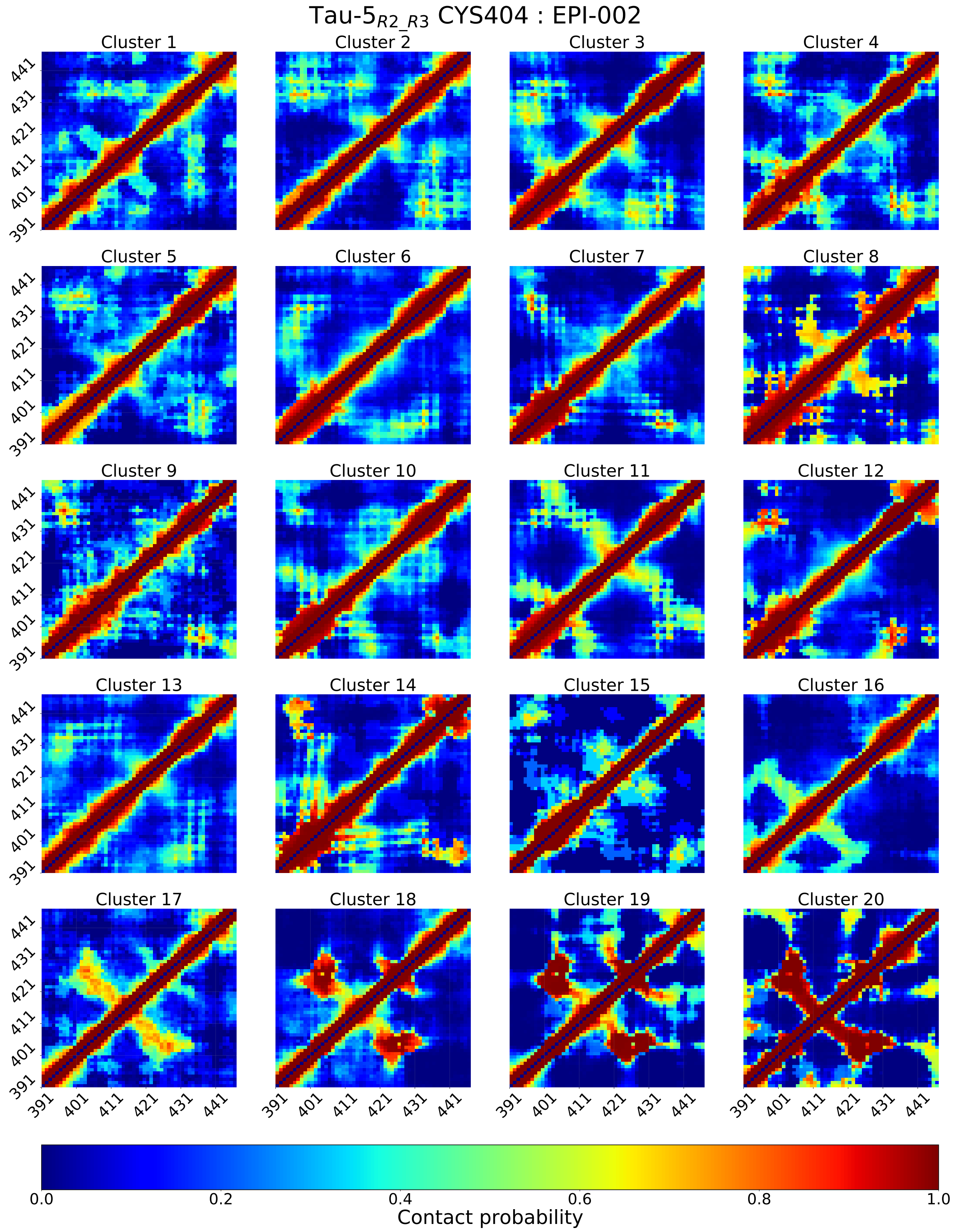


**Supplementary Figure 22: Intramolecular contact populations of Tau-5$_{R2_R3}$-CYS404:EPI-002 covalent adduct conformational states identified by t-SNE clustering with $N$=20 clusters.** Intramolecular contact populations of Tau-5$_{R2_R3}$-CYS404:EPI-002 covalent adduct conformational states identified by t-SNE clustering with $perp$ = 300 and $N$=20 clusters. Contacts between residues are defined using a distance cutoff of 12Å between closest heavy atoms.

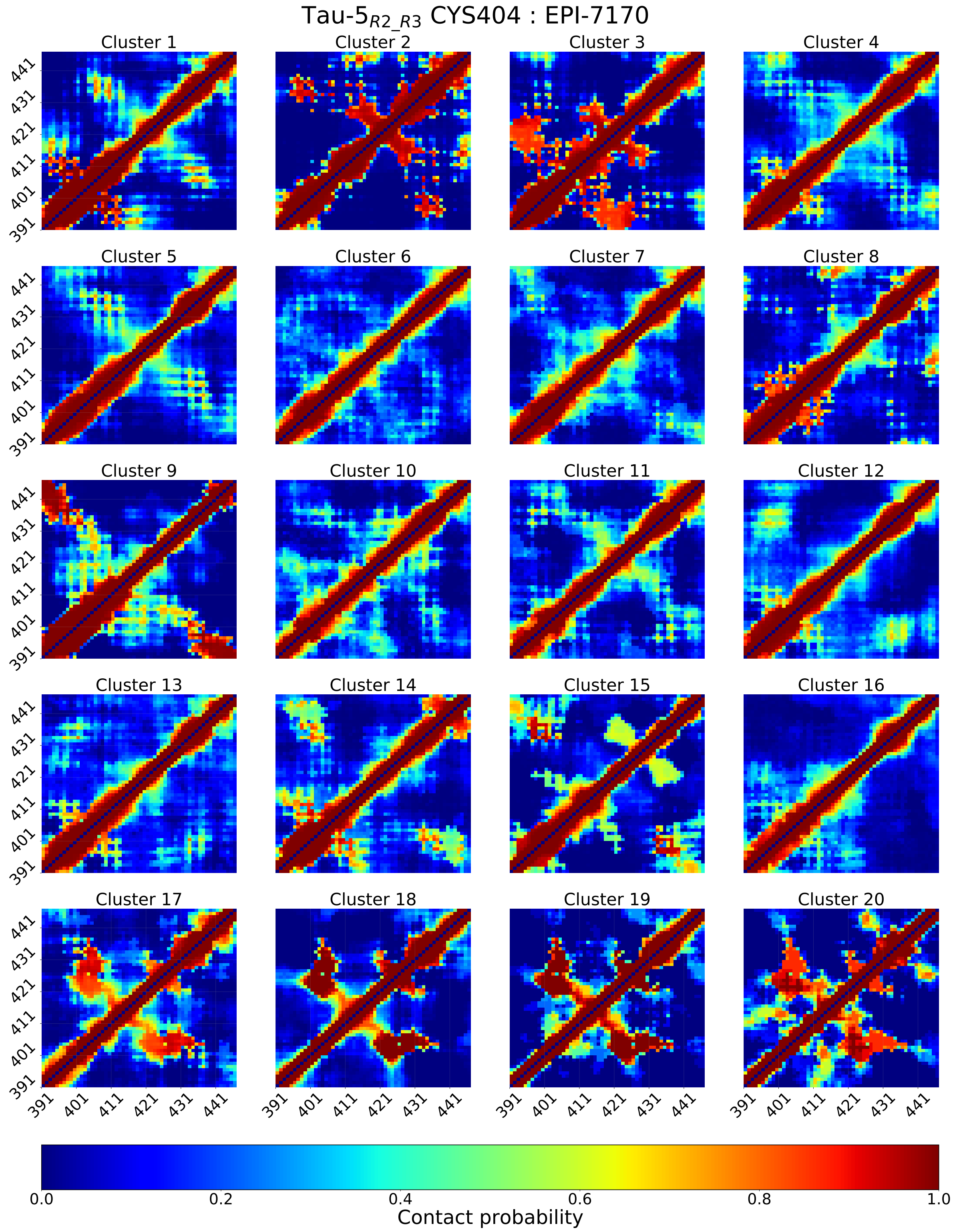


**Supplementary Figure 23: Intramolecular contact populations of Tau-5$_{R2_R3}$-CYS404:EPI-7170 covalent adduct conformational states identified by t-SNE clustering with $N$=20 clusters.** Intramolecular contact populations of Tau-5$_{R2_R3}$-CYS404:EPI-7170 covalent adduct conformational states identified by t-SNE clustering with $perp$ = 300 and $N$=20 clusters. Contacts between residues are defined using a distance cutoff of 12Å between closest heavy atoms.

**Supplementary Table 2:** Cluster population ($p$), bound fraction (BF), helical globule population ($p_{Glob}$) and helix fraction (HF) of clusters obtained from t-SNE clustering of a merged ensemble containing all frames from Tau-$5_{\mathrm{R2\_R3}}$:EPI-002 and Tau-$5_{\mathrm{R2\_R3}}$:EPI-7170 non-covalent binding simulations with $N$=18 clusters. We compare the properties of the clusters in the merged ensemble to the properties of the clustered conformations from the individual Tau-$5_{\mathrm{R2\_R3}}$:EPI-002 and Tau-$5_{\mathrm{R2\_R3}}$:EPI-7170 non-covalent binding simulations.

| Cluster | Merged Ensembles | | | | Tau-$5_{\mathrm{R2\_R3}}$:EPI-002 | | | | Tau-$5_{\mathrm{R2\_R3}}$:EPI-7170 | | | |
|---|---|---|---|---|---|---|---|---|---|---|---|---|
| | $p$ | BF | $p_{Glob}$ | HF | $p$ | BF | $p_{Glob}$ | HF | $p$ | BF | $p_{Glob}$ | HF |
| 1 | 0.08 | 0.74 | 0.89 | 0.38 | 0.07 | 0.67 | 0.90 | 0.36 | 0.09 | 0.80 | 0.88 | 0.40 |
| 2 | 0.05 | 0.66 | 0.78 | 0.36 | 0.04 | 0.60 | 0.70 | 0.35 | 0.06 | 0.70 | 0.82 | 0.37 |
| 3 | 0.07 | 0.68 | 0.90 | 0.35 | 0.04 | 0.57 | 0.81 | 0.32 | 0.10 | 0.72 | 0.93 | 0.37 |
| 4 | 0.06 | 0.60 | 0.68 | 0.32 | 0.04 | 0.69 | 0.78 | 0.38 | 0.07 | 0.55 | 0.63 | 0.29 |
| 5 | 0.04 | 0.53 | 0.89 | 0.32 | 0.07 | 0.50 | 0.95 | 0.31 | 0.02 | 0.65 | 0.71 | 0.34 |
| 6 | 0.07 | 0.66 | 0.61 | 0.31 | 0.02 | 0.33 | 0.62 | 0.25 | 0.11 | 0.73 | 0.61 | 0.32 |
| 7 | 0.06 | 0.53 | 0.69 | 0.29 | 0.08 | 0.41 | 0.70 | 0.32 | 0.05 | 0.70 | 0.68 | 0.26 |
| 8 | 0.06 | 0.59 | 0.70 | 0.28 | 0.06 | 0.39 | 0.47 | 0.23 | 0.07 | 0.75 | 0.88 | 0.32 |
| 9 | 0.05 | 0.47 | 0.69 | 0.27 | 0.09 | 0.40 | 0.64 | 0.25 | 0.02 | 0.76 | 0.89 | 0.34 |
| 10 | 0.05 | 0.68 | 0.18 | 0.26 | 0.02 | 0.62 | 0.22 | 0.16 | 0.07 | 0.69 | 0.17 | 0.28 |
| 11 | 0.05 | 0.65 | 0.66 | 0.25 | 0.03 | 0.45 | 0.29 | 0.16 | 0.06 | 0.73 | 0.80 | 0.28 |
| 12 | 0.05 | 0.48 | 0.00 | 0.25 | 0.03 | 0.36 | 0.00 | 0.12 | 0.08 | 0.51 | 0.00 | 0.28 |
| 13 | 0.06 | 0.26 | 0.44 | 0.23 | 0.11 | 0.21 | 0.43 | 0.22 | 0.01 | 0.75 | 0.56 | 0.27 |
| 14 | 0.03 | 0.71 | 0.71 | 0.22 | 0.00 | 0.65 | 0.61 | 0.17 | 0.05 | 0.72 | 0.72 | 0.22 |
| 15 | 0.05 | 0.56 | 0.19 | 0.19 | 0.06 | 0.37 | 0.07 | 0.16 | 0.05 | 0.76 | 0.14 | 0.27 |
| 16 | 0.05 | 0.52 | 0.09 | 0.15 | 0.05 | 0.57 | 0.03 | 0.12 | 0.05 | 0.49 | 0.14 | 0.14 |
| 17 | 0.05 | 0.49 | 0.08 | 0.14 | 0.06 | 0.47 | 0.05 | 0.08 | 0.04 | 0.51 | 0.13 | 0.16 |
| 18 | 0.06 | 0.30 | 0.00 | 0.03 | 0.13 | 0.30 | 0.00 | 0.03 | - | - | - | - |

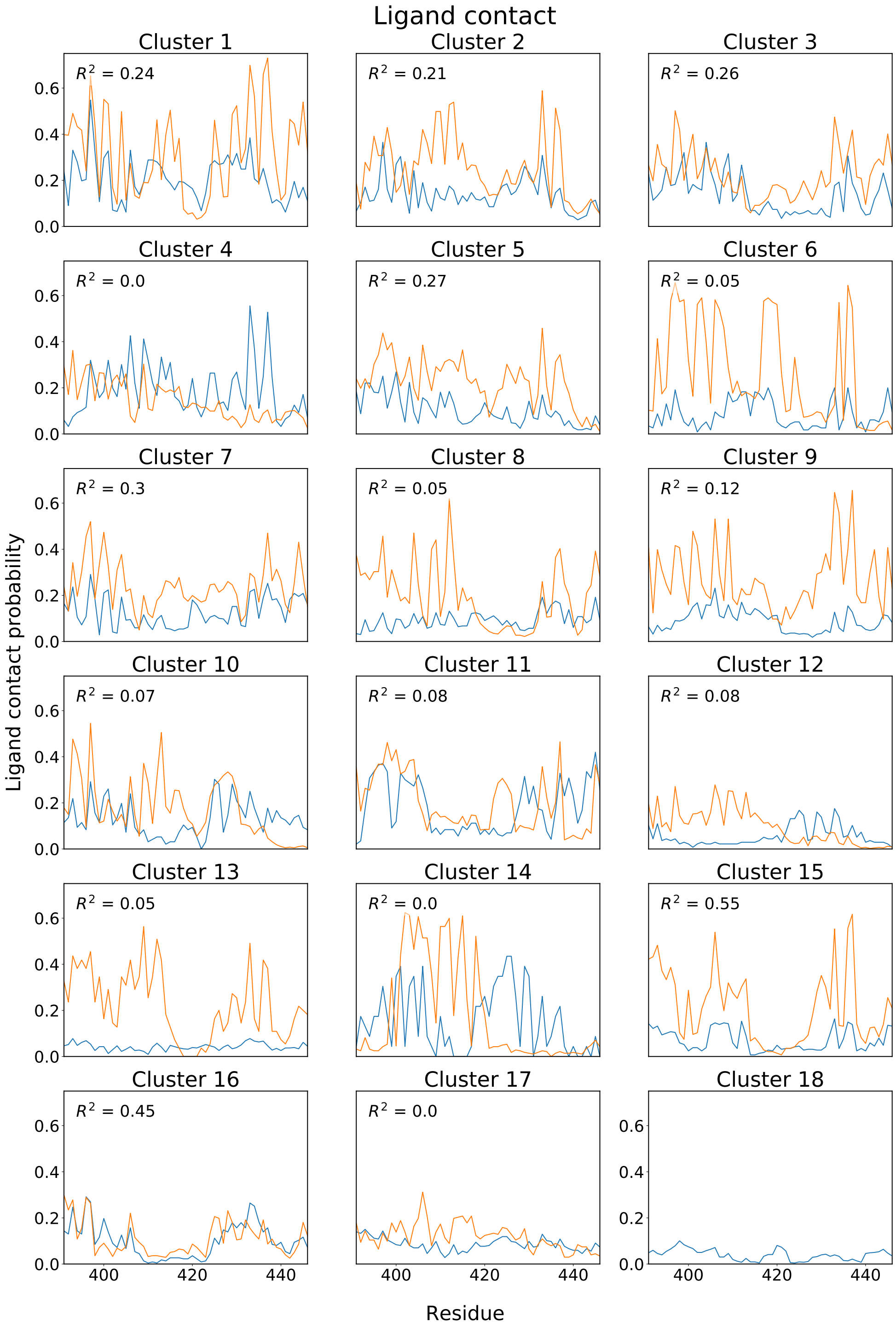


**Supplementary Figure 24: Populations of intermolecular protein-ligand contacts in Tau-5$_{\text{R2_R3}}$ conformational states identified by t-SNE clustering of non-covalent ligand binding simulations of EPI-002 and EPI-7170 with $N$=18 clusters.** Populations of intermolecular contacts between Tau-5$_{\text{R2_R3}}$ and EPI-002 (blue) and Tau-5$_{\text{R2_R3}}$ and EPI-7170 (orange) are shown for each cluster identified by t-SNE clustering with *perp*=200 and $N$=18 clusters. The coefficient of determination ($R^2$) of the populations of intramolecular contacts formed between Tau-5$_{\text{R2_R3}}$ and EPI-002 and Tau-5$_{\text{R2_R3}}$ and EPI-7170 are reported for each cluster.

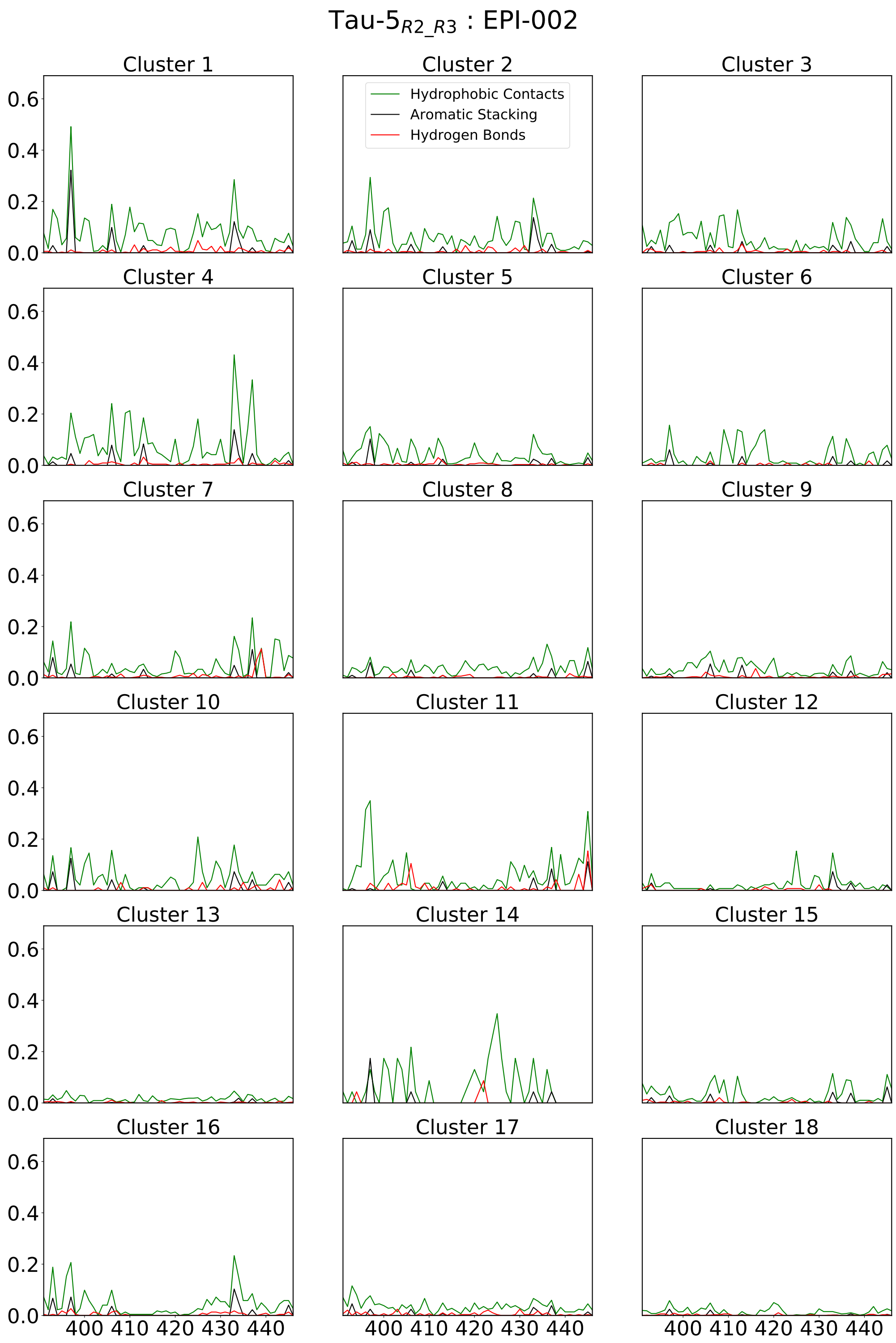


**Supplementary Figure 25: Populations of Tau-5$_{R2_R3}$:EPI-002 intermolecular protein-ligand interactions in Tau-5$_{R2_R3}$ conformational states identified by t-SNE clustering non-covalent ligand binding simulations of EPI-002 and EPI-7170 with $N$=18 clusters.** Populations of intermolecular interactions between EPI-002 and Tau-5$_{R2_R3}$ in each cluster.

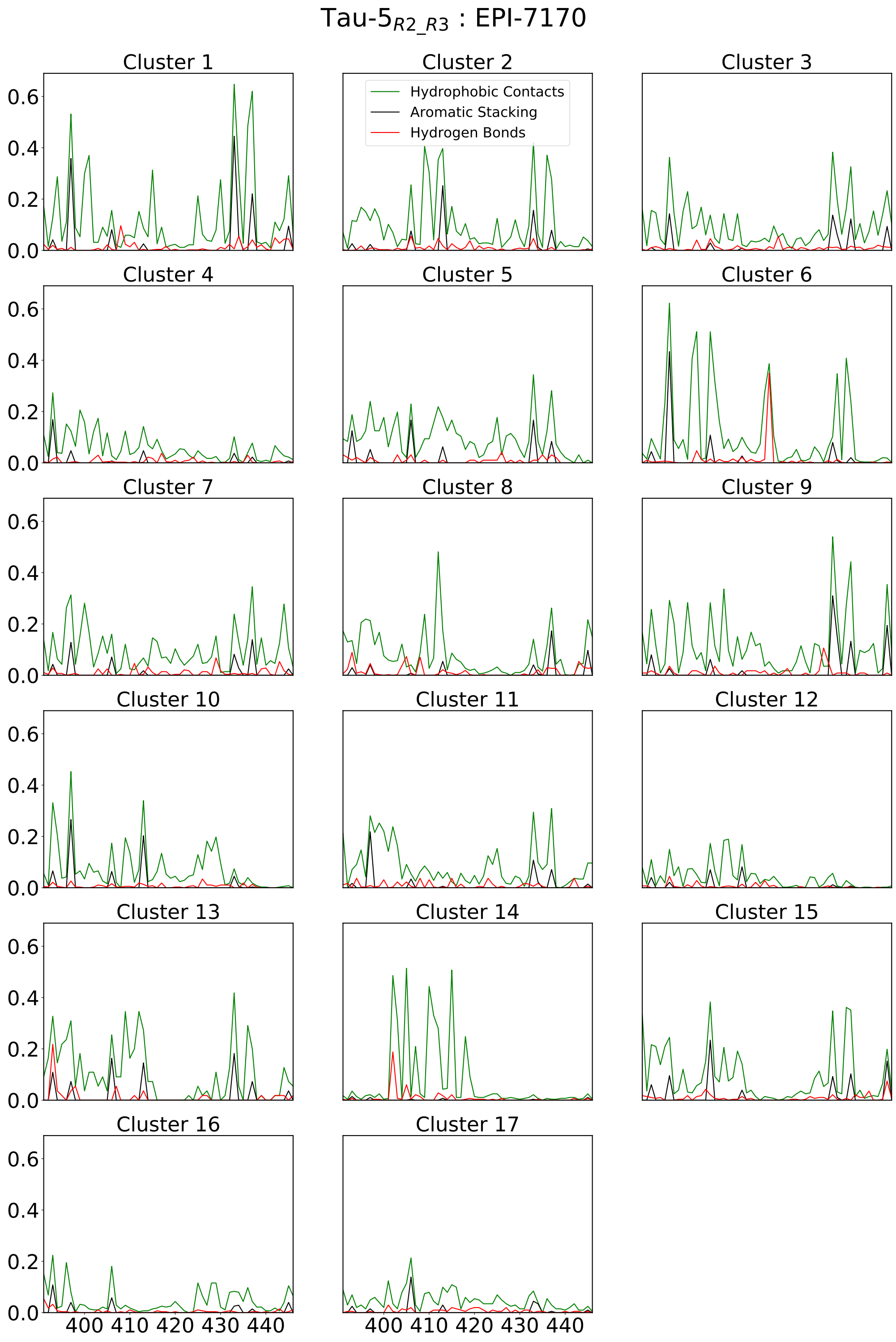


**Supplementary Figure 26: Populations of Tau-5$_{R2_R3}$:EPI-7170 intermolecular protein-ligand interactions in Tau-5$_{R2_R3}$ conformational states identified by t-SNE clustering non-covalent ligand binding simulations of EPI-002 and EPI-7170 with $N$=18 clusters.** Populations of intermolecular interactions between EPI-7170 and Tau-5$_{R2_R3}$ in each cluster. The interaction plot of cluster 18 is excluded because only one frame from this ensemble was assigned to the this cluster.

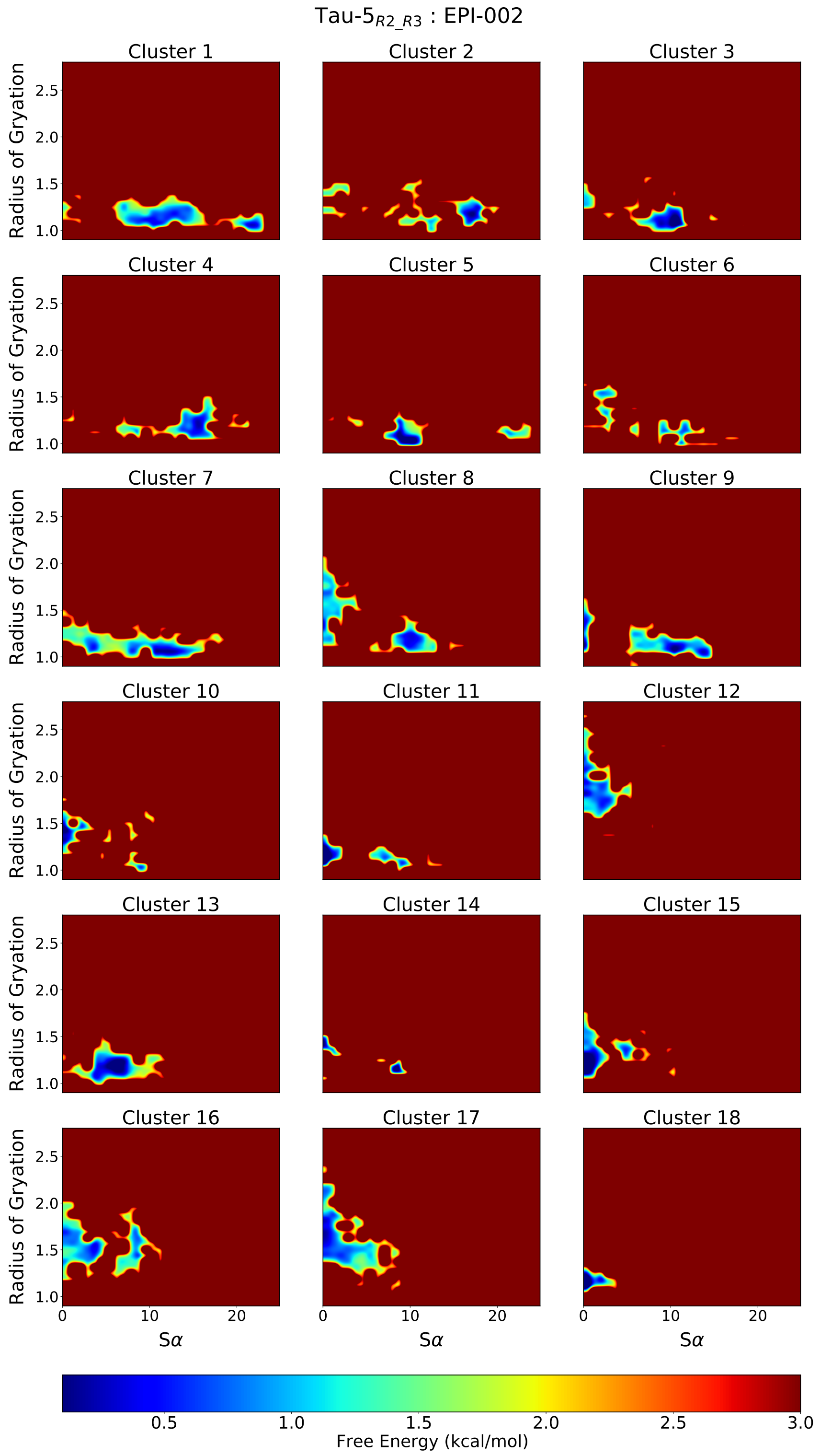


**Supplementary Figure 27: Free energy surfaces of conformational states from a non-covalent ligand binding simulation of Tau-5$_{\mathbf{R2_R3}}$ and EPI-002 identified by t-SNE clustering with $N$=18 clusters.** Free energy surfaces as a function of the radius of gyration (reported in nm) and S$\alpha$ of Tau-5$_{\mathrm{R2_R3}}$ conformations of conformational states from a non-covalent ligand-binding simulation of Tau-5$_{\mathrm{R2_R3}}$ in the presence of EPI-002 identified by t-SNE clustering with $perp$ = 200 and $N$=18 clusters.

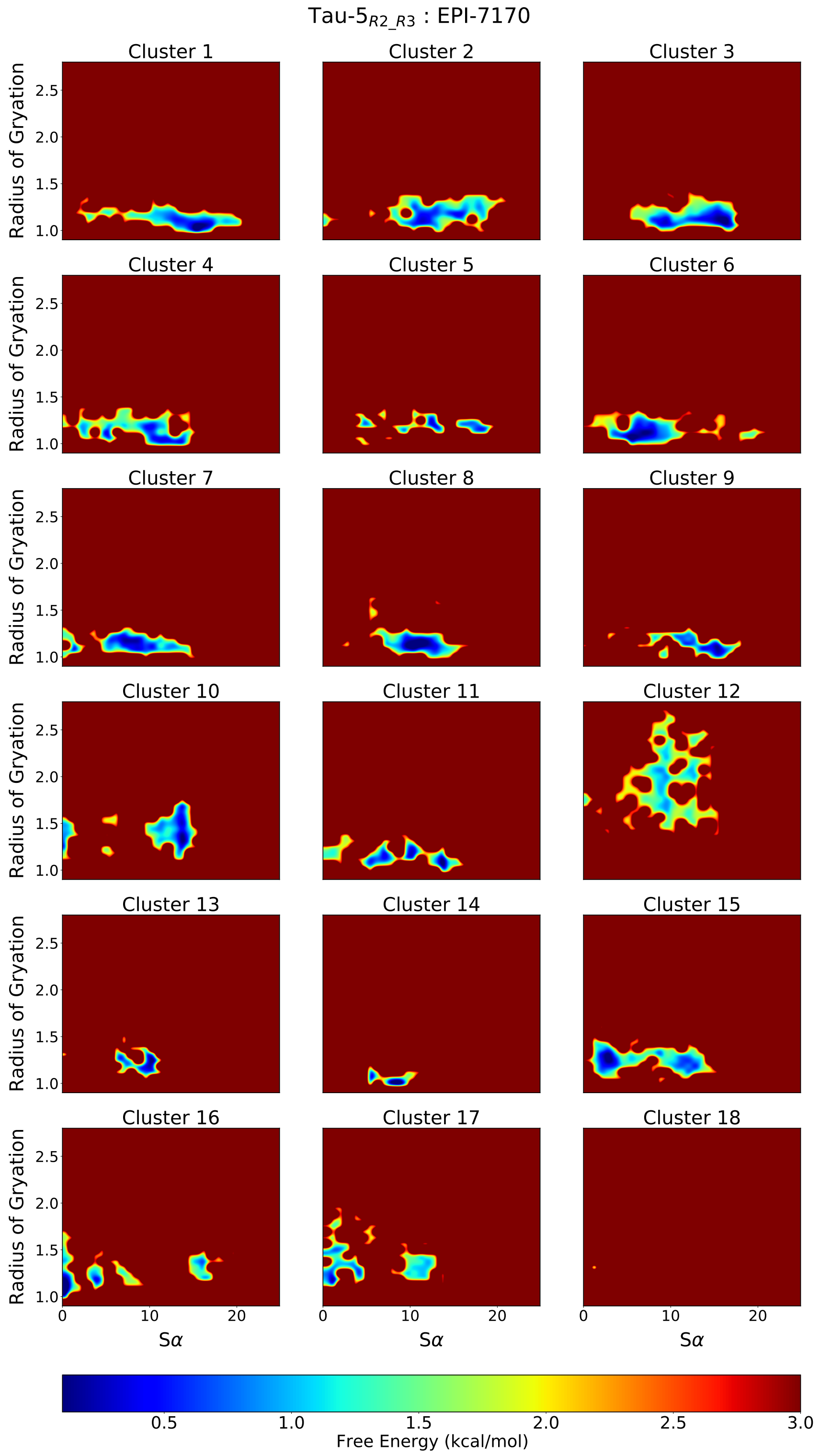


**Supplementary Figure 28: Free energy surfaces of conformational states from a non-covalent ligand binding simulation of Tau-5$_{\mathrm{R2_R3}}$ and EPI-7170 identified by t-SNE clustering with $N$=18 clusters.** Free energy surfaces as a function of the radius of gyration (reported in nm) and S$\alpha$ of Tau-5$_{\mathrm{R2_R3}}$ conformations of conformational states from a non-covalent ligand-binding simulation of Tau-5$_{\mathrm{R2_R3}}$ in the presence of EPI-7170 identified by t-SNE clustering with $perp$ = 200 and $N$=18 clusters.

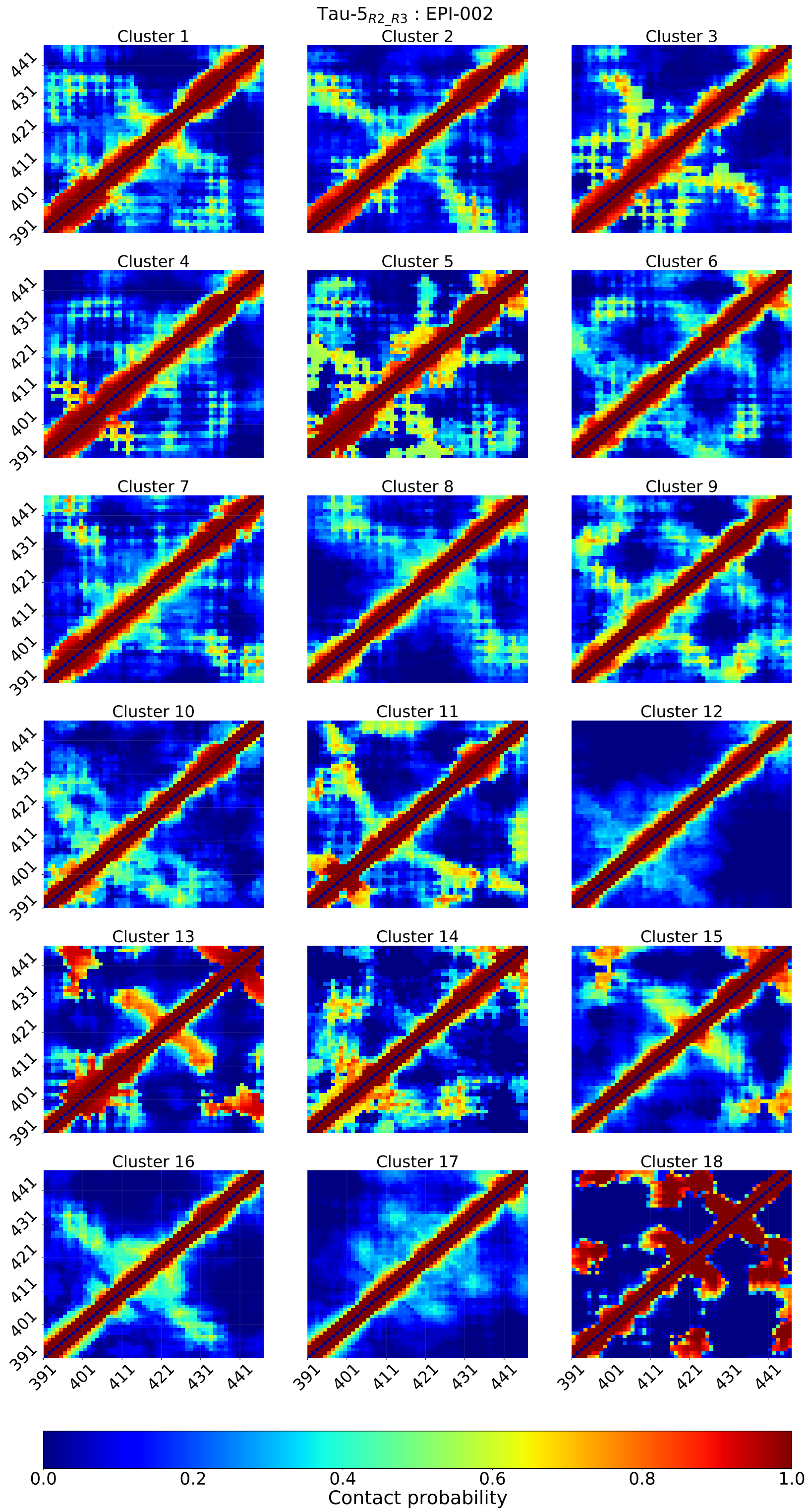


**Supplementary Figure 29: Intramolecular contact populations of Tau-5$_{\mathbf{R2_R3}}$ conformational states identified by t-SNE clustering of a non-covalent ligand-binding simulation of EPI-002 with $N$=18 clusters.** Intramolecular contact populations of Tau-5$_{\mathrm{R2_R3}}$ conformational states identified from a non-covalent EPI-002 binding simulation by t-SNE clustering with $perp$ = 200 and $N$=18 clusters. We note only one frame from this ensemble was assigned to cluster 18. Contacts between residues are defined using a distance cutoff of 12Å between closest heavy atoms.

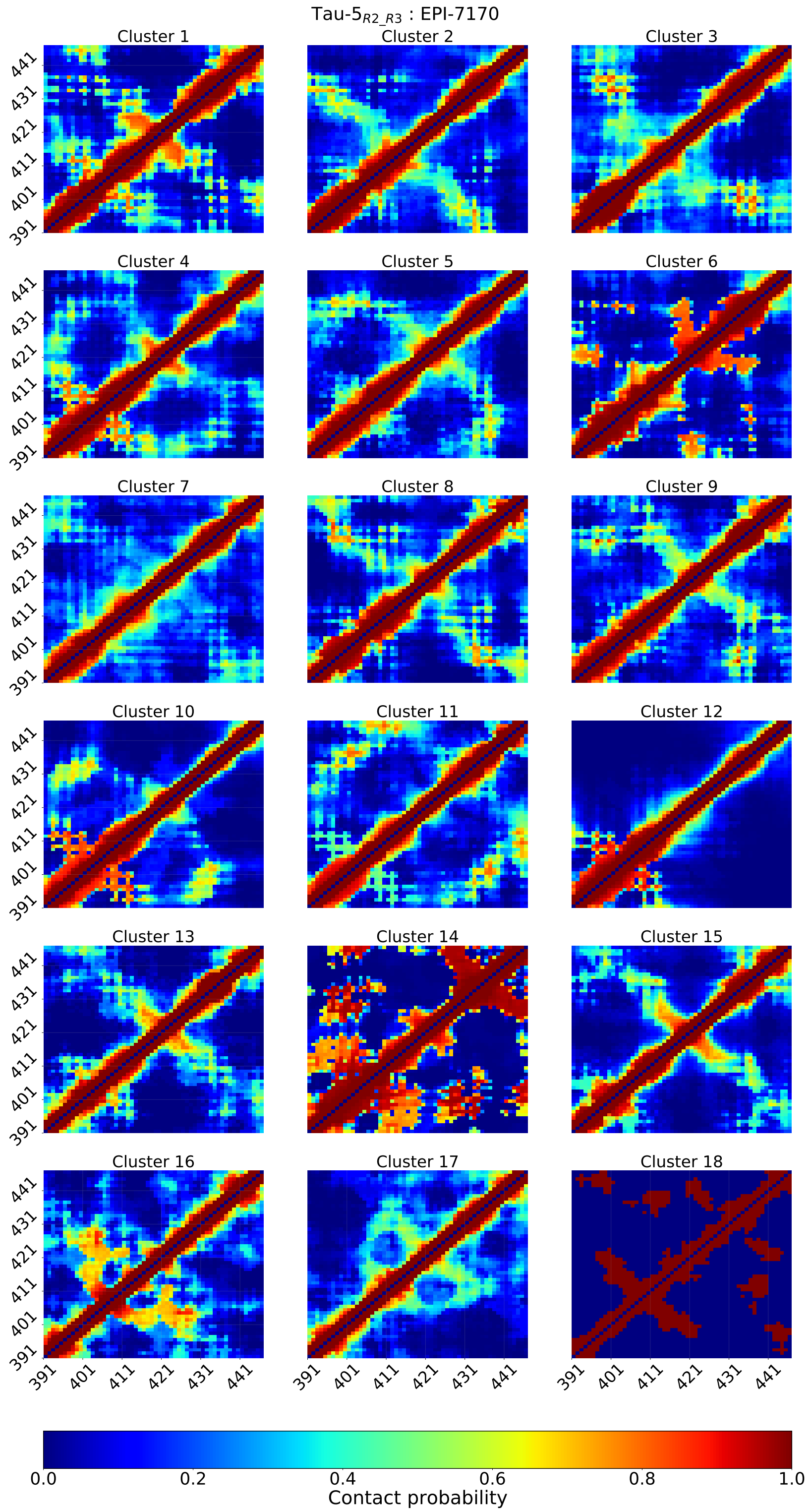


**Supplementary Figure 30: Intramolecular contact populations of Tau-5$_{\mathbf{R2_R3}}$ conformational states identified by t-SNE clustering of a non-covalent ligand-binding simulation of EPI-7170 with $N$=18 clusters.** Intramolecular contact populations of Tau-5$_{\mathrm{R2_R3}}$ conformational states identified from a non-covalent EPI-7170 binding simulation by t-SNE clustering with $perp$ = 200 and $N$=18 clusters. Contacts between residues are defined using a distance cutoff of 12Å between closest heavy atoms.

CYS:EPI-002 (CYE2)

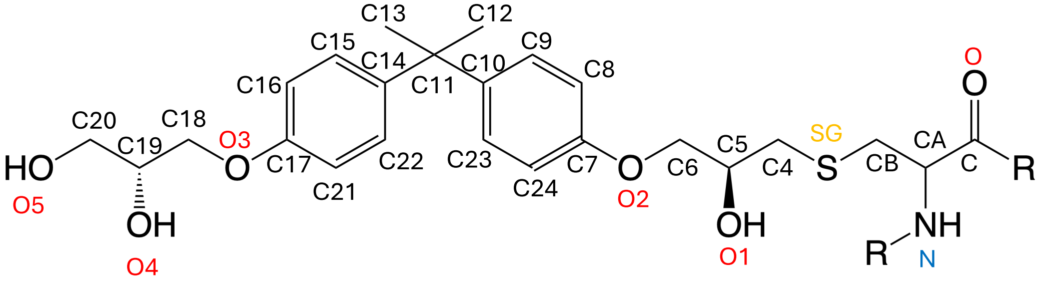


CYS:EPI-7170 (CYE7)

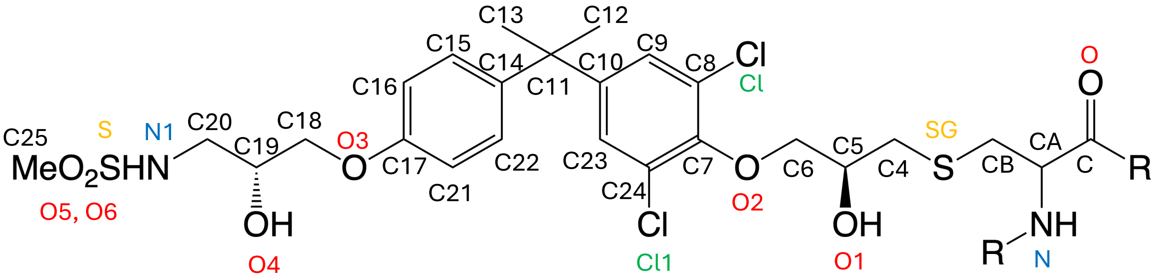


**Supplementary Figure 31: Atom name definitions for CYS:EPI-002 (CYE2) and CYS:EPI-7170 (CYE7) covalently modified cysteine residue force field parameters.** Atom names correspond to the force field parameters described in SI tables 3-9.

**Supplementary Table 3: Partial charge force field parameters for the modified cysteine residue CYS:EPI-002 (CYE2).** Atom names, atom types, atom numbers, partial charges and bond list for atoms contained in the modified cysteine residue CYS:EPI-002 (CYE2).

| [ CYE2 ] | | | | | | | | | | | |
|---|---|---|---|---|---|---|---|---|---|---|---|
| [ atoms ] | | | | | | | | [ bonds ] | | | |
| N | N | -0.59637 | 1 | O5 | OH | -0.59230 | 34 | -C | N | C12 | H17 |
| H | H | 0.34653 | 2 | C21 | CA | -0.17795 | 35 | N | H | C12 | H18 |
| CA | CT | 0.05603 | 3 | C22 | CA | -0.09146 | 36 | N | CA | C13 | H19 |
| HA | H1 | 0.09526 | 4 | C23 | CA | -0.09245 | 37 | CA | CB | C13 | H20 |
| CB | CT | -0.03211 | 5 | C24 | CA | -0.17795 | 38 | CA | HA | C13 | H21 |
| HB1 | H1 | 0.09978 | 6 | H9 | H1 | 0.10431 | 39 | CA | C | C14 | C15 |
| HB2 | H1 | 0.09978 | 7 | H10 | H1 | 0.10431 | 40 | CB | SG | C14 | C22 |
| SG | S | -0.29345 | 8 | H11 | HO | 0.42146 | 41 | CB | HB1 | C15 | C16 |
| C | C | 0.64386 | 9 | H12 | H1 | 0.06659 | 42 | CB | HB2 | C15 | H22 |
| O | O | -0.64528 | 10 | H13 | H1 | 0.06659 | 43 | SG | C4 | C16 | C17 |
| C4 | CT | -0.02315 | 11 | H14 | HA | 0.14786 | 44 | C4 | C5 | C16 | H23 |
| C5 | CT | 0.14896 | 12 | H15 | HA | 0.13831 | 45 | C4 | H9 | C17 | O3 |
| H1 | H1 | 0.06407 | 13 | H16 | HC | 0.04027 | 46 | C4 | H10 | C17 | C21 |
| O1 | OH | -0.60025 | 14 | H17 | HC | 0.04027 | 47 | C5 | O1 | O3 | C18 |
| C6 | CT | 0.08390 | 15 | H18 | HC | 0.04027 | 48 | C5 | C6 | C18 | C19 |
| O2 | OS | -0.34685 | 16 | H19 | HC | 0.04027 | 49 | C5 | H1 | C18 | H24 |
| C7 | CA | 0.12483 | 17 | H20 | HC | 0.04027 | 50 | O1 | H11 | C18 | H25 |
| C8 | CA | -0.17795 | 18 | H21 | HC | 0.04027 | 51 | C6 | O2 | C19 | O4 |
| C9 | CA | -0.09245 | 19 | H22 | HA | 0.13931 | 52 | C6 | H12 | C19 | C20 |
| C10 | CA | -0.10568 | 20 | H23 | HA | 0.14334 | 53 | C6 | H13 | C19 | H2 |
| C11 | CT | 0.04989 | 21 | H24 | H1 | 0.05955 | 54 | O2 | C7 | O4 | H26 |
| C12 | CT | -0.08559 | 22 | H25 | H1 | 0.05955 | 55 | C7 | C8 | C20 | O5 |
| C13 | CT | -0.08559 | 23 | H26 | HO | 0.42347 | 56 | C7 | C24 | C20 | H27 |
| C14 | CA | -0.10170 | 24 | H27 | H1 | 0.06407 | 57 | C8 | C9 | C20 | H28 |
| C15 | CA | -0.09146 | 25 | H28 | H1 | 0.06407 | 58 | C8 | H14 | O5 | H29 |
| C16 | CA | -0.17795 | 26 | H29 | HO | 0.40336 | 59 | C9 | C10 | C21 | C22 |
| C17 | CA | 0.12483 | 27 | H30 | HA | 0.14334 | 60 | C9 | H15 | C21 | H30 |
| O3 | OS | -0.34486 | 28 | H31 | HA | 0.13931 | 61 | C10 | C11 | C22 | H31 |
| C18 | CT | 0.09093 | 29 | H32 | HA | 0.13831 | 62 | C10 | C23 | C23 | C24 |
| C19 | CT | 0.10672 | 30 | H33 | HA | 0.14786 | 63 | C11 | C12 | C23 | H32 |
| H2 | H1 | 0.04798 | 31 | | | | | C11 | C13 | C24 | H33 |
| O4 | OH | -0.59528 | 32 | | | | | C11 | C14 | C | O |
| C20 | CT | 0.12815 | 33 | | | | | C12 | H16 | | |

**Supplementary Table 4: Dihedral force field parameters of the modified cysteine residue CYS:EPI-002 (CYE2).** CYE2 dihedral parameters are applied using the improper dihedral section of the GROMACS CYE2 amino acid template.

| [ impropers ] | | | | | | |
|---|---|---|---|---|---|---|
| -C | N | CA | C | 90.0 | 1.67360 | 2 |
| N | CA | C | +N | 160.888 | 2.71960 | 1 |
| N | CA | C | +N | 90.0 | -1.0669 | 1 |
| N | CA | C | +N | 90.0 | 0.3138 | 2 |
| N | CA | C | +N | 90.0 | 0.2385 | 3 |
| N | CA | C | +N | 90.0 | 0.1046 | 4 |
| N | CA | C | +N | 90.0 | -0.0460 | 5 |
| -C | CA | N | H | | | |
| CA | +N | C | O | | | |
| C7 | C9 | C8 | H14 | 180.00 | 4.60240 | 2 |
| C7 | C23 | C24 | H33 | 180.00 | 4.60240 | 2 |
| C8 | C10 | C9 | H15 | 180.00 | 4.60240 | 2 |
| C8 | C24 | C7 | O2 | 180.00 | 4.60240 | 2 |
| C9 | C23 | C10 | C11 | 180.00 | 4.60240 | 2 |
| C10 | C24 | C23 | H32 | 180.00 | 4.60240 | 2 |
| C14 | C16 | C15 | H22 | 180.00 | 4.60240 | 2 |
| C14 | C21 | C22 | H31 | 180.00 | 4.60240 | 2 |
| C15 | C17 | C16 | H23 | 180.00 | 4.60240 | 2 |
| C15 | C22 | C14 | C11 | 180.00 | 4.60240 | 2 |
| C16 | C21 | C17 | O3 | 180.00 | 4.60240 | 2 |
| C17 | C22 | C21 | H30 | 180.00 | 4.60240 | 2 |

**Supplementary Table 5: Partial charge force field parameters for the modified cysteine residue CYS:EPI-7170 (CYE7).** Atom names, atom types, atom numbers, partial charges and bond list for atoms contained in the modified cysteine residue CYS:EPI-7170 (CYE7).

| [ CYE7 ] | | | | | | | | | | | |
|---|---|---|---|---|---|---|---|---|---|---|---|
| [ atoms ] | | | | | | | | [ bonds ] | | | |
| N | N | -0.59637 | 1 | C24 | CA | 0.03303 | 37 | -C | N | C13 | H20 |
| H | H | 0.34653 | 2 | H9 | H1 | 0.10431 | 38 | N | H | C13 | H21 |
| CA | CT | 0.05603 | 3 | H10 | H1 | 0.10431 | 39 | N | CA | C14 | C15 |
| HA | H1 | 0.09526 | 4 | H11 | HO | 0.42146 | 40 | CA | CB | C14 | C22 |
| CB | CT | -0.03211 | 5 | H12 | H1 | 0.06659 | 41 | CA | HA | C15 | C16 |
| HB1 | H1 | 0.09978 | 6 | H13 | H1 | 0.06659 | 42 | CA | C | C15 | H22 |
| HB2 | H1 | 0.09978 | 7 | CL1 | Cl | -0.05300 | 43 | CB | SG | C16 | C17 |
| SG | S | -0.29345 | 8 | H15 | HA | 0.13831 | 44 | CB | HB1 | C16 | H23 |
| C | C | 0.64386 | 9 | H16 | HC | 0.04027 | 45 | CB | HB2 | C17 | O3 |
| O | O | -0.64528 | 10 | H17 | HC | 0.04027 | 46 | SG | C4 | C17 | C21 |
| C4 | CT | -0.02315 | 11 | H18 | HC | 0.04027 | 47 | C4 | C5 | O3 | C18 |
| C5 | CT | 0.14896 | 12 | H19 | HC | 0.04027 | 48 | C4 | H9 | C18 | C19 |
| H1 | H1 | 0.06407 | 13 | H20 | HC | 0.04027 | 49 | C4 | H10 | C18 | H24 |
| O1 | OH | -0.60025 | 14 | H21 | HC | 0.04027 | 50 | C5 | O1 | C18 | H25 |
| C6 | CT | 0.08390 | 15 | H22 | HA | 0.13931 | 51 | C5 | C6 | C19 | O4 |
| O2 | OS | -0.34685 | 16 | H23 | HA | 0.14334 | 52 | C5 | H1 | C19 | C20 |
| C7 | CA | 0.10655 | 17 | H24 | H1 | 0.06698 | 53 | O1 | H11 | C19 | H2 |
| C8 | CA | 0.03303 | 18 | H25 | H1 | 0.06698 | 54 | C6 | O2 | O4 | H26 |
| C9 | CA | -0.09343 | 19 | H26 | HO | 0.42347 | 55 | C6 | H12 | C20 | H27 |
| C10 | CA | -0.10568 | 20 | H27 | H1 | 0.06407 | 56 | C6 | H13 | C20 | H28 |
| C11 | CT | 0.04989 | 21 | H28 | H1 | 0.06407 | 57 | O2 | C7 | C21 | C22 |
| C12 | CT | -0.08559 | 22 | H30 | HA | 0.14334 | 58 | C7 | C8 | C21 | H30 |
| C13 | CT | -0.08559 | 23 | H31 | HA | 0.13931 | 59 | C7 | C24 | C22 | H31 |
| C14 | CA | -0.10170 | 24 | H32 | HA | 0.13831 | 60 | C8 | C9 | C23 | C24 |
| C15 | CA | -0.09146 | 25 | CL2 | Cl | -0.05300 | 61 | C8 | Cl1 | C23 | H32 |
| C16 | CA | -0.17795 | 26 | N1 | NT | -0.92124 | 62 | C9 | C10 | C24 | Cl2 |
| C17 | CA | 0.12483 | 27 | H29 | H | 0.45770 | 63 | C9 | H15 | C20 | N1 |
| O3 | OS | -0.34486 | 28 | S | SO | 1.47504 | 64 | C10 | C11 | N1 | S |
| C18 | CT | 0.08274 | 29 | O6 | O | -0.65356 | 65 | C10 | C23 | N1 | H29 |
| C19 | CT | 0.12162 | 30 | O5 | O | -0.65356 | 66 | C11 | C12 | S | C25 |
| H2 | H1 | 0.05694 | 31 | C25 | CT | -0.36377 | 67 | C11 | C13 | S | O6 |
| O4 | OH | -0.60228 | 32 | H34 | H1 | 0.11510 | 68 | C11 | C14 | S | O5 |
| C20 | CT | 0.22976 | 33 | H35 | H1 | 0.11510 | 69 | C12 | H16 | C25 | H34 |
| C21 | CA | -0.17795 | 34 | H36 | H1 | 0.11510 | 70 | C12 | H17 | C25 | H35 |
| C22 | CA | -0.09146 | 35 | | | | | C12 | H18 | C25 | H36 |
| C23 | CA | -0.09343 | 36 | | | | | C13 | H19 | C | O |

**Supplementary Table 6: Dihedral force field parameters of the modified cysteine residue CYS:EPI-7170 (CYE7).** CYE7 dihedral parameters are applied using the improper dihedral section of the GROMACS CYE7 amino acid template.

| [ impropers ] | | | | | | |
|---|---|---|---|---|---|---|
| -C | N | CA | C | 90.0 | 1.67360 | 2 |
| N | CA | C | +N | 160.888 | 2.71960 | 1 |
| N | CA | C | +N | 90.0 | -1.0669 | 1 |
| N | CA | C | +N | 90.0 | 0.3138 | 2 |
| N | CA | C | +N | 90.0 | 0.2385 | 3 |
| N | CA | C | +N | 90.0 | 0.1046 | 4 |
| N | CA | C | +N | 90.0 | -0.0460 | 5 |
| -C | CA | N | H | | | |
| CA | +N | C | O | | | |
| C7 | C9 | C8 | Cl1 | 180.00 | 4.60240 | 2 |
| C7 | C23 | C24 | Cl2 | 180.00 | 4.60240 | 2 |
| C8 | C10 | C9 | H15 | 180.00 | 4.60240 | 2 |
| C8 | C24 | C7 | O2 | 180.00 | 4.60240 | 2 |
| C9 | C23 | C10 | C11 | 180.00 | 4.60240 | 2 |
| C10 | C24 | C23 | H32 | 180.00 | 4.60240 | 2 |
| C14 | C16 | C15 | H22 | 180.00 | 4.60240 | 2 |
| C14 | C21 | C22 | H31 | 180.00 | 4.60240 | 2 |
| C15 | C17 | C16 | H23 | 180.00 | 4.60240 | 2 |
| C15 | C22 | C14 | C11 | 180.00 | 4.60240 | 2 |
| C16 | C21 | C17 | O3 | 180.00 | 4.60240 | 2 |
| C17 | C22 | C21 | H30 | 180.00 | 4.60240 | 2 |

**Supplementary Table 7: Added force field bond length parameters for modified cysteine residues CYE2 and CYE7.** These parameters were added to the existing bond length parameters in the a99SB-*disp* force field.

| [ bondtypes ] | | | | | |
|---|---|---|---|---|---|
| i | j | func | b0 | kb | |
| OS | CA | 1 | 0.13696 | 315140.0 | ; Amber99Sb-disp |
| CT | HO | 1 | 0.10969 | 276650.0 | ; Amber99Sb-disp |
| NT | SO | 1 | 0.16720 | 265350.0 | ; Amber99Sb-disp |
| NT | H | 1 | 0.10100 | 363170.0 | ; Amber99Sb-disp |
| SO | CT | 1 | 0.18080 | 195390.0 | ; Amber99Sb-disp |
| SO | O | 1 | 0.14530 | 429030.0 | ; Amber99Sb-disp |
| CT | NT | 1 | 0.14710 | 307110.0 | ; Amber99Sb-disp |

**Supplementary Table 8: Added force field bond angle parameters for modified cysteine residues CYE2 and CYE7.** These parameters were added to the existing bond angle parameters in the a99SB-*disp* force field.

| [ angletypes ] | | | | | | |
|---|---|---|---|---|---|---|
| i | j | k | func | th0 | cth | |
| CT | SH | CT | 1 | 99.240 | 503.750 | ; Amber99Sb-disp |
| CT | OS | CA | 1 | 117.960 | 523.000 | ; Amber99Sb-disp |
| OS | CA | CA | 1 | 119.200 | 582.410 | ; Amber99Sb-disp |
| CA | CT | CA | 1 | 112.240 | 532.200 | ; Amber99Sb-disp |
| CA | CA | OS | 1 | 119.200 | 582.410 | ; Amber99Sb-disp |
| CA | OS | CT | 1 | 117.960 | 523.000 | ; Amber99Sb-disp |
| CT | CT | HO | 1 | 109.560 | 388.280 | ; Amber99Sb-disp |
| OH | CT | HO | 1 | 110.260 | 425.930 | ; Amber99Sb-disp |
| HO | CT | H1 | 1 | 108.460 | 328.030 | ; Amber99Sb-disp |
| CA | CA | Cl | 1 | 118.800 | 585.760 | ; Amber99Sb-disp |
| CT | CT | NT | 1 | 111.200 | 669.440 | ; Amber99Sb-disp |
| CT | NT | SO | 1 | 116.550 | 525.510 | ; Amber99Sb-disp |
| CT | NT | H | 1 | 109.500 | 418.400 | ; Amber99Sb-disp |
| NT | CT | H1 | 1 | 109.500 | 418.400 | ; Amber99Sb-disp |
| NT | SO | CT | 1 | 101.970 | 537.230 | ; Amber99Sb-disp |
| NT | SO | O | 1 | 107.430 | 595.800 | ; Amber99Sb-disp |
| SO | NT | H | 1 | 109.600 | 374.890 | ; Amber99Sb-disp |
| SO | CT | H1 | 1 | 107.150 | 361.500 | ; Amber99Sb-disp |
| CT | SO | O | 1 | 108.610 | 547.270 | ; Amber99Sb-disp |
| O | SO | O | 1 | 120.050 | 615.880 | ; Amber99Sb-disp |

**Supplementary Table 9: Added force field dihedral angle parameters for modified cysteine residues CYE2 and CYE7.** These parameters were added to the existing dihedral angle parameters in the a99SB-*disp* force field.

| [ dihedraltypes ] | | | | | | | | |
|---|---|---|---|---|---|---|---|---|
| i | j | k | l | func | phase | kd | pn | |
| CT | OS | CT | CA | 9 | 180.0 | 3.76560 | 0 | ; Amber99Sb-disp |
| CA | CA | OS | CT | 9 | 180.0 | 3.76560 | 0 | ; Amber99Sb-disp |
| OS | CA | CA | Cl | 9 | 180.0 | 15.16700 | 2 | ; Amber99Sb-disp |
| CA | CA | CA | Cl | 9 | 180.0 | 15.16700 | 2 | ; Amber99Sb-disp |
| Cl | CA | CA | HA | 9 | 180.0 | 15.16700 | 2 | ; Amber99Sb-disp |
| CT | CT | NT | SO | 9 | 0.00 | 1.25520 | 3 | ; Amber99Sb-disp |
| CT | CT | NT | H | 9 | 0.00 | 1.25520 | 3 | ; Amber99Sb-disp |
| CT | NT | SO | CT | 9 | 0.00 | 13.10987 | 2 | ; Amber99Sb-disp |
| CT | NT | SO | O | 9 | 0.00 | 13.10987 | 2 | ; Amber99Sb-disp |
| NT | SO | CT | H1 | 9 | 0.00 | 0.60436 | 3 | ; Amber99Sb-disp |
| SO | NT | CT | H1 | 9 | 0.00 | 1.25520 | 3 | ; Amber99Sb-disp |
| CT | SO | NT | H | 9 | 0.00 | 13.10987 | 2 | ; Amber99Sb-disp |
| H1 | CT | SO | O | 9 | 0.00 | 0.60436 | 3 | ; Amber99Sb-disp |
| O | SO | NT | H | 9 | 0.00 | 13.10987 | 2 | ; Amber99Sb-disp |
| CT | CT | CT | NT | 9 | 0.00 | 0.65084 | 3 | ; Amber99Sb-disp |
| O | CT | CT | NT | 9 | 0.00 | 0.65084 | 3 | ; Amber99Sb-disp |
| NT | CT | CT | H1 | 9 | 0.00 | 0.65084 | 3 | ; Amber99Sb-disp |
| H | NT | CT | H1 | 9 | 0.00 | 1.25520 | 3 | ; Amber99Sb-disp |
